# Elevated cerebrospinal fluid cytokine levels in tuberculous meningitis predict survival in response to dexamethasone

**DOI:** 10.1101/2020.11.23.394437

**Authors:** Laura Whitworth, Rajan Troll, Antonio J. Pagán, Francisco Roca, Paul H. Edelstein, Mark Troll, David Tobin, Nguyen Hoan Phu, Nguyen Duc Bang, Guy Thwaites, Nguyen Thuy Thuong Thuong, Roger Sewell, Lalita Ramakrishnan

**Affiliations:** Molecular Immunity Unit, Department of Medicine, University of Cambridge, and MRC Laboratory of Molecular Biology, Cambridge CB2 0QH, UK; Trinity College, Cambridge CB2 1TQ, UK; Department of Pathology and Laboratory Medicine, Perelman School of Medicine, University of Pennsylvania, Philadelphia, PA 19104, USA; Department of Molecular Genetics and Microbiology, and Department of Immunology, Duke University School of Medicine, Durham, North Carolina, USA; Oxford University Clinical Research Unit, Ho Chi Minh City, Vietnam; Hospital for Tropical Diseases, Ho Chi Minh City, Vietnam; Pham Ngoc Thach Hospital for Tuberculosis and Lung Disease, Ho Chi Minh City, Vietnam; Centre for Tropical Medicine and Global Health, Nuffield Department of Medicine, University of Oxford, Oxford, UK

## Abstract

Adjunctive treatment with anti-inflammatory corticosteroids like dexamethasone increases survival in tuberculosis meningitis. Dexamethasone responsiveness associates with a C/T variant in *Leukotriene A4 Hydrolase (LTA4H*), which regulates expression of the pro-inflammatory mediator leukotriene B4 (LTB4). TT homozygotes, with increased LTB4, have the highest survival when treated with dexamethasone and the lowest survival without. While the T allele is present in only a minority of the world’s population, corticosteroids confer modest survival benefit worldwide. Using Bayesian methods, we examined how pre-treatment levels of cerebrospinal fluid (CSF) pro-inflammatory cytokines affect survival in dexamethasone-treated tuberculous meningitis. *LTA4H* TT homozygosity was associated with global cytokine increases, including TNF. Association between higher cytokine levels and survival extended to non-TT patients, suggesting that other genetic variants may also induce dexamethasone-responsive pathological inflammation. These findings warrant studies that tailor dexamethasone therapy to pre-treatment CSF cytokine concentrations, while searching for additional genetic loci shaping the inflammatory milieu.

## INTRODUCTION

Tuberculous meningitis is the most lethal form of tuberculosis with a mortality of 25-40% in drug-sensitive HIV uninfected adults [1–3]. Drug resistant infection and HIV co-infection leads to even higher mortality [1, 3]. Because multiple investigations suggest that dysregulated inflammation plays a role in mortality from this disease, corticosteroids, which are broadly acting anti-inflammatory drugs, are now routinely used as adjunctive therapy to anti-tubercular antibiotics [4–6]. The relatively modest reduction of mortality with corticosteroids suggests that tuberculous meningitis may elicit different inflammatory responses, with corticosteroids helping those with high levels of inflammation. Genetic variation is likely to control these heterogeneous responses and a common functional variant in the *Leukotriene A4 Hydrolase* (*LTA4H*) gene is associated with responsiveness to dexamethasone, a potent corticosteroid [7–9]. LTA4H is a key enzyme in arachidonic acid metabolism that catalyzes the production of leukotriene B4, a pro-inflammatory lipid mediator with pleiotropic inflammatory effects [10, 11] . A C/T transition in the promoter modulates *LTA4H* gene and thereby protein expression. Consistent with its expression mediating an inflammatory milieu, CC and TT homozygotes have the lowest and highest *LTA4H* expression, respectively, with intermediate expression in CT heterozygotes. TT homozygotes have the greatest survival benefit from dexamethasone while suffering the highest mortality among those not given this drug [8].

The role of LTA4H in controlling inflammation and survival in the context of mycobacterial infections was first identified in a zebrafish forward genetic screen where animals with both low and high LTA4H expression were more susceptible to *Mycobacterium marinum* infection than their wild type counterparts [8, 12]. In the zebrafish, high LTA4H-mediated susceptibility is due to its increased product LTB_4_ inducing excessive Tumor Necrosis Factor (TNF) [8]. TNF causes pathogenic programmed necrosis of macrophages in *Mycobacterium marinum-*infected zebrafish as well as *Mycobacterium tuberculosis-*infected human macrophages [13, 14].

Because tuberculous meningitis is characterized by a necrotizing granulomatous reaction and macrophage-rich meningeal exudates [15, 16], we wanted to determine if the *LTA4H* TT genotype mediates increases in cerebrospinal fluid (CSF) cytokines and if these increases are associated with dexamethasone responsiveness. Consistent with the findings of Tobin et al. [8], analysis of a second Vietnam tuberculous meningitis cohort, where all individuals had been treated with dexamethasone, showed that *LTA4H* TT HIV-uninfected individuals had increased survival over their non-TT counterparts [7, 9]. The same study also determined CSF cytokine levels to test the prediction that TT individuals have a hyper-inflammatory CSF profile reflected by elevated cytokines [7]. The original analysis of the cytokine profiles in this study was conducted using linear trend tests and found that the median levels of all 10 assayed cytokines were increased in TT compared to CC and CT patients, but in a multiple comparison test only the interleukins IL-1β, IL-2, and IL-6 increases achieved statistical significance [7].

Here, we have re-analyzed the cytokine data from HIV-uninfected adults with tuberculous meningitis using Bayesian methods. Bayesian analysis can detect significant results and relationships not detected by frequentist methods because they do not impose a penalty for multiple comparisons and can effectively detect significant differences that are hidden by Type 2 errors in frequentist analysis [9]. Moreover, we don’t know the class of distribution (e.g. Normal, Gamma, etc) from which the cytokine values come. Bayesian methods can identify the likely class of distribution and take this information into account for the analyses even without having surety about the correct class.

Using Bayesian methods (detailed in Appendix 1 and Appendix 2), we find that survival in response to dexamethasone is associated with significant increases in all cytokines tested before or at the start of treatment, representing innate pro-inflammatory, helper T cell-associated and immunomodulatory classes. While the *LTA4H* TT genotype was associated with increases in these cytokines, we also found that increased cytokines are associated with survival in an *LTA4H*-independent manner in this dexamethasone-treated cohort.

## RESULTS

### In tuberculous meningitis patients, LTA4H TT genotype is associated with increased CSF levels of multiple cytokines, including TNF

Analysis of the CSF cytokine values from Thuong et al. [7] (Table 1) showed that none followed a normal (Gaussian) distribution. For nearly all values, log-skew-Student (log-noncentral-t distribution) was the preferred distribution class both in the dataset as a whole and for the various subsets considered in our analyses (Appendix 2). Comparisons were performed using restricted geometric means as is appropriate for such heavy-tailed approximately logarithmically distributed data (Box 1 and Appendix 2). Furthermore, unlike the previous analysis, we made no assumption that there would be a linear trend with the number of T-alleles in a given patient. Using this method of analysis we found that TT patients had significant increases in all measured CSF cytokines, except Interferon-γ(IFNγ) and IL-4, compared to both CC and CT patients who had similar levels to one another (Figure 1A and Supplementary Table 1). Similarly, a comparison of cytokine levels in TT patients to those in combined non-TT (CT and CC) patients showed that these levels in TT patients were significantly higher for all cytokines except for IFN-γ and IL-4 (Figure 1B). The finding that a single T allele does not have a discernible influence on inflammatory pathways is consistent with the CC and CT patients in this cohort having similarly lower survival than TT patients when all patients were receiving dexamethasone therapy [7, 9]. Thus, TT homozygosity is associated with increased cytokine concentrations across the board, including cytokines that are associated with an acute inflammatory response (TNF, IL-1β, and IL-6); with T cell activation and regulatory T cell homeostasis (IL-2), innate and adaptive type-1 immunity (IL-12 and IFNγ); innate and adaptive type-2 immunity (IL-4, IL-5, and IL-13), and immune modulation (IL-10). Importantly, TNF, which drives the pathogenesis caused by LTA4H excess in the zebrafish model of TB [8, 13, 14] is significantly increased in TT patients.

**Table 1.** Patient cohort characteristics.

|  | **Thuong 2017 cohort**<br>**(cytokines measured)** | **Tobin 2012** |
| --- | --- | --- |
| **Total n** | 306 | 179 |
| **Age in years**<br>***median (range)*** | 39 (18-87) | 34 (15-83) |
| **BMRC TBM grade**<br>***no. (% of total)*** |  |  |
| 1 | 109 (35.6) | 43 (24.0) |
| 2 | 141 (46.1) | 84 (46.9) |
| 3 | 56 (18.3) | 52 (29.1) |
| **Overall mortality**<br>***no. (%)*** | 57(18.6) | 38 (21.2) |
| **LTA4H rs17525495**<br>**genotype**<br>***no. (% of total)*** |  |  |
| CC | 148 (48.4) | 84 (46.9) |
| CT | 142 (46.4) | 73 (40.8) |
| TT | 16 (5.2) | 22 (12.3) |

**Figure 1.**
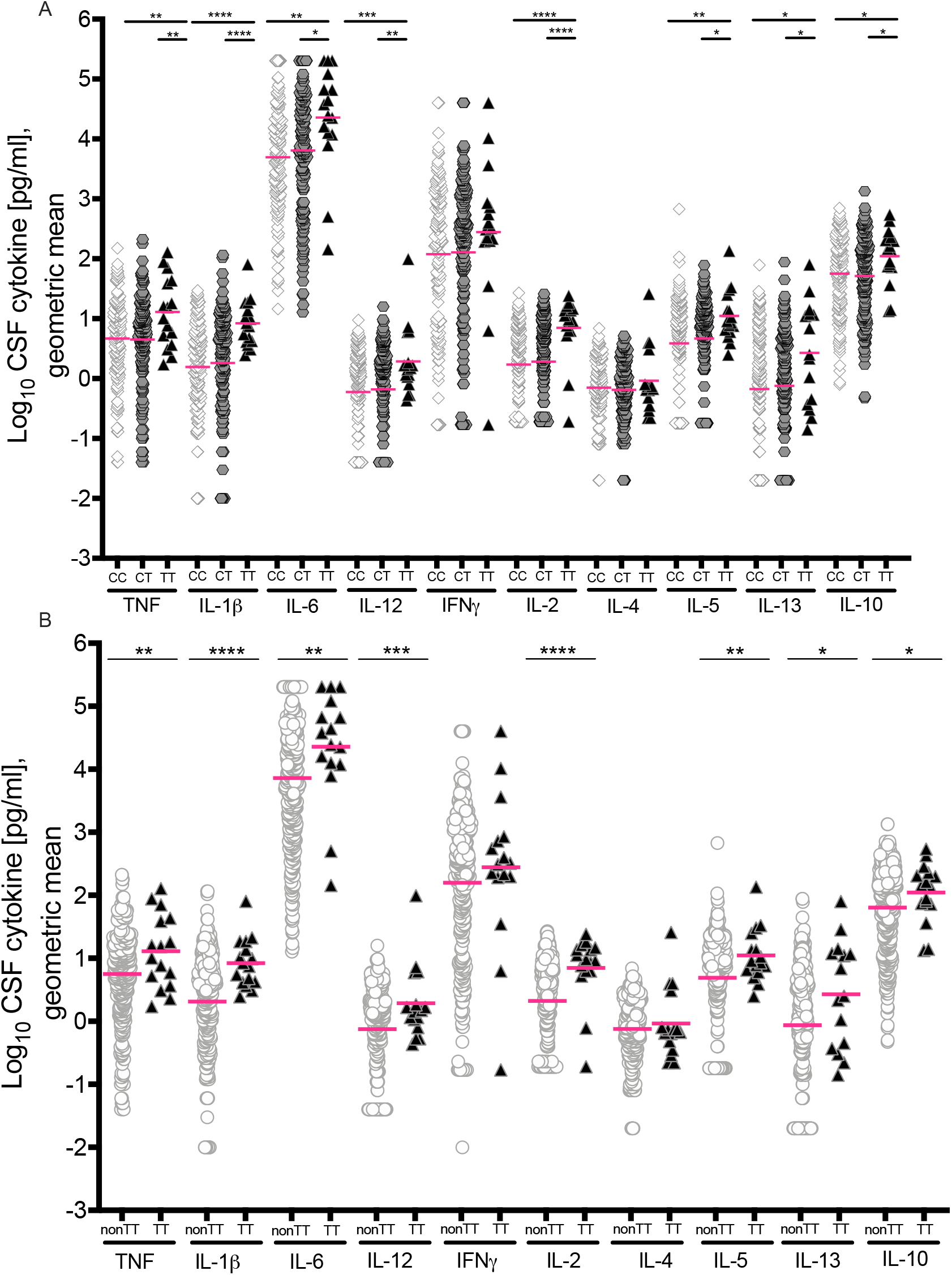
Cerebrospinal fluid (CSF) cytokine levels in HIV-negative patients grouped by *LTA4H* genotype. (A) Cytokine levels in CSF from CC (n=148), CT (n=142), and TT (n=16) patients. (B) Cytokine levels compared between non-TT (n=290) and TT (n=16) genotypes. Magenta lines indicate geometric means. Asterisks indicate probability that right-hand group values are significantly greater than the left (* ≥ 0.95, ** ≥ 0.99, *** ≥ 0.999, **** ≥ 0.9999). Unspecified comparisons are not significant.

### The LTA4H TT genotype exerts a compensatory regulation on CSF cytokine levels in more severe disease

Tuberculous meningitis patients can present with a wide-ranging disease severity, reflected by the presence or absence of focal neurological signs, or a generalized decrease in responsiveness including coma [17]. The modified British Medical Research Council tuberculous meningitis grading system categorizes patients into three grades in increasing order of severity [3, 17]. Prior analysis of this cohort found that a disease grade was associated with a trend to increased cytokines across the board with a significant increase for only one, IFNγ [7]. Our analysis found all to be increased with increasing grade, with significant increases between grades for seven of the ten (Figure 2A and Supplementary Table 1). Since there were increased cytokine levels for both the TT genotype and for higher disease grades, we predicted that these levels would be highest in TT patients in the higher disease grades. Whereas non-TT patients had a similar pattern of increased cytokines with increasing disease grade as the overall cohort (Figure 2B), we were surprised to find that in TT patients, the pattern was reversed. The majority of the cytokines were lower in Grades 2 and 3 than in Grade 1, significantly so in many cases (Figure 2C). The major shift occurred between Grades 1 and 2. Grade 3 cytokines were not lower than Grade 2; the levels were either similar in these two grades or Grade 2 levels were non-significantly lower. These findings suggest the existence of compensatory mechanisms in TT patients that limit extreme increases in cytokine levels driven by increased disease severity. Consistent with this hypothesis, when we compared cytokine levels in non-TT to those in TT patients stratified by disease grade, cytokine levels in TT patients were higher in all grades, with increases that were the greatest and most significant in Grade 1, rather than in Grades 2 and 3 (Figure 2D-F).

**Figure 2.**
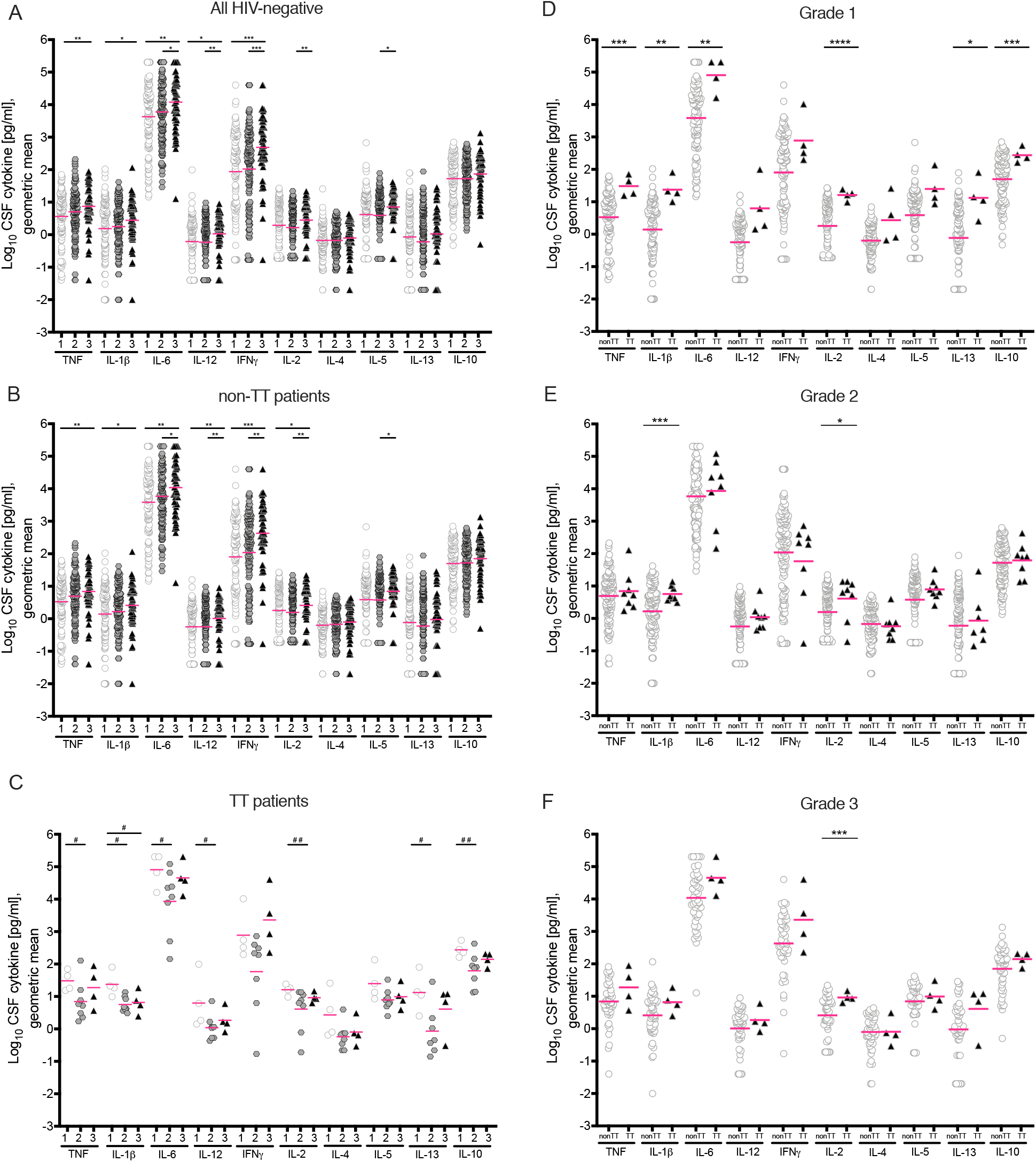
Cytokine levels in HIV-negative patients by grade and by genotype. (A) All patients (Grade 1 n=109, Grade 2 n=141, Grade 3 n=55); (B) Non-TT patients (Grade1 n=105, Grade 2 n=133, Grade 3 n=51), and (C) TT patients (Grade 1 n=4, Grade 2 n=8, Grade 3 n=4). (D) Grade 1 patients; (E) Grade 2 patients; (F) Grade 3 patients. Magenta lines indicate geometric means. Asterisks indicate probability that right-hand group values are significantly greater than the left (* ≥ 0.95, ** ≥ 0.99, *** ≥ 0.999, **** ≥ 0.9999). Hash symbols indicate probability that left-hand group values are significantly greater than the right (# ≥0.95, # # ≥0.99). Unspecified comparisons are not significant.

### Both LTA4H TT-dependent and -independent CSF cytokine increases are associated with survival in response to dexamethasone

Thuong et al. [7] compared cytokine levels independent of *LTA4H* genotype in tuberculous meningitis survivors to non-survivors following adjunctive dexamethasone treatment and found that survivors had increased cytokine levels. Our re-analysis confirmed this result - all cytokines were significantly increased in survivors compared to those who died (Figure 3A). Because TT patients had significantly increased survival with dexamethasone as compared to non-TT patients [7, 9], we hypothesized that the increased cytokine levels in survivors overall would be restricted to TT patients. However, even among non-TT patients only, survivors had significantly increased cytokines across the board when compared to those who died (Figure 3B). We could not compare TT survivors to non-survivors as the four TT patients who died did not have CSF cytokine measurements. Comparison of TT survivors to non-TT survivors revealed that most cytokines were significantly higher in TT survivors than in the non-TT survivors (Figure 3B). CC and CT survivors each also had higher cytokines than non-survivors overall and in all three disease grades (Figure 3-figure supplement 1). Together these analyses show that the default inflammatory response to tuberculous meningitis includes global increases in CSF cytokines, and suggest they are associated with a survival benefit from dexamethasone. Overlaid on these are further increases mediated by the *LTA4H* TT genotype.

**Figure 3.**
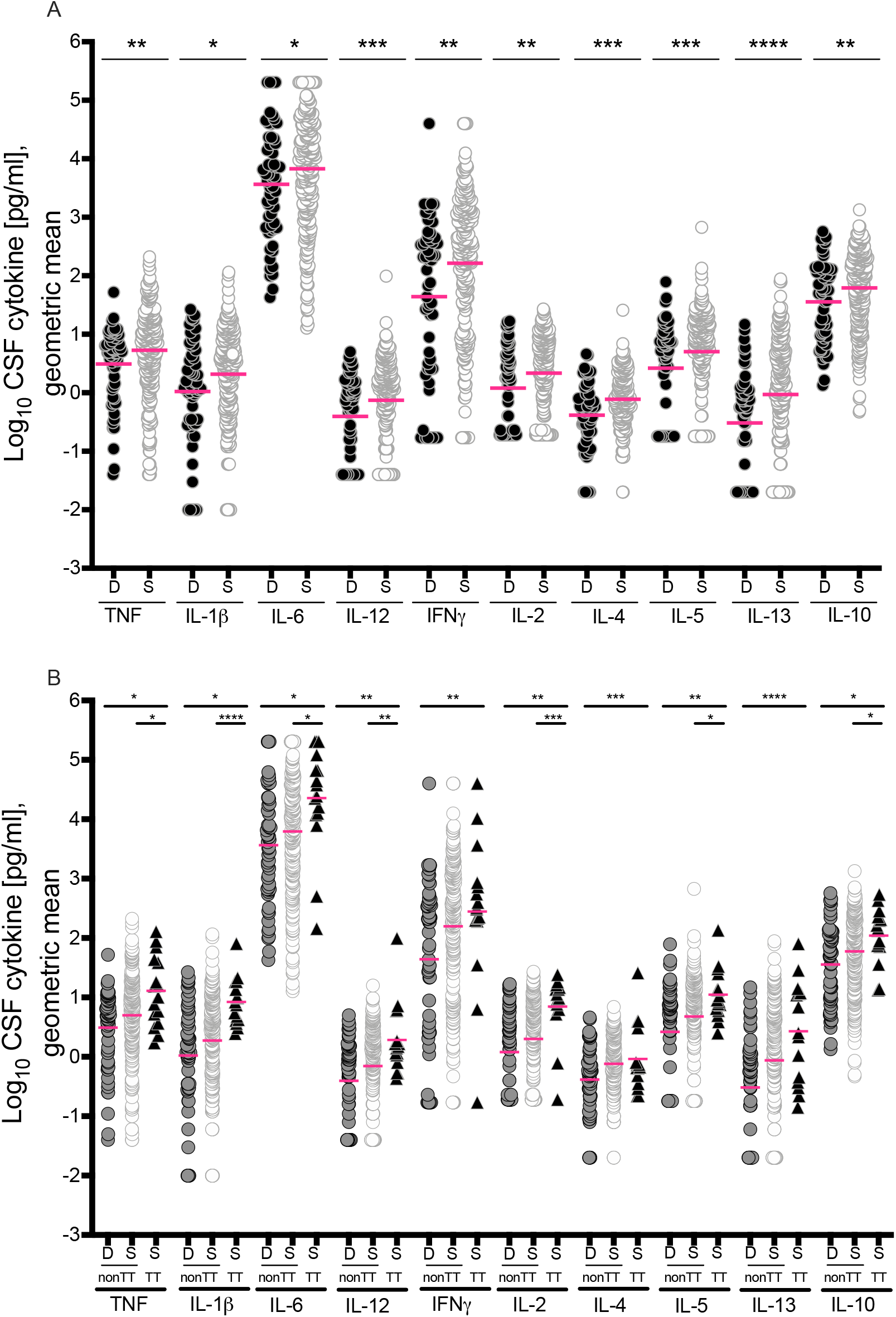
Cytokine levels in HIV-negative survivors and non-survivors. (A) Cytokine levels in patients who survived (S, n=248) versus those who died (D, n=57); (B) separated by genotypes into non-TT (deaths, n=57, survivors, n=232) and TT (all survived, n=16). Magenta lines indicate geometric means. Asterisks indicate probability that right-hand group values are significantly greater than the left (* ≥ 0.95, ** ≥ 0.99, *** ≥ 0.999, **** ≥ 0.9999). Comparisons performed for each cytokine: non-TT dead vs survived and non-TT survived vs TT survived. Cytokine levels between non-TT dead and TT survived were not compared. See also Figure 3 - figure supplement 1.

The finding that the *LTA4H* TT patients have increased CSF cytokines over their non-TT counterparts provides an explanation for why they survive better when treated with dexamethasone than their non-TT counterparts [8]. However, we had now shown in this study that among dexamethasone-treated tuberculous meningitis patients, *LTA4H* TT-independent cytokine increases are also associated with survival, raising the question of whether non-TT patients might also benefit from dexamethasone. By the time the cytokine analysis study was undertaken, adjunctive dexamethasone had become standard-of-care treatment so that all patients were given this drug [7]. Therefore, to answer the question, we re-analyzed the survival data from the Tobin et al. study (Table 1), using recently-described Bayesian methods, which had compared survival of patients of the three *LTA4H* genotypes with and without adjunctive dexamethasone [8, 9].

The survival of dexamethasone-treated CT heterozygotes was different between the two studies, appearing more similar to that of the TT patients in the Tobin et al. study but more similar to that of the CC patients in the Thuong et al. study [7, 8]. So we re-analyzed both studies using Bayesian methods, separating the non-TT patients into the individual CC and CT genotypes. In the Tobin study, in the absence of dexamethasone treatment, TT patients had worse survival than both CC and CT patients with the difference being just short of being significant (maximum posterior probability 0.946) (Figure 4A). There was no significant difference between CC and CT patients (Figure 4A). Among dexamethasone treated patients, TT survival was significantly higher than CC survival (Figure 4B). CT survival was in between the two, significantly higher than CC and non-significantly lower than TT (Figure 4B). In the Thuong study, CT survival was significantly worse than TT and not significantly different from CC (Figure 4C). We confirmed this shift in CT survival between the two studies by a direct comparison of the Tobin and Thuong studies. CT survival was significantly worse in the Thuong study whereas CC and TT survival were not significantly different in the two studies (Figure 4 D-F).

**Figure 4.**
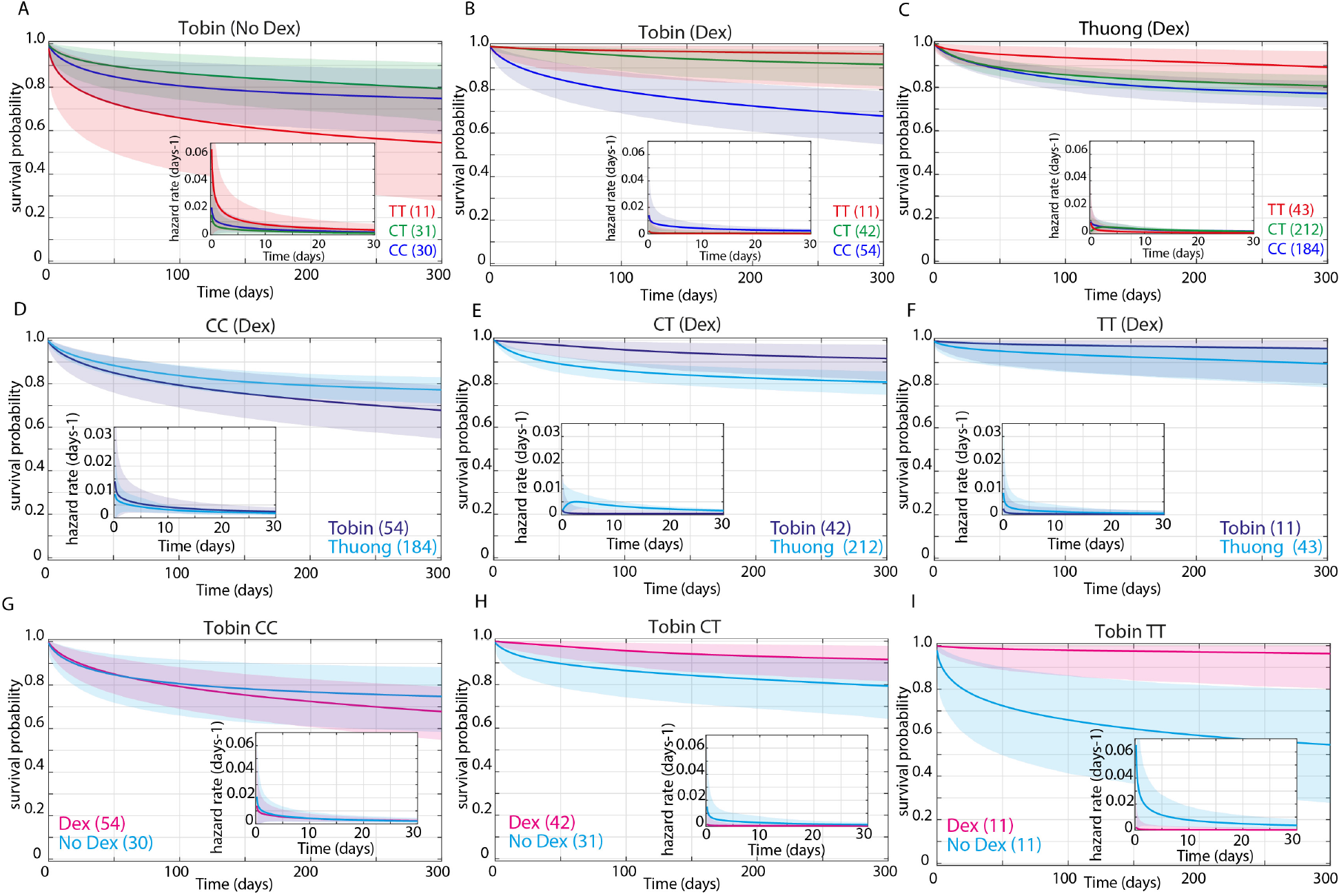
Effect of dexamethasone and *LTA 4 H* rs 17525495 genotype on survival probability of HIV-negative patients from Tobin 2012 and Thuong 2017. Mean posterior survival probability curves; inset plots represent mean posterior hazard rates for the first 30 days. Shaded areas represent the 95% Bayesian confidence limits for posterior probability. (A) In Tobin 2012 (No Dex), TT survival was non-significantly reduced compared to non-TT (maximum probability 0.946). (B) In Tobin 2012 (Dex), TT survival was significantly greater than non-TT from day 40 onwards (maximum probability 0.976, survival gap 17%). Probability that TT hazard rate (inset) is lower than non-TT is >0.95 from day 2 to day 252 (maximum probability 0.972, peak ratio at day 97). CT survival was significantly greater than CC from day 3 onwards (maximum probability 0.999, survival gap 23%), and CT hazard rate significantly lower than CC from day 1 onwards (maximum probability 0.996, ratio peaks at 12 on day 3 and remains >3 throughout). (C) In Thuong 2017 patients (Dex), CC and CT survival comparisons do not differ significantly (maximum probability 0.91). TT survival was significantly greater than CC from day 42 onwards (maximum probability 0.987, survival gap 12%). Probability that TT hazard rate is lower than CC is >0.95 days 15-138 (maximum probability 0.991, ratio peaks at 3.4 on day 62 and remains >1 until day 250). TT survival was also significantly greater than CT from day 53 to day 254 (maximum probability 0.964, survival gap 9%). Probability that TT hazard rate is lower than CT is >0.95 from day 7 to day 73 (maximum probability 0.979, ratio peaks at 2.9 on day 22 and remains >1 to day 234). (D) In CC (+Dex) patients, survival was non-significantly greater in the Thuong cohort (maximum probability 0.939, survival gap 9%). (E) In CT (+Dex) patients, survival was significantly greater in the Tobin cohort from day 5 onwards (maximum probability 0.993, survival gap 11%). Tobin CT (+Dex) hazard rate was significantly lower than Thuong CT from day 2 to day 45 (maximum probability 0.997, peak ratio 9.8 on day 4). (F) In TT patients (+Dex), survival was non-significantly greater in the Tobin cohort (maximum probability 0.90, survival gap 6%). (G) Tobin CC patient survival did not differ significantly with and without Dex treatment (maximum probability 0.80). (H) Tobin CT patient survival was significantly greater with Dex from day 7 to day 88 (maximum probability 0.964, survival gap 11%). CT (+Dex) hazard rate was significantly lower than CT (No Dex) from day 3 to day 18 (maximum probability 0.967, peak ratio 9 on day 2 and remains >1 throughout). (I) Tobin TT patient survival was significantly greater with Dex from day 1 onwards (maximum probability 0.997, survival gap 41%). TT (+Dex) hazard rate was significantly lower from day 1 onwards (maximum probability 0.996, peak ratio 35 on day 2). See also Figure 4 -figure supplement 2.

Finally, we asked whether and how dexamethasone influenced the survival of each genotype in the Tobin study. Directly comparing survival of each of the three genotypes with and without dexamethasone, we found that TT patients derived the greatest benefit from dexamethasone, CT patients had a smaller but still significant benefit, and CC patients were neither helped nor harmed by dexamethasone (Figure 4G-I).

In sum, because the CC and TT patients survived similarly in response to dexamethasone in the Thuong and Tobin studies, we can use the survival with and without dexamethasone in the Tobin study together with the pre-treatment cytokine levels in the Thuong study to draw the following two conclusions: (1) TT patients benefit very substantially from dexamethasone, consistent with their higher pre-treatment cytokine levels; and (2) among dexamethasone-treated patients, CC survivors have higher pre-treatment cytokine levels than non-survivors. These two findings can be reconciled by a model (Figure 5) where dexamethasone reduces the higher pre-treatment cytokine levels in TT patients to a level optimal for survival, whereas the lower cytokine pre-treatment levels in many of the CC patients are lowered further by this treatment to suboptimal levels, so that there is no apparent benefit of the drug to the cohort overall.

**Figure 5.**
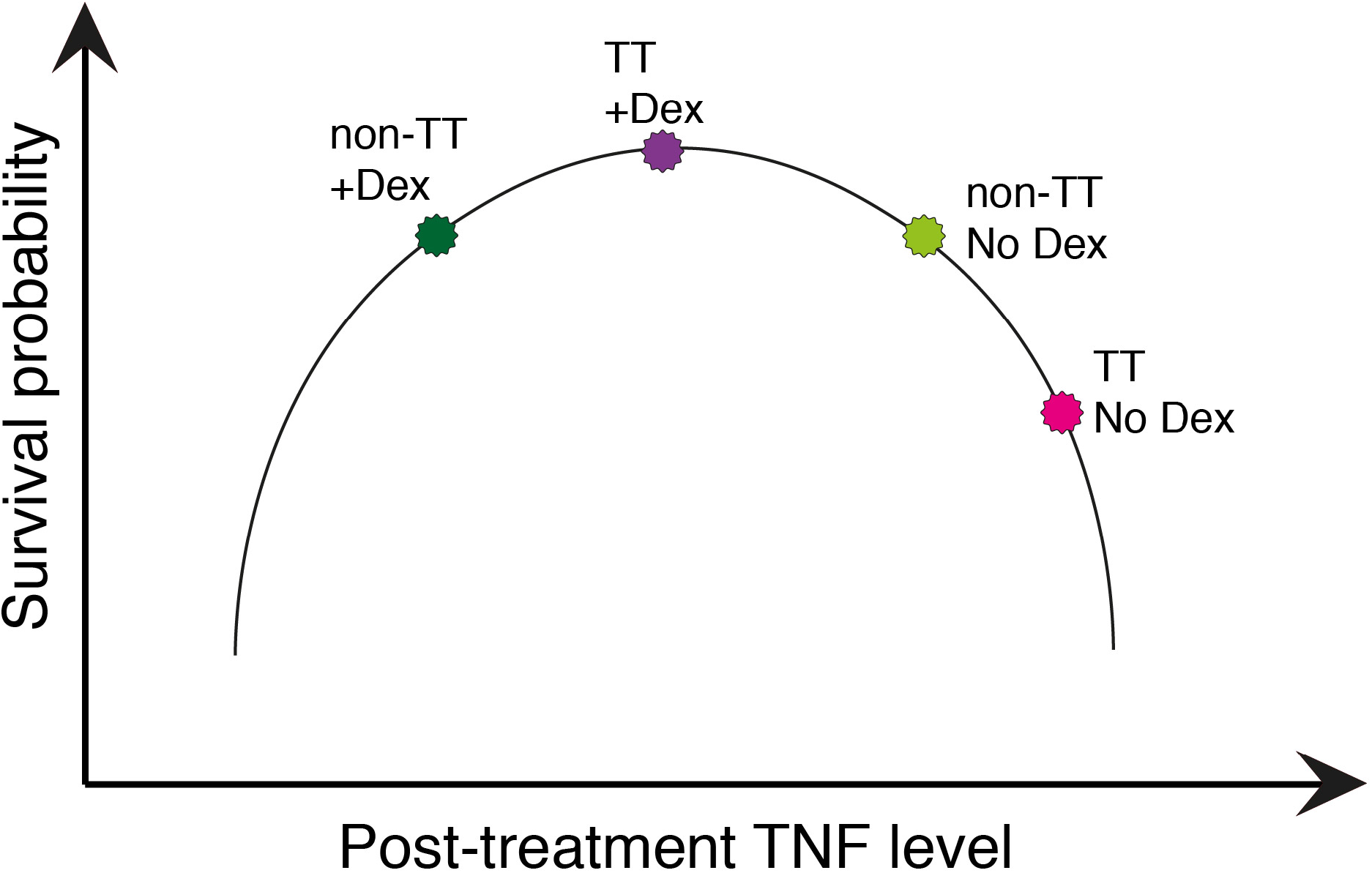
Proposed model of dexamethasone-mediated effects on survival probability interacting with *LTA4H* rs17525495 genotypes.

## DISCUSSION

Dysregulated intracerebral inflammation has long been thought to be responsible for the high mortality and morbidity of tuberculous meningitis. Multiple cytokines can be major effectors of dysregulated immune responses, yet pre-treatment cytokine data from TBM patients are limited [18, 19]. Therefore, the Thuong et al. study [7] where CSF cytokines were collected in 306 HIV-uninfected patients, all of whom were treated with dexamethasone, together with comprehensive clinical information and survival analyses, provided an unprecedented opportunity. This cohort allowed for an analysis of CSF cytokine concentrations in TBM with respect to disease severity on presentation and outcome following dexamethasone treatment. Moreover, this study confirmed the survival benefit of the *LTA4H* TT genotype, providing the opportunity to ask if this hyperinflammatory genotype was associated with increased cytokines. Comparison of CSF cytokines applying frequentist statistical methods (linear trends tests) to median cytokine values showed that increased pre-treatment levels of most (8 of the 10 tested) were significantly associated with survival with dexamethasone, indicating that these higher levels are pathogenic. All cytokines were increased with increased disease grade, but only one of these, IFNγ, was significantly increased in those with Grade 3 disease. The *LTA4H* TT genotype was also associated with global increases in cytokine concentrations across the board in comparison to the non-TT genotype patients, but the differences were significant in only 3 of the 10 cytokines. Given that most cytokines are induced by interrelated and often shared signal transduction networks, notably the NFκB family of transcription factors [20], these patchy statistically significant differences were more likely to represent a Type 2 statistical error than biologically relevant patterns. So we turned to Bayesian methods to re-analyze these data and looked for associations between pre-treatment cytokine levels with disease severity, survival and *LTA4H* genotype, not only singly, but also in combination.

Bayesian analysis shows that all cytokines tested, representing multiple functional classes - innate pro-inflammatory, Th1- and Th2-associated and immunomodulatory - are associated with survival in this dexamethasone-treated cohort. This global induction of cytokines is consistent with the induced inflammatory trigger mediating effects upstream of a common signaling axis for all of them without significant additional downstream regulation. The finding that *LTA4H* TT genotype further increases all ten cytokines is consistent with prior work showing that LTB_4_ binding to its receptors activates the NFκB pathway [21]. Furthermore, TNF, which has been implicated in the pathogenesis of tuberculosis, including tuberculous meningitis [13, 14, 22], appears to be dysfunctionally increased both in an *LTA4H*-independent and -dependent manner. Finally, this work highlights the role of the previously described regulatory circuits that dampen *LTA4H* TT - mediated inflammation (i.e., leukotriene B_4_) in the context of one of the most lethal infectious diseases of humans [11, 23].

### Association of increased CSF cytokines and survival even independent of LTA4H genotype

Our initial goal in performing these analyses was to ask whether the *LTA4H* TT genotype is associated with global increases in pre-treatment CSF cytokines, as would be predicted by its activation of the NFκB pathway [21]. *LTA4H* TT individuals with tuberculous meningitis have a striking survival benefit from dexamethasone, which causes a global reduction in cytokines, and we find that the TT genotype is indeed associated with higher pre-treatment CSF cytokines across the board. However, even in Europe and Africa where the *LTA4H* T allele is much rarer (10% frequency) than in Asia (up to 33%; from https://tinyurl.com/y4c232e3) [8, 9, 24], multiple small studies have a modest survival benefit from corticosteroids comparable to that seen in Asia [4, 5, 25, 26]. In fact, the earliest studies suggesting corticosteroid benefit, that gave the impetus for the larger randomized controlled trial in Vietnam [6], were done in patient populations that were mostly of European descent, in which the LTA4H TT homozygote genotype frequency would have been rare (1-4% of population) [24–26]. Perhaps our most important finding is that even among *LTA4H* non-TT individuals, higher CSF cytokines are associated with higher survival in response to dexamethasone. However, when we analyze a prior cohort that enables comparison of survival with and without dexamethasone (but lacks CSF cytokine analysis), we find that while TT patients gain a major survival advantage from dexamethasone, non-TT patients’ survival is neither helped nor harmed by it. These two findings can be reconciled in two ways. The first is predicated on the idea that a major mechanism of dexamethasone’s survival benefit is through its cytokine-reducing effect [27, 28]. Non-TT pre-treatment CSF cytokines are lower on average than TT, and the currently-used dexamethasone dosages may decrease their cytokine levels to suboptimal levels (Figure 5). In optimal concentrations, many of these cytokines play a role in the host responses that eliminate bacteria, and this function may be lost if they are lowered below a threshold. The model that dexamethasone optimizes TT cytokine levels while reducing non-TT levels too much can be tested when the results of an ongoing randomized clinical trial of dexamethasone for non-TT patients become available (trial registration:NCT03100786) [29]. Non-TT survivors in the control arm would not be expected to have higher cytokine levels than non-survivors. Our findings could pave the way for stratification of patients for dexamethasone therapy both by *LTA4H* genotype and by CSF cytokine levels, potentially indicating higher and lower doses of the drug or no drug at all. Because corticosteroids exhibit a dose-dependent degree of immunosuppression of cytokines, including TNF [30], it is possible that the lower non-TT grades may benefit from lower doses of these drugs.

Alternatively, it is possible that the much greater benefit of dexamethasone on TT survival derives from its countering other inflammatory pathways activated by LTB_4_, including neutrophil chemotaxis and degranulation, and production of reactive oxygen and nitrogen species [10], which would also be inhibited by corticosteroids [27, 28].

### Regulatory networks may keep LTA4H TT cytokine increases in check

The finding that disease severity increases cytokine levels in non-TT but not in TT patients suggests *LTA4H* TT-specific compensatory mechanisms that appear to dampen disease grade mediated cytokine increases. LTB_4_ mediates its activity through two receptors BLT1 and BLT2 [31]. BLT1, the high affinity receptor is associated with pro-inflammatory responses, and is itself down regulated by increased inflammatory determinants, including TNF [11, 23]. One could imagine a scenario where the interplay between disease severity and TT driven cytokine increases are sufficiently high so as to down regulate BLT1, which would halt LTB_4_-BLT1 mediated cytokine increases. Moreover, BLT1 down regulation would promote LTB_4_ interactions with its BLT2 receptor which, having a ~ 50-fold lower affinity [31], would ordinarily not be in play. LTB_4_-BLT2 interactions can promote both pro- and anti-inflammatory responses in different scenarios [11, 23, 31, 32]. Germane to this study, LTB_4_-BLT2 interactions are reported to downregulate macrophage activation and all four cytokines tested - TNF, IL-1β and IL-6 and IFNγ - in a mouse inflammatory colitis model [32]. Even if the interaction results in pro-inflammatory responses, the substantial reduction in binding affinity would be expected to result in reduced downstream effects.

### LTA4H TT-dependent and LTA4H-independent TNF dysregulation

Our finding that dexamethasone is associated with increased survival in both LTA4H TT and non-TT patients is tantalizing from a therapeutic standpoint. In the zebrafish, high LTA4H increases disease severity through increasing TNF, which in excess causes increased disease pathogenesis through a newly-identified programmed macrophage necrosis [13, 14]. We were particularly interested in this question because in the zebrafish, several pathway-specific drugs that inhibit macrophage necrosis without being broadly anti-inflammatory have been identified, all of which are have a decades-long history of use in humans for other conditions [13, 14]. Therefore, unlike glucocorticoids, these drugs would be beneficial to those with excessive TNF while being neutral to the other patients, as they target the downstream effects of excess TNF, without broadly reducing overall cytokines to levels which are detrimental to survival.

### Conclusions and Implications for future studies

The use of Bayesian methods has enabled important insights into the induction and regulation in tuberculous meningitis and the possible detrimental effects of their dysregulation. On-going randomized control trials that will enroll >1200 participants are examining the role of adjunctive dexamethasone adults with tuberculous meningitis, stratifying participants according to HIV infection and *LTA4H* genotype [29, 33]. Pre- and post-treatment CSF cytokine analysis will be performed in all participants [29, 33]. These trials will allow for validation of the analyses presented here as well as test the models and hypotheses that have arisen from them. Finally, some (though not all) studies done over decades across the globe have found adjunctive corticosteroid treatment to have a modest early benefit in the contagious and most common form of tuberculosis, that involving the lung, both in reducing inflammation and bacterial burdens [34–37]. Spatial studies of human tuberculous granulomas combining mass spectrometry find that necrotic tuberculous granulomas are enriched for LTA4H and TNF as compared to non-necrotic granulomas from the same lung, making it plausible that corticosteroids exert their effects through these determinants [38]. The analytical methods developed here could be readily tailored to examine the role of corticosteroids as drugs that may improve outcome in pulmonary TB as well.

## MATERIALS AND METHODS

The anonymized tuberculous meningitis patient cohort CSF cytokine level data used here has been previously described [7, 9] (Table 1). The tuberculous meningitis cohort used for survival analysis has also been previously described [6, 8] (Table 1). Patients were admitted to one of two tertiary care referral hospitals in Ho Chi Minh City, Vietnam: Pham Ngoc Thach Hospital for Tuberculosis and Lung Disease (Hospital 1), or the Hospital for Tropical Diseases (Hospital 2). Patients were grouped on study entry according to the modified British Medical Research Council tuberculous meningitis grade: patients with a Glasgow Coma Scale (GCS) score of 15, with no focal neurologic signs, were designated Grade 1; patients with GCS score of either 11-14, or 15 with focal neurologic signs, were designated Grade 2; and patients with GCS scores of 10 or less were designated as Grade 3 [6]. The Tobin et al., 2012 [8] study had a higher proportion of patients with more severe disease (Table 1). Patients with cytokine measurements were classified as having definite, probable, or possible tuberculous meningitis in accordance with published diagnostic criteria [39]. Patients in the Tobin et al., 2012 study all had definite tuberculous meningitis [8]. *LTA4H* rs17525495 genotypes were determined by Taqman assay [7, 8]. The cohort used for survival analysis comprises patients from a trial where patients were randomized to get either dexamethasone for the first 6-8 weeks or placebo [6, 8]. Patients with moderate to severe disease (Grades 2 and 3) were given intravenous (IV) treatment for 4 weeks (0.4mg/kg/day week 1, 0.3mg/kg/day week 2, 0.2mg/kg/day week 3, and 0.1mg/kg/day week 4), followed by oral administration for 4 weeks, starting at 4mg/day and decreasing by 1mg each week. Patients with mild disease (Grade 1) received 2 weeks of IV treatment (0.3mg/kg/day week 1, 0.2mg/kg/day week 2), followed by oral therapy for 4 weeks, starting at 0.1mg/kg/day in week 3, then a total of 3mg/day in week 4, decreasing by 1 mg/week.

All patients in the cohort used for cytokine analysis were treated with adjunctive dexamethasone as above and CSF specimens collected on enrollment from 306 HIV-negative and 219 HIV-positive patients [7]. CSF for cytokine measurements was available in a smaller proportion of patients admitted to Hospital 1 than Hospital 2 (55.9% vs. 96.6%) (Supplementary Table 2). Further analysis showed that Hospital 1 TT patients were significantly less likely to have had cytokine measurements than non-TT patients (Supplementary Table 3). There was no such bias in relation to survival status or disease grade severity (Supplementary Table 3). To determine if the Hospital 1 collection bias in the TT vs. non-TT patients changed the analysis, all data were analyzed for the whole cohort and Hospital 2 alone (Supplementary Table 1). 349 of the 350 comparisons yielded similar patterns in the two datasets with one exception, with differences being in the same direction, with some loss of significance when considering Hospital 2 alone, due to the reduction in the number of patients. The only comparison where there was a change in the direction was IL-5, Grade 1 versus Grade 2, which was significantly greater in Grade 2 than in Grade 1 in Hospital 2 alone and nonsignificantly greater in Grade 1 than Grade 2 in the overall cohort. The analysis for the overall cohort are presented in Figures 1–3.

As described in Thuong et al 2017 [7], 10 cytokines - TNF, IL-1β, IL-6, IL-12, IFNγ, IL-2, I-L4, IL-5, IL-13, and IL-10 - were measured in the stored CSF samples by Luminex multiplex bead array analysis. The cytokine levels reported in the previous study were re-analyzed using Bayesian methods comparing restricted geometric mean values from groups of patients separated by HIV-status, survival status (“survivors” are patients who were either known to have survived, or, if lost to follow-up, were censored at the time of last recorded outcome), BMRC tuberculous meningitis grade, and *LTA4H* rs17525495 genotype (Appendix 2). A number of CSF samples yielded cytokine concentrations that were outside the linear ranges of the assays and had been assigned specified high or low fixed values. To handle apparent limits to the measurable value, resulting in many observations of the maximum and minimum seen value, these maximal and minimal values were spread in a process called dithering. (Appendix 2). The geometric means of both the undithered and dithered data are in Supplementary Table 1. The geometric means indicated in Figures 1–3 are based on the undithered data. Significance analyses for all comparisons were performed on both dithered and undithered data (Supplementary Table 1) and the results from the analyses of the dithered data are presented in Figures 1–3. In 869 of the 890 comparisons, the significance for the majority of comparisons did not change between analysis of undithered and dithered data. Of the 21 comparisons which were changed by dithering, significance was lost upon dithering in 15 and gained upon dithering in 6. The instances where significance changed upon dithering are color-coded blue and marked with a minus sign in the less significant cell in Supplementary Table 1.

The Bayesian methods used for the survival analysis in Figure 4 are detailed in Whitworth et al.[9].

## Supporting information

Software

### Box 1 Definitions and usages

#### Definitions

- *Posterior probability* – the probability after seeing the data
- *Survivors* (as opposed to nonsurvivors) – those who were not known to have died during the period that they were observed, i.e. those who either were censored or died.
- *Restricted geometric mean* – the geometric mean of a distribution, restricted to an interval excluding only extreme tail events; see Appendix 2 for reasons for using this concept
- *Dithering* – A process that spreads out data points concentrated on the maximum and minimum seen values, as these likely come from a limit in the measurement procedure

#### Abbreviated and example for Figures 1–3 and associated text

*“A was significantly greater than B”* - the posterior probability given the dithered data that the restricted geometric mean of the variable A in the relevant population was greater than that of the variable B was at least 0.95 (and similarly for “*significant increase*”, etc.)

*“A was not significantly different from B”* - the posterior probability given the dithered data that the restricted geometric mean of the variable A in the relevant population was greater than that of the variable B was between 0.05 and 0.95 (and similarly for “*no significant difference*”, etc.)

“*A was greater than B*” - the (restricted) geometric mean of the undithered data in A was greater that of B (note that the presence or absence of “restricted” makes no difference in this definition as all the observed data is within the range of restriction)

“*The probability that A was greater than B was p*” - the posterior probability given the dithered data that the restricted geometric mean of the variable A in the relevant population was greater than that of the variable B was p

**Figure 3 - figure supplement 1.**
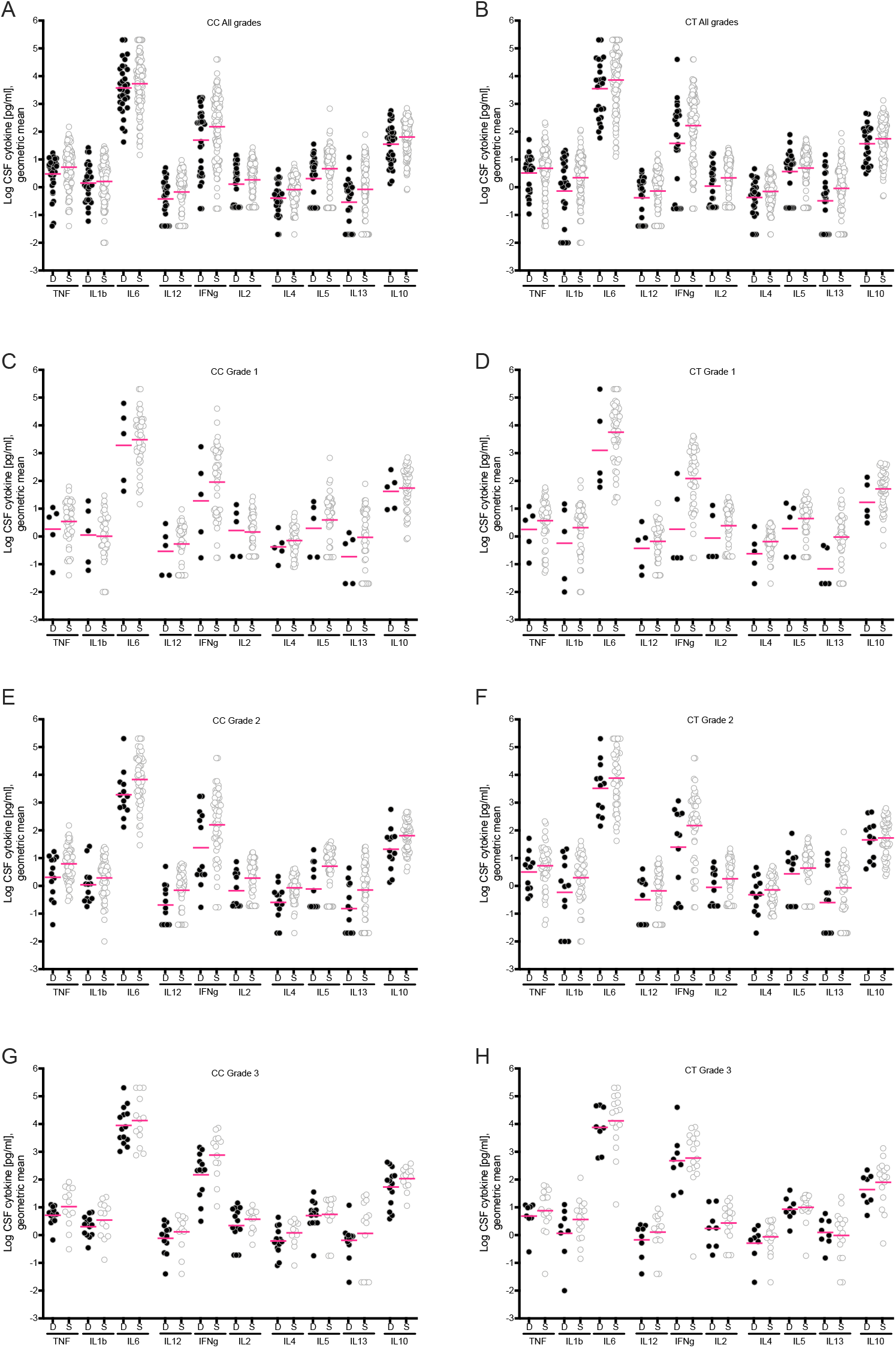
Cytokine levels in HIV-negative CC and CT patients, survivors and non-survivors. (A, B) Cytokine levels in patients who died (D) versus those who survived (S), in CC patients (deaths n=32, survivors n=116) and CT patients (deaths n=25, survivors n=117). (C, D) Cytokine levels in Grade 1 CC (deaths n=5, survivors n=46) and CT (deaths n=5, survivors n=49) patients. (E, F) Cytokine levels in Grade 2 CC (deaths n=13, survivors n= 57) and CT (deaths n=12, survivors n=51) patients. (G, H) Cytokine levels in Grade 3 CC (deaths n=14, survivors n=13) and CT (deaths n=8, survivors n=16) patients. Magenta lines indicate geometric means of non-dithered data. Statistical comparisons of the geometric means were not calculated for these subsets.

**Figure 4 - figure supplement 2.**
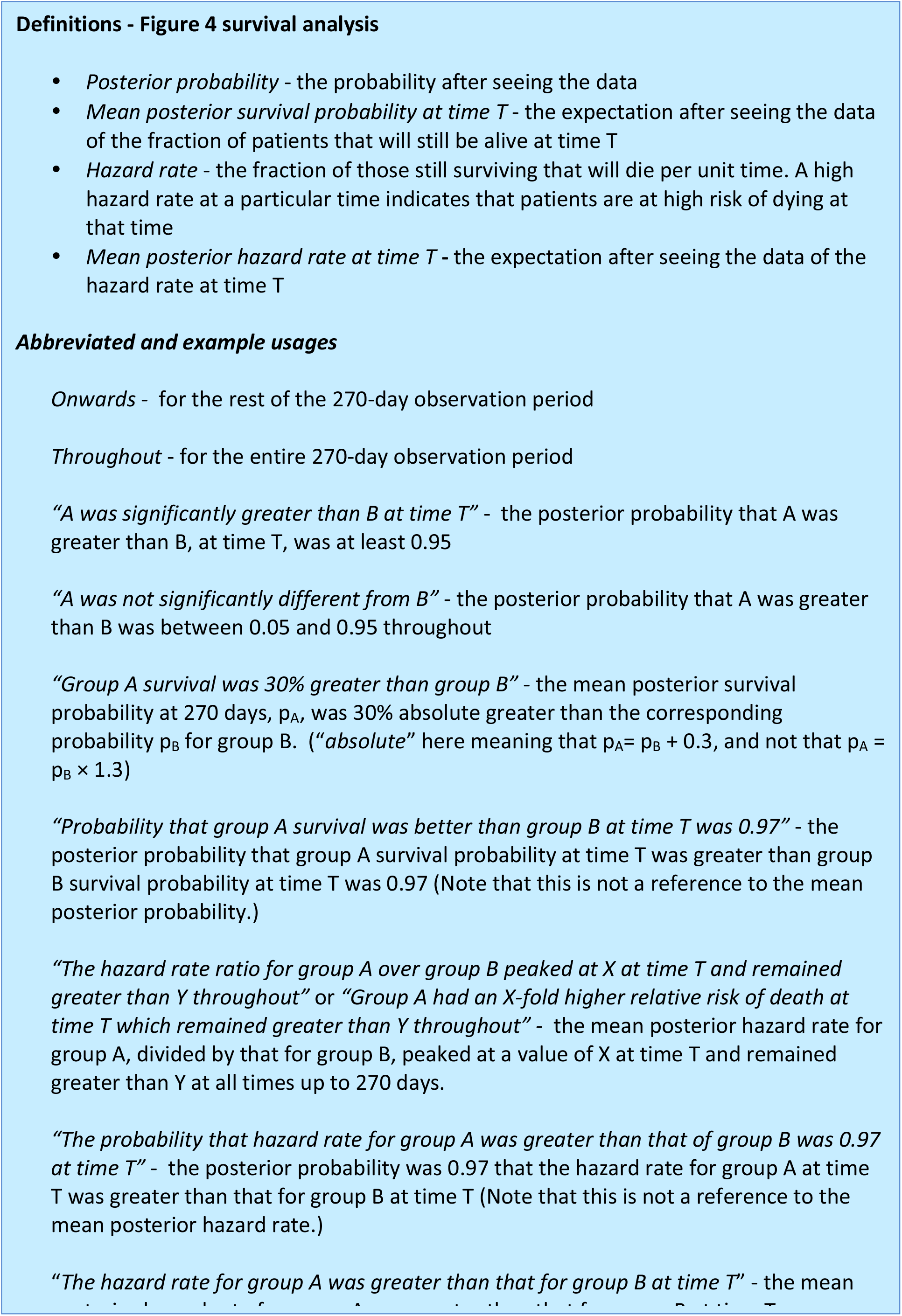

**Supplementary Table 2.** Thuong 2017 cohort separated by hospital.

|  | All of Thuong 2017 cohort (n= 439) |  |
| --- | --- | --- |
|  | Hospital 1 | Hospital 2 |
| **Total n** | 290 | 149 |
| **Age in years**<br>***median (range)*** | 44 (18-93) | 35 (18-86) |
| **BMRC TBM grade**<br>***no. (% of total)*** |  |  |
| 1 | 118 (40.7) | 45 (30.2) |
| 2 | 133 (45.9) | 73 (49.0) |
| 3 | 39 (13.4) | 31 (20.8) |
| **Overall mortality** | 63 (23.5) | 20 (13.4) |
| **Patients with cytokine data** | 162 (55.9) | 144 (96.6) |

**Supplementary Table 3.** Probabilities of selection bias in Hospital 1.

| **Comparison Groups** | **Probability that first group more likely to have cytokine data than second group** |
| --- | --- |
| **CC than non-CC** | **0.999991** |
| **CT than non-CT** | 0.21 |
| **TT than non-TT** | **$3.2 \times 10^{-9}$** |
| **Survivors than Nonsurvivors** | 0.31 |
| **BMRC Grade 1 than Grades 2 and 3** | 0.32 |
| **BMRC Grade 2 than Grades 1 and 3** | 0.29 |
| **BMRC Grade 3 than Grades 1 and 2** | 0.92 |

## Notes

### Competing Interest Statement

The authors have declared no competing interest.

