## Appendix 1 for "Elevated cerebrospinal fluid cytokine levels in tuberculous meningitis predict survival in response to dexamethasone"

### Appendix 1
### An intuitive description of the distribution comparison problem

Roger Sewell

November 2, 2020

#### Contents

**1 Introduction** **1**

**2 An intuitive way of thinking about the distribution comparison problem** **1**

**3 Some warnings** **2**

Version 1.34

#### 1 Introduction

This appendix aims to give an intuitive introduction, for those who haven't encountered Bayesian methods before, to how to think about the distribution comparison problem: given two sets of numbers, how to work out the probability that the numbers in one set are in some sense bigger than those in the other set, when both are distributed according to members of the same, unknown, family of probability distributions.

#### 2 An intuitive way of thinking about the distribution comparison problem

We will be considering various possible families of probability distributions that might describe our data, and within each family, various specific distributions. The reader may perhaps be thinking "we should fit probability density functions to histograms of the data", but we would like to *dissuade* the reader from doing this, and instead adopt the viewpoint of a detective who is trying to work out in retrospect how nature did something.

We think of the data generation process as follows: Assume nature has a kit-bag $F$ of various families of probability distributions it can use. To generate the dataset for a particular cytokine for a subset $S$ of the patients, nature first picks a distribution family $f$ out of its kit-bag. That distribution family has some parameters; e.g. the Gaussian has $\mu$ and $\sigma$ (mean and standard deviation) in the most commonly used parameterisation, while the log-skew-Student has four parameters as we shall see.

So nature next picks a set of parameters $\theta_f$ for $f$, getting a specific distribution. Having done that, some "simple" calculation could (if we had the information on $f$ and $\theta_f$) tell us for example the median $M_S$ of that distribution.

Finally, nature draws random values (one for each patient in the subset) from that distribution to give the data $A$ for that cytokine for that subset $S$ of the patients.

Nature then does the same thing for another subset $T$ of the patients, leaving the investigator with two sets of real numbers $A$ and $B$ representing cytokine levels in the two subsets of patients $S$ and $T$ respectively. Nature also knows (but the investigator doesn't) the values $M_S$ and $M_T$.

The investigator is then cast in the role of detective who wants to know whether $M_S$ or $M_T$ is greater, and how sure he can be of this answer. Bayesian methods then tell him that his first step is to decide how likely it was — before seeing the data ! — that nature picked each of the distribution families in $F$, and his second is to decide how likely it was for each $f$ in $F$ that nature picked each particular possible set of parameters $\theta_f$.

Having done that, some calculations in probability theory enable him to work out what is the probability, given the data, that $M_S$ is bigger than $M_T$. We never have to use traditional curve-fitting, but of course do end up with a belief in the distribution of the observations. Note that because we are dealing here exclusively with continuous distributions, the probability that $M_S = M_T$ exactly is zero — there will always be some small difference between the two, and we may or may not be able to be sure which way round that difference is.

Note also that in what follows we will be assuming that nature has used the *same* family $f$ for making both $A$ and $B$, it is just the parameter values that are different between the two subsets.

### 3 Some warnings

In general Bayesian inference starts from a prior probability distribution on some unknown variables, collects some data, and calculates the posterior distribution on those variables, i.e. the probability distribution that tells us what we then know about those variables. Typically, as here, that is a distribution on a vector of variables containing more than one variable.

In this situation, there are a number of pitfalls for the unwary; we mention the following:

1. It may be tempting to suppose that we should just report the value of the variables at the point where the posterior probability density is greatest; this point is known as the *maximum a posteriori* (MAP) value, and is the mode of the posterior distribution.

   However, unless we are in the very special situation that any error is equally bad as any other[1], this is usually a really bad idea, despite the fact that it has been taught for decades in engineering departments. There are several reasons for this:

   (a) For a single variable $x$, and a function $g$, the mode of the distribution of the function of $x$ is usually *not* at the function of the mode of the distribution of $x$. A good example of this with $g(x) = \log(x)$ is if

   $$P(x) = 2(1 - x)[x \in (0, 1)]$$

   and $y = \log(x)$, in which case

   $$P(y) = 2e^y(1 - e^y)[y < 0].$$

   Then the mode of $x$ is at 0 but the mode of $y$ is at $-0.693 \approx -\log(2) \neq \log(0) = -\infty$.

   (b) If the mode of the distribution of $(\mu, m, s)$ is at $(\hat{\mu}, \hat{m}, \hat{s})$, then the mode of the resulting marginal distribution of $\mu$ is usually *not* at $\hat{\mu}$. A 2-d rather than 3-d example of this is shown in Figure 1, where the mode of $x$ is at about 2.09, well to the left of the mode of $(x, y)$ (whose $x$-coordinate is at 3.55).

[1] In most situations different errors carry differing importance — e.g. concluding that $x$ is 10 times its true value is worse than concluding that it is twice its true value. Similarly concluding that a distribution, that is in truth uniform on an interval, is Gaussian would usually be a less bad error than concluding that it is Cauchy.

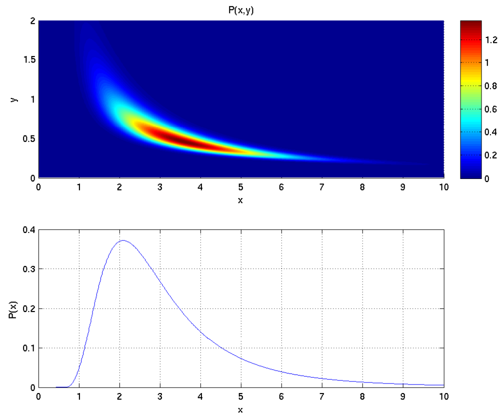


Figure 1: A probability distribution in two variables $(x, y)$, with the marginal distribution in $x$ shown underneath, to illustrate that the projection of the mode of the joint distribution, at 3.55, is in a different place from the mode of the marginal distribution at 2.09.

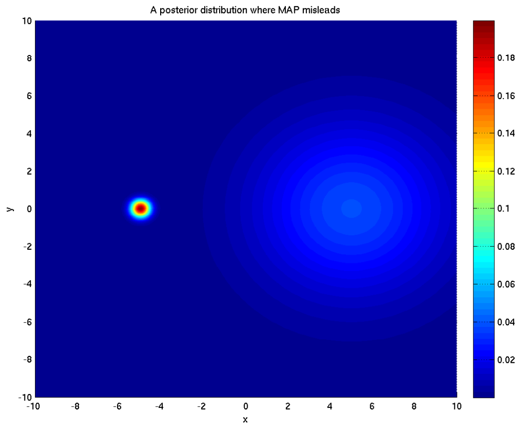


Figure 2: A probability distribution in two variables $(x, y)$ which is bimodal. The MAP value of $(x, y)$ is $(-5, 0)$, but 95% of the probability lies in the broader peak centred at $(+5, 0)$. For another view of this distribution see Figure 3.

(c) Reporting the MAP value is often misleading as far as where the truth is likely to lie — a good example of this is shown in Figure 2, where the MAP value of $x$ is in the middle of the peak centred at $(-5, 0)$, but 95% of the distribution lies in the peak centred at $(+5, 0)$.

For another example, this time from this paper, see section 8.2 of Appendix 2, on testing the software. In this case the MAP often lands not on the correct choice of distribution family, but on one that is under the particular circumstances (parameters) in hand and for the purposes in hand not importantly different from the correct answer; the correct subsequently deduced answer to the problem in hand is correct in all the cases tested.

Nonetheless it is occasionally a reasonable second-best to use MAP, and we have done in this paper for the specific purposes of commenting on software testing and of dithering coincident data values.

2. Various distributions have parameters whose symbols are the same as those in other distributions where they happen to denote specific attributes of the distribution. For example for a Gaussian, $\mu$ is both the mean and the median. But for the log-Gaussian, in the parameterisation used here, $\mu$ is the log of the median but is *not* the log of the mean.

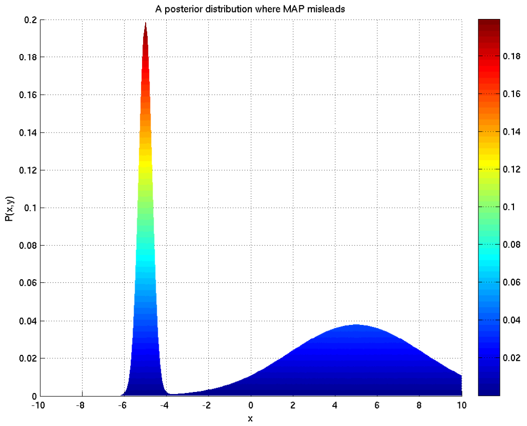


Figure 3: The probability distribution of Figure 2 seen from the direction of $(0, -\infty)$. The MAP value of $(x, y)$ is $(-5, 0)$, but 95% of the probability lies in the broader peak centred at $(+5, 0)$, which is not only much wider than the left-hand peak in the $x$-direction (as shown) but is also much wider in the $y$-direction which cannot be seen in this view.
