## Appendix 2 for "Elevated cerebrospinal fluid cytokine levels in tuberculous meningitis predict survival in response to dexamethasone"

### Appendix 2
### Supplementary Methods

Roger Sewell

November 2, 2020

#### Contents

**1 Introduction — The two problems discussed** **2**

**2 Overview of the distribution comparison problem** **2**

**3 Distributions considered** **3**

**4 Priors on the distribution families** **5**

**5 Priors on the parameters of each distribution family** **5**

**6 Choice of $Q$ - the restricted geometric mean** **22**

**7 Carrying out the calculations** **23**

**8 Testing the software** **23**

**11 References** **31**

Version 1.33

### 1 Introduction — The two problems discussed

The majority of this description of methods focuses on the question of how to work out whether, out of two sets of values, one is bigger than the other — the distribution comparison problem.

However at the end a final section discusses applying similar methods to work out whether, of two probabilities, one is bigger than the other — the probability comparison problem.

Readers who have not encountered Bayesian methods before, and Bayesian model choice in particular, may like to read an informal intuitive introduction in Appendix 1.

### 2 Overview of the distribution comparison problem

We adopt the Bayesian paradigm. We suppose that we are given two sets of observations $A$ and $B$ of real numbers both distributed according to one of a set $F$ of possible distribution families but with different parameter values. Let $Q(A)$ (resp. $Q(B)$) denote some real scalar property of the distribution from which $A$ (resp. $B$) is drawn (e.g. arithmetic or geometric mean, median, etc.). We address the general question "What is the probability that $Q(A) > Q(B)$ given $A$ and $B$ ?". We assume in advance that the posterior probability that $Q(A) = Q(B)$ exactly is zero, so the alternative hypothesis being considered is that $Q(A) < Q(B)$.

This problem can be divided into two parts:

1. First we estimate the probabilities, given the data, that each of the distribution families $f \in F$ gives rise to the data. Mathematically, we calculate

$$P(f|A,B) = \frac{P(f)P(A|f)P(B|f)}{\sum_{f\in F} P(f)P(A|f)P(B|f)}.$$

   This involves quantities such as $P(A|f)$, which we calculate by integrating over all possible parameter choices for this family, weighted by their performance on the observed data:

$$P(A|f) = \int P(A|\theta_f, f)P(\theta_f|f)\, d\theta_f,$$

   where $\theta_f$ is the vector of parameters required by distribution family $f$.

2. Second, we calculate the probability given the data that $Q(A) > Q(B)$, by

$$P(Q(A) > Q(B)|A, B) = \sum_{f \in F} P(f|A, B) P(Q(A) > Q(B)|f, A, B).$$

In words, we sum the fraction of our hypothesis space, now weighted by evidence, in which $Q(A) > Q(B)$.

In order to do this, we also need to be able to calculate $P(Q(A) > Q(B)|f, A, B)$ which is given by

$$P(Q(A) > Q(B)|f, A, B) = \int_{Q(f,\theta_{f,A})>Q(f,\theta_{f,B})} P(\theta_{f,A}|f, A) P(\theta_{f,B}|f, B)\, d(\theta_{f,A}, \theta_{f,B}),$$

where $Q(f, \theta_f)$ denotes the relevant property of the distribution family $f$ with parameter values $\theta_f$.

The remaining ingredients for these calculations are the prior probabilities $P(f)$ and $P(\theta_f|f)$ for each $f \in F$. These will be discussed in sections 4 and 5 below.

Finally, we report a "significant" result if either $P(Q(A) > Q(B)|f, A, B) > 0.95$ or $P(Q(A) > Q(B)|f, A, B) < 0.05$.

All these methods are standard and long-established; see e.g. [1] chapter 28 or [2].

### 3 Distributions considered

Based on previous experience with a variety of real world distributions, we chose several reasonable possibilities, and some that can be considered negative controls — the Gaussian, Student, and skew-Student — which fail to exhibit vital characteristics of the data, (e.g., Gaussian and Student distributions are symmetric and put non-zero probability on negative values) and should be rejected if the software is correct. In particular we include the Gaussian because it is a common (but usually false) assumption. The reader should note that these are a set of reasonable possibilities; it is not practical to consider the infinitely many different families of continuous distributions on the positive reals that exist.

When discussing the following distributions the reader may be tempted to assume that e.g. $\mu$ denotes the arithmetic mean of the distribution. Although this is true for some of them (e.g. the Gaussian, or the Student when the shape parameter $m > \frac{1}{2}$), in general it is not the case: e.g. the Student has no mean when $m \leq \frac{1}{2}$, and for the log-Gaussian $\mu$ is the log of the geometric mean.

Accordingly we define a set $F$ of possible distributions to be the set of the following families, where in each case the vector of parameters following the $|$ on the left-hand side forms the vector $\theta_f$ whose posterior distribution will be inferred as part of the procedure (for definitions of symbols not appearing on the LHS of the definitions see after the whole list):

- The Gaussian distribution $P(x|\mu, s) = \sqrt{\frac{s}{2\pi}} e^{-\frac{1}{2}sy^2}$ in which $\mu$ is the mean and median and $s$ is the scale, i.e. the reciprocal of the variance;
- The Student distribution $P(x|\mu, s, m) = \frac{1}{\sqrt{2\pi}} \frac{\Gamma(m+\frac{1}{2})}{\Gamma(m)} \frac{r^m}{(r+\frac{1}{2}y^2)^{m+\frac{1}{2}}}$ in which $\mu$ is the median, $s$ is the scale (but not necessarily the reciprocal of the variance), and $m$ is a shape parameter which reduces the heaviness of the tails as it gets bigger;
- The log-Gaussian distribution $P(x|\mu, s) = \sqrt{\frac{s}{2\pi}} \frac{1}{x} e^{-\frac{1}{2}sz^2}$ in which $\mu$ is the log of the geometric mean and $s$ is the scale, now the reciprocal of the variance of the log value;

- The log-Student distribution $P(x|\mu,s,m) = \frac{1}{\sqrt{2\pi}}\frac{\Gamma(m+\frac{1}{2})}{x\Gamma(m)}\frac{r^m}{(r+\frac{1}{2}z^2)^{m+\frac{1}{2}}}$ in which $\mu$ is the log of the median, $s$ is the scale (don't assume anything further), and $m$ is a shape parameter;
- The Gamma distribution $P(x|m,s) = \frac{r^m}{\Gamma(m)}x^{m-1}e^{-rx}$ where $m$ is a shape parameter which makes the distribution narrower as it gets bigger and $s$ is the scale;
- The inverse-Gamma distribution $P(x|m,s) = \frac{r^m}{\Gamma(m)}x^{-m-1}e^{-r/x}$, where the reciprocal of the value is Gamma-distributed;
- The skew-Student distribution (also known as the non-central Student $t$ distribution) [3]

$$P(x|\xi,m,s,\eta) = \frac{r^m}{\Gamma(m)}\sqrt{\frac{1}{2\pi}}\frac{1}{(r+\frac{1}{2}y^2)^{m+\frac{1}{2}}}e^{-\frac{1}{2}\nu^2}\times$$
$$\left(\Gamma(m+\frac{1}{2})_1F_1\left(m+\frac{1}{2},\frac{1}{2},\frac{(y\nu)^2}{4(r+\frac{1}{2}y^2)}\right)+\Gamma(m+1)\frac{y\nu}{\sqrt{(r+\frac{1}{2}y^2)}}{}_1F_1\left(m+1,\frac{3}{2},\frac{(y\nu)^2}{4(r+\frac{1}{2}y^2)}\right)\right);$$

where now $\xi$ is a location parameter, $m$ is a shape parameter (that also affects the skewness), $s$ is a scale parameter, and $\nu$ is the skewness parameter;
- The log-skew-Student distribution

$$P(x|\xi,m,s,\eta) = \frac{r^m}{x\Gamma(m)}\sqrt{\frac{1}{2\pi}}\frac{1}{(r+\frac{1}{2}z^2)^{m+\frac{1}{2}}}e^{-\frac{1}{2}\nu^2}\times$$
$$\left(\Gamma(m+\frac{1}{2})_1F_1\left(m+\frac{1}{2},\frac{1}{2},\frac{(z\nu)^2}{4(r+\frac{1}{2}z^2)}\right)+\Gamma(m+1)\frac{z\nu}{\sqrt{(r+\frac{1}{2}z^2)}}{}_1F_1\left(m+1,\frac{3}{2},\frac{(z\nu)^2}{4(r+\frac{1}{2}z^2)}\right)\right);$$

where now the log of the value is skew-Student-distributed;
- The Gamma power distribution $P(x|m,s,k) = |k|\frac{r^m}{\Gamma(m)}x^{mk-1}e^{-rx^k}$ where $x^k$ is Gamma-distributed;

where we make the following identifications:

- $r = ms^k$ (with $k$ taken to be 1 if not applicable, i.e. for all but the Gamma power distribution);
- $y = x - \mu$;
- $z = \log(x) - \mu$;
- $\mu = \xi - \frac{\Gamma(m-\frac{1}{2})}{\Gamma(m)}\nu\sqrt{r}$ for the cases of the (log-)skew-Student only;
- $\nu = \frac{\eta^3+\alpha\eta}{\eta^2+\beta}$;
- $\alpha = 100, \beta = 1000$;.

and where $\Gamma$ denotes the Gamma function defined by $\Gamma(m) = \int_0^\infty x^{m-1}e^{-x}\,dx$, and where ${}_1F_1$ is the hypergeometric function defined by

$${}_1F_1(a,b,z) = \sum_{n=0}^{\infty}\frac{\Gamma(a+n)\Gamma(b)z^n}{\Gamma(a)\Gamma(b+n)n!}.$$

In the case that the data are inherently all positive, as in the present application, those distributions whose support includes the negative reals (i.e. the Gaussian, Student, and skew-Student) should be seen as "straw men"; they cannot possibly be the correct distributions of the data. However, we include them both because Gaussianity is a common (but usually false) assumption, and in order to demonstrate the ability of the Bayesian approach to reject these possibilities.

We note that all these distributions have several possible ways of parameterising them; e.g. the Gaussian can be parameterised by $\mu, \sigma$ where $\sigma$ is the standard deviation instead of by $\mu, s$ where $s$ is the reciprocal of the variance. Unfortunately which parameterisation is used *does* make a difference to grid resolution needed (and also to speed of convergence of MCMC algorithms) even though it doesn't make any difference to the underlying nature of the distributions; the parameterisations shown are those we recommend.

### 4 Priors on the distribution families

We consider two priors on the distribution families:

1. $P(f) = \frac{1}{9}$ for all $f \in F$ (i.e. each distribution equally likely).
2. $P(f) = \begin{cases} 1 & (f \text{ is the log-skew-Student family}) \\ 0 & \text{(otherwise).} \end{cases}$, i.e. only the log-skew-Student is considered.

It will later turn out that the conclusions are the same for both priors, even though they are very different. In other words, the log-skew-Student family turns out to be so much more likely (given the data) than the others that any reasonable prior doesn't make any difference.

### 5 Priors on the parameters of each distribution family

#### 5.1 Reference point and units

We arbitrarily use the geometric mean of the data values for a particular cytokine over all subsets being compared as an absolute scaling factor. In the following that is assumed to be 1.

Logarithms are natural logarithms, i.e. to base $e$.

In general the choice of parameters for the priors (also known as hyperparameters) was made knowing the basics about the data, namely that it was all positive, but with a wide dynamic range of about $10^5$-fold within each cytokine, and with some asymmetry even in the log domain. Beyond that the choices were made to allow a very wide range of distributions for the data to be possible.

#### 5.2 Gaussian

We use the joint conjugate prior to $(\mu, s)$ given by

$$P(s|m_s, r_s) = \frac{r_s^{m_s}}{\Gamma(m_s)} s^{m_s - 1} e^{-r_s s} [s > 0],$$

$$P(\mu|s, K_\mu, \mu_\mu) = \sqrt{\frac{K_\mu s}{2\pi}} e^{-\frac{1}{2} K_\mu s (\mu - \mu_\mu)^2},$$

where $r_s = m_s s_s$, and set the parameters to $m_s = 4, s_s = 400, K_\mu = 0.2, \mu_\mu = 150$.

This results in the joint distribution for $(\mu, s)$ shown in Figure 1, while samples from this family are then shown in Figure 2 on a linear axis and Figure 3 on a logarithmic axis. Note that the prior has been chosen so that there is only a very small chance of distributions allocating more than a few percent of the total probability to the negative reals; this necessarily results in the centre of the distribution family

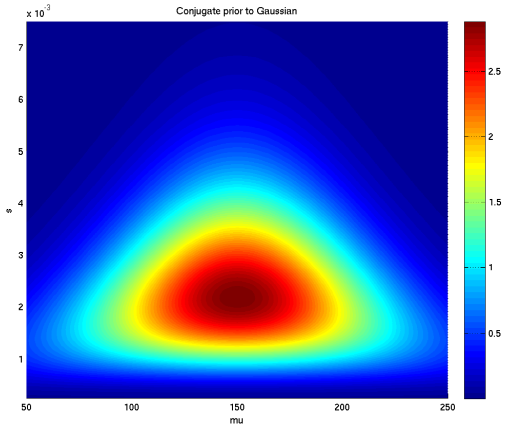

Figure 1: Prior distribution adopted for the parameters $\mu$ and $s$ of the Gaussian distribution family

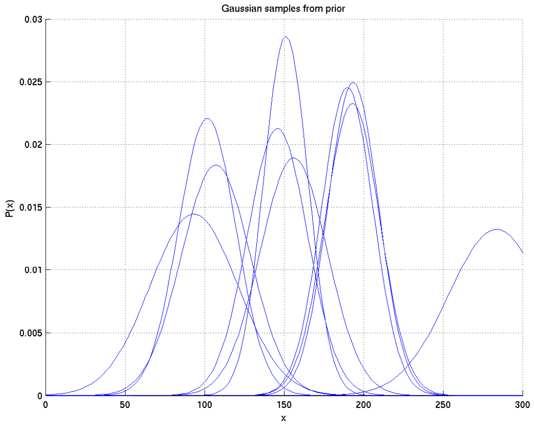

Figure 2: Samples of Gaussian distributions drawn from the prior

being much more positive than we expect the geometric mean of our real life data to be, but to be able to cover the larger samples (which range up to several hundred times the geometric mean).

If we had allowed a substantial fraction of these distributions to be on the negative reals (e.g. if all had 3% there), then we would be penalising the Gaussian family by a factor of $0.97^{250} = 5 \times 10^{-4}$ relative to positive-support families, so Gaussian would get very low posterior probability in step 1 of section 2 above, and there would be even less point including it.

##### 5.3 Log-Gaussian

We adopt the same prior family as for the Gaussian in the previous section, but now with the parameters $m_s = 1.5, s_s = 0.4, K_\mu = 0.2, \mu_\mu = 0$. This results in the joint distribution for $(\mu, s)$ shown in Figure 4, while samples from the family are then shown in Figures 5 and 6.

##### 5.4 Student

In this case we use independent priors on each of the parameters $\mu, m, s$. For $\mu$ we use a Gaussian with parameters $\mu_\mu, s_\mu$ (i.e. mean $\mu_\mu$, variance $1/s_\mu$). For $m$ we use the prior conjugate to the Gamma with

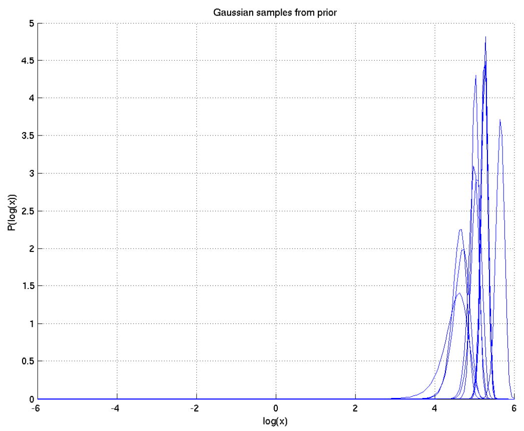

Figure 3: Samples of Gaussian distributions drawn from the prior

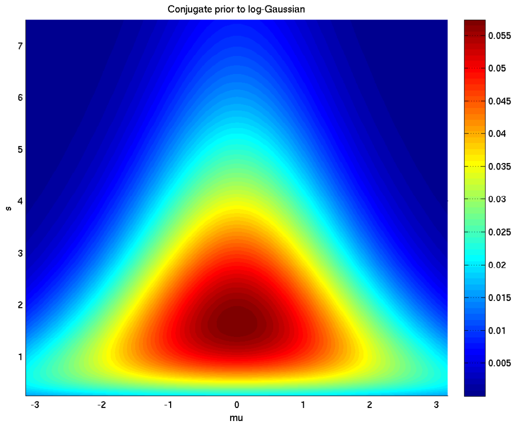

Figure 4: Prior distribution adopted for the parameters $\mu$ and $s$ of the log-Gaussian distribution family

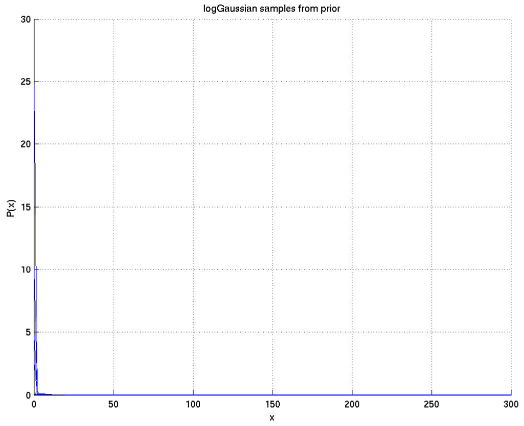

Figure 5: Samples of log-Gaussian distributions drawn from the prior

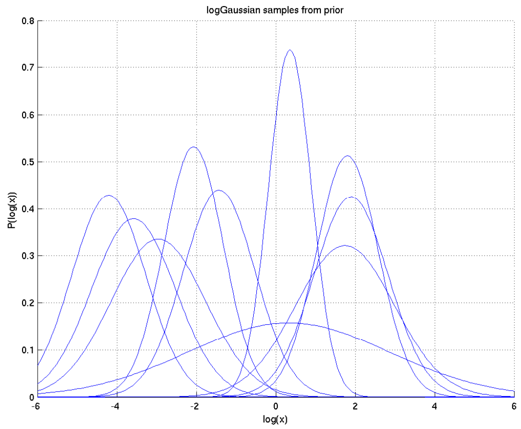

Figure 6: Samples of log-Gaussian distributions drawn from the prior

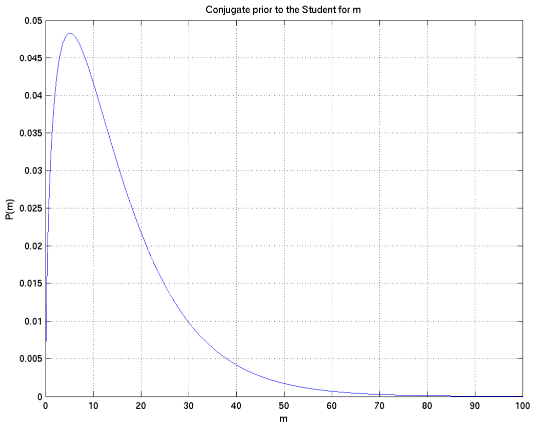

Figure 7: Prior distribution adopted for the parameter $m$ of the Student distribution family

respect to the parameters $m, s$, namely

$$P(m|a_m, b_m) \propto \frac{e^{-m(a_m+b_m)} m^{b_m m}}{\Gamma(m)^{b_m}} [m > 0],$$

where $a_m > 0, b_m > -1$. For $s$ we use a Gamma prior

$$P(s|m_s, s_s) = \frac{r_s^{m_s}}{\Gamma(m_s)} s^{m_s - 1} e^{-r_s s} [s > 0],$$

where $r_s = m_s s_s$, and both $r_s$ and $s_s$ are positive. We set the parameters to $a_m = 0.1, b_m = 1, \mu_\mu = 150, s_\mu = 0.000625, m_s = 1.1, s_s = 0.0025$.

This gives priors on $\mu, m, s$ shown in Figures 9, 7, 8 and samples shown in Figures 10 and 11.

#### 5.5 Log-Student

We adopt the same prior families as for the Student above, but with the following parameters: $a_m = 0.2, b_m = 4, \mu_\mu = 0, s_\mu = 1, m_s = 1.1, s_s = 1$. This results in prior distributions on the parameters shown in Figures 12, 13, 14 and samples shown in Figures 15 and 16.

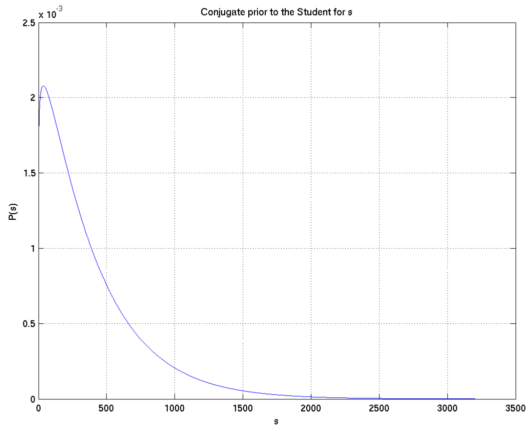

Figure 8: Prior distribution adopted for the parameter $s$ of the Student distribution family

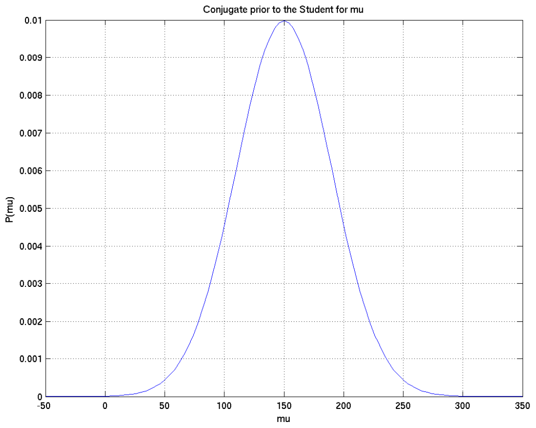

Figure 9: Prior distribution adopted for the parameter $\mu$ of the Student distribution family

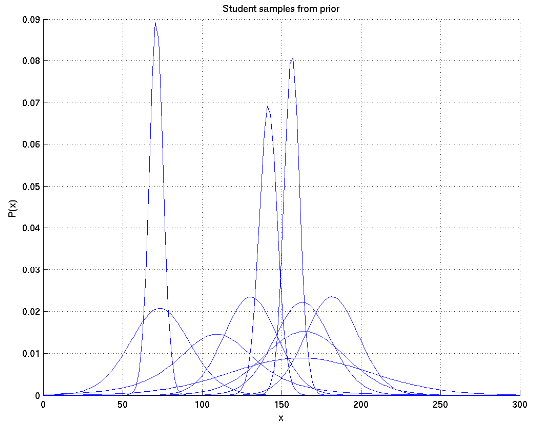

Figure 10: Samples of Student distributions drawn from the prior

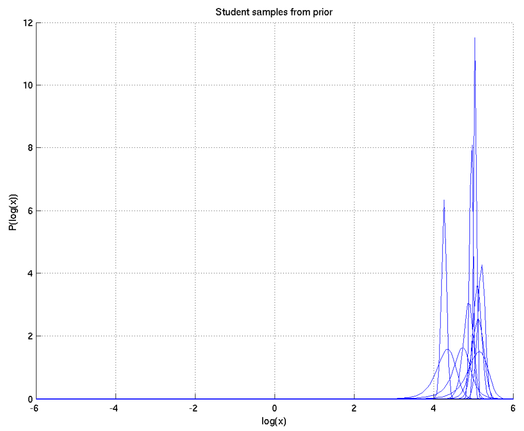

Figure 11: Samples of Student distributions drawn from the prior

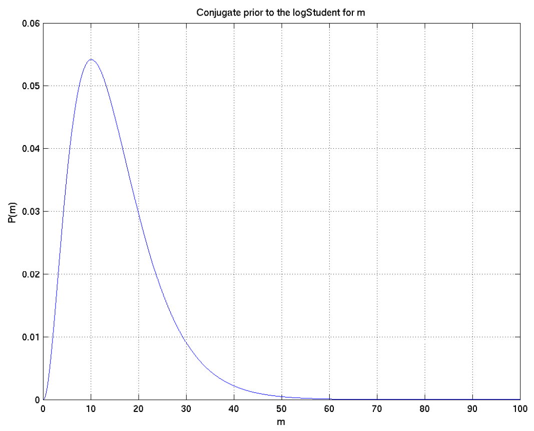

Figure 12: Prior distribution adopted for the parameter $m$ of the log-Student distribution family

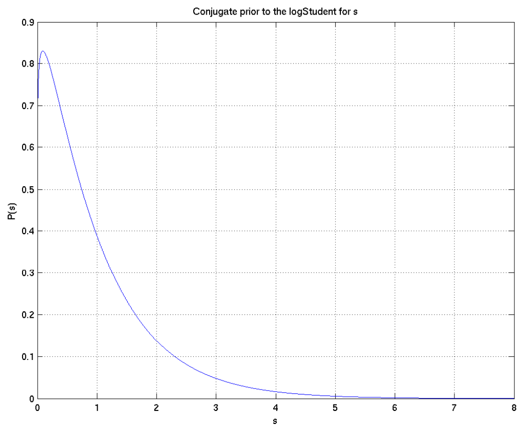

Figure 13: Prior distribution adopted for the parameter $s$ of the log-Student distribution family

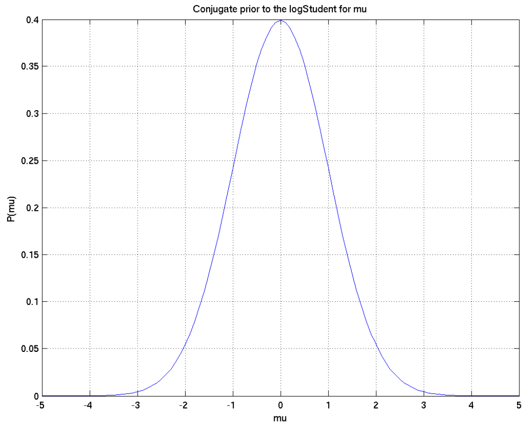

Figure 14: Prior distribution adopted for the parameter $\mu$ of the log-Student distribution family

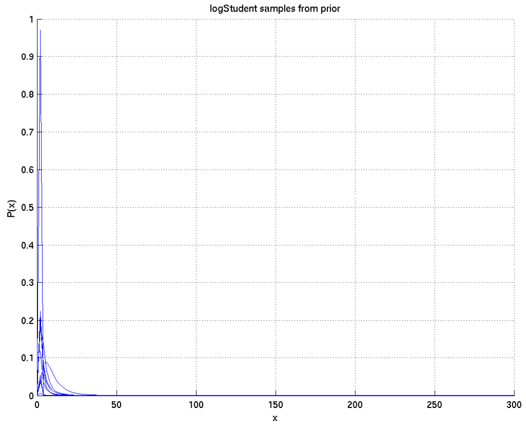

Figure 15: Samples of log-Student distributions drawn from the prior

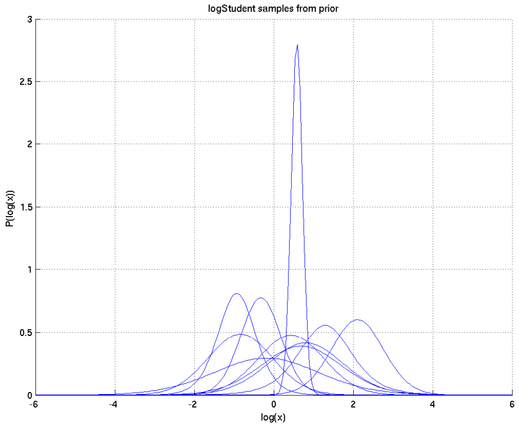

Figure 16: Samples of log-Student distributions drawn from the prior

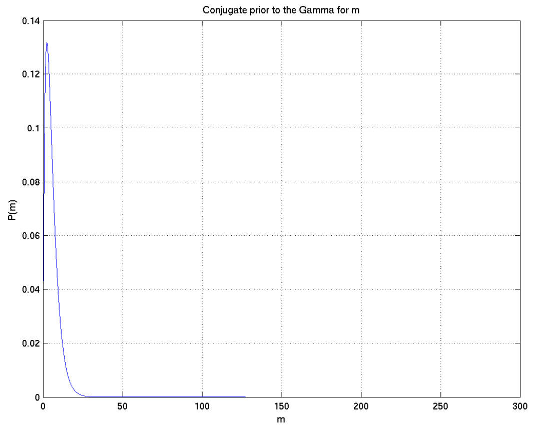

Figure 17: Prior distribution adopted for the parameter $m$ of the Gamma distribution family

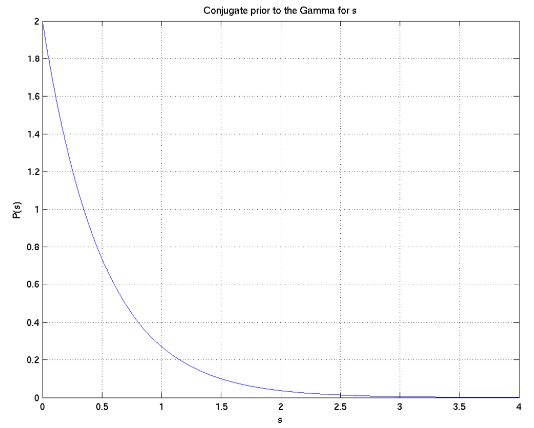

Figure 18: Prior distribution adopted for the parameter $s$ of the Gamma distribution family

#### 5.6 Gamma

We again adopt independent priors for the two parameters. For $m$ we use the prior conjugate to the Gamma with respect to the parameters $m, s$, namely

$$P(m|a_m, b_m) \propto \frac{e^{-m(a_m+b_m)} m^{b_m m}}{\Gamma(m)^{b_m}} [m > 0],$$

where $a_m > 0, b_m > -1$. For $s$ we use a Gamma prior

$$P(s|m_s, s_s) = \frac{r_s^{m_s}}{\Gamma(m_s)} s^{m_s - 1} e^{-r_s s} [s > 0],$$

where $r_s = m_s s_s$, and both $r_s$ and $s_s$ are positive. We set the parameters to $a_m = 0.02, b_m = 1.5, m_s = 1, s_s = 2$, which results in the prior distributions shown in Figures 17 and 18 and the samples shown in Figures 19 and 20.

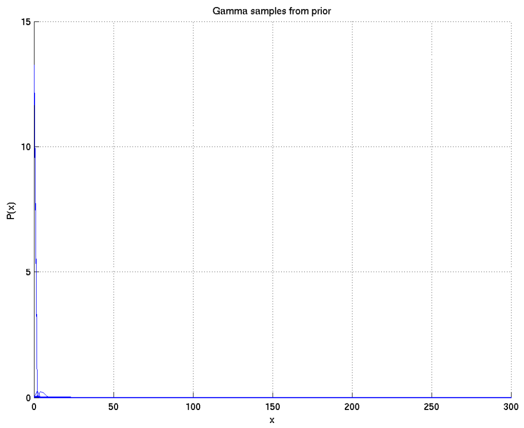

Figure 19: Samples of Gamma distributions drawn from the prior

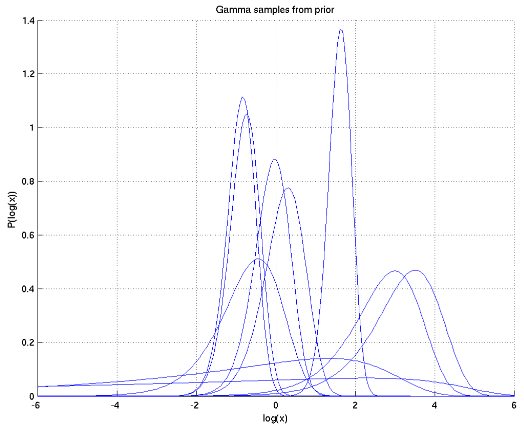

Figure 20: Samples of Gamma distributions drawn from the prior

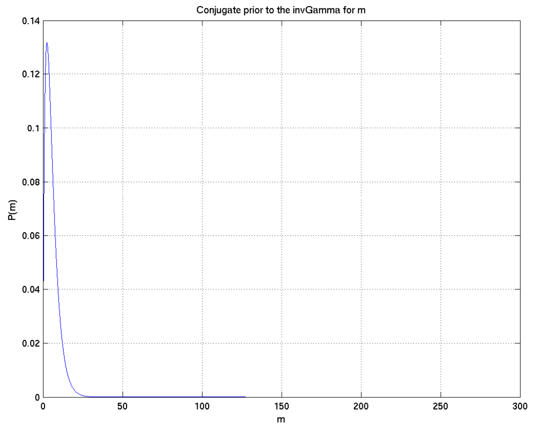

Figure 21: Prior distribution adopted for the parameter $m$ of the inverse-Gamma distribution family

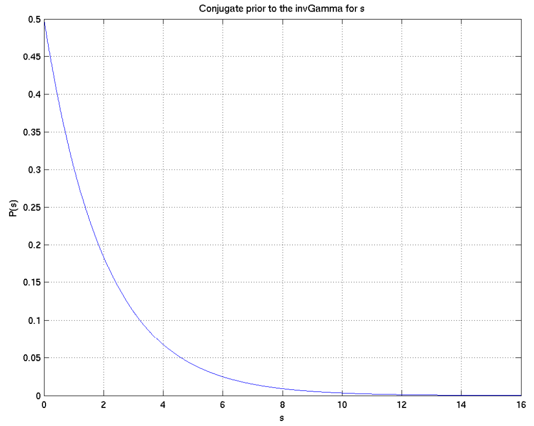

Figure 22: Prior distribution adopted for the parameter $s$ of the inverse-Gamma distribution family

#### 5.7 Inverse Gamma

We use the same priors as for the Gamma distribution, but with the parameters set to $a_m = 0.02, b_m = 1.5, m_s = 1, s_s = 0.5$. This results in the priors on the parameters shown in Figures 21 and 22 and the samples shown in Figures 23 and 24.

#### 5.8 Skew Student

The parameter $s$ has the same prior families as the Student, while $\xi$ is Gaussian with mean $\mu_\xi$ and scale $s_\xi$. We give $m$ the prior family

$$P(m|a_m, b_m) \propto \frac{e^{-m(a_m+b_m)} m^{b_m m}}{\Gamma(m)^{b_m}} [m > \frac{1}{2}].$$

We give $\eta$ the distribution

$$P(\eta|s_\nu, \alpha, \beta) = \sqrt{\frac{s_\nu}{2\pi}} e^{-\frac{1}{2} s_\nu \left(\frac{\eta^3+\alpha\eta}{\eta^2+\beta}\right)^2} \frac{\eta^4 + (3\beta - \alpha)\eta^2 + \alpha\beta}{(\eta^2 + \beta)^2},$$

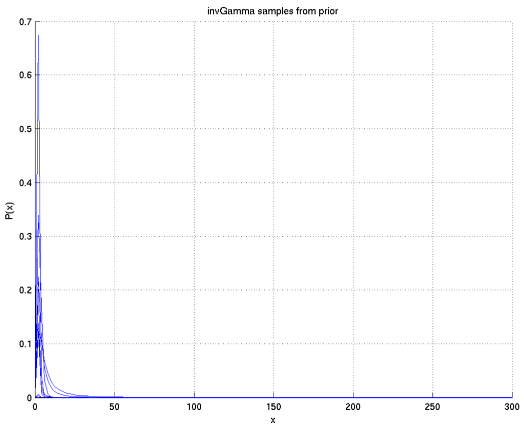

Figure 23: Samples of inverse-Gamma distributions drawn from the prior

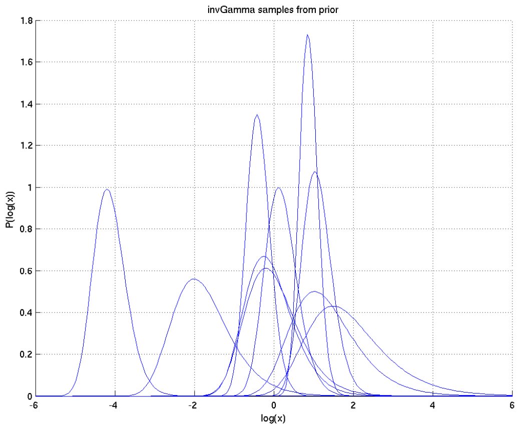

Figure 24: Samples of inverse-Gamma distributions drawn from the prior

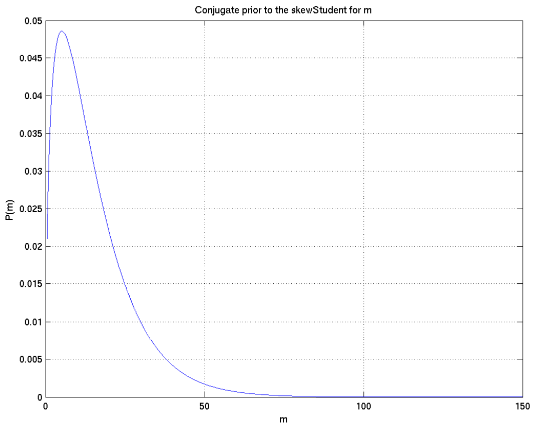

Figure 25: Prior distribution adopted for the parameter $m$ of the skew-Student distribution family

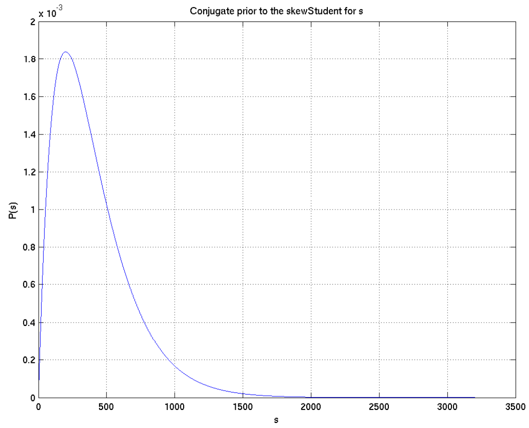

Figure 26: Prior distribution adopted for the parameter $s$ of the skew-Student distribution family

which amounts to making $\nu$ Gaussian with zero mean and scale $s_\nu$. The parameters $\alpha$ and $\beta$ have no effect on the prior distribution of $\nu$, but instead adjust the parameterisation to enable easier numerical integration by making the posterior density smoother, while not changing the posterior distribution.

We adopt the parameters $\alpha = 100, \beta = 1000, s_\nu = 0.25, a_m = 0.1, b_m = 1, m_s = 2, s_s = 0.0025, \mu_\xi = 150, s_\xi = 0.000625$. This results in the prior distributions shown in Figures 25, 26, 27, and 28, and the samples shown in Figures 29 and 30.

Finally we note that in all the final runs for submitted version of the paper the (non-log) skew-Student distribution was omitted from the list of possibilities examined; this was because

- including it nearly doubled the running time (as almost all the running time was spent on the skew-Student and log-skew-Student) and
- preliminary runs had shown that it was not a serious contender to explain these obviously logarithmically-distributed datasets.

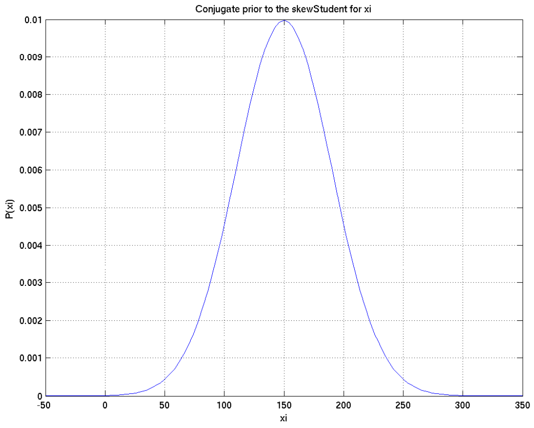

Figure 27: Prior distribution adopted for the parameter $\xi$ of the skew-Student distribution family

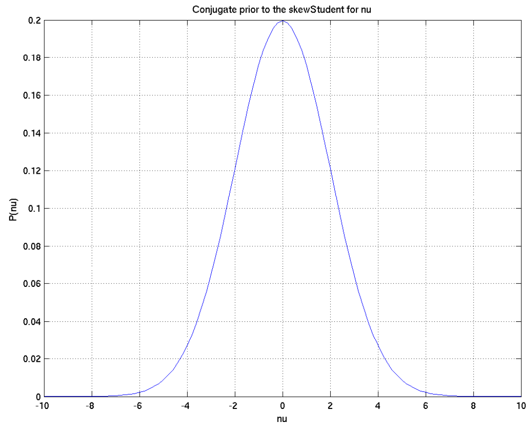

Figure 28: Prior distribution adopted for the parameter $\nu$ of the skew-Student distribution family

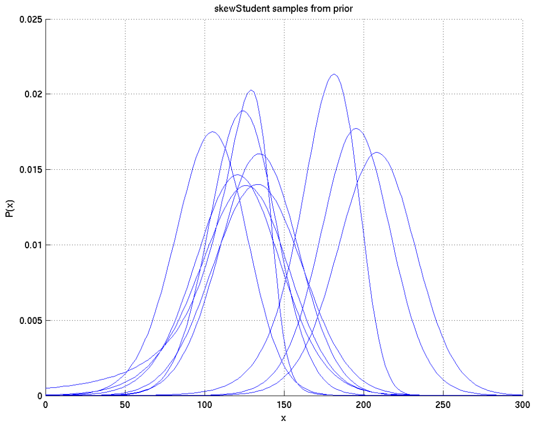

Figure 29: Samples of skew-Student distributions drawn from the prior

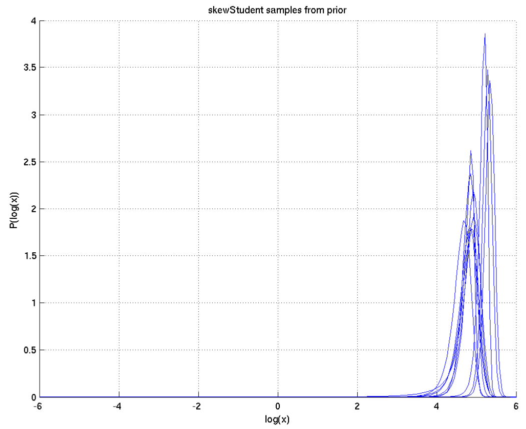

Figure 30: Samples of skew-Student distributions drawn from the prior

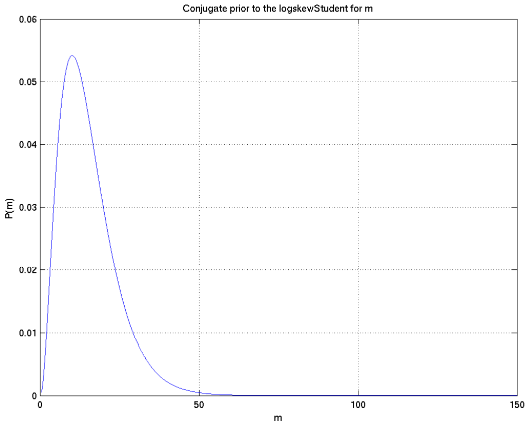

Figure 31: Prior distribution adopted for the parameter $m$ of the log-skew-Student distribution family

#### 5.9 Log-skew-Student

The prior distribution families for the log-skew-Student were chosen to be the same as for the skew-Student, but with the parameters $\alpha = 100, \beta = 1000, s_\nu = 0.025, a_m = 0.2, b_m = 4, m_s = 1.1, s_s = 1, \mu_\xi = 0, s_\xi = 1$. This gives the prior distributions on the parameters shown in Figures 31, 32, 33, and 34, and the samples shown in Figures 35 and 36.

#### 5.10 Gamma power

We take the priors on $m$ and $s$ from the same families with the same parameter values as for the Gamma distribution, except that we set $s_s = 1$.

For $k$ we again choose the conjugate distribution, in this case $P(k|N, a, b, c) \propto k^a \prod_{n=1}^{N} b_n^{kc_n} e^{-c_n b_n^k} [k > 0]$, where $N$ is a positive integer, $a$ a positive scalar, and $b, c$ are $N$-dimensional vectors of positive reals.

We take $N = 2, a = 1, b = (0.2, 0.2), c = (0.5, 0.5)$, giving the distribution on $k$ shown in Figure 39 and the sample distributions shown in Figures 40 and 41.

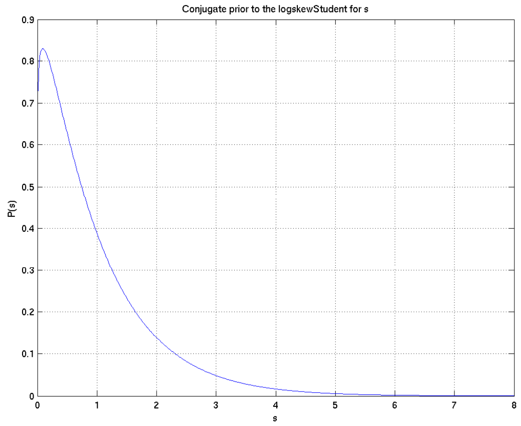

Figure 32: Prior distribution adopted for the parameter $s$ of the log-skew-Student distribution family

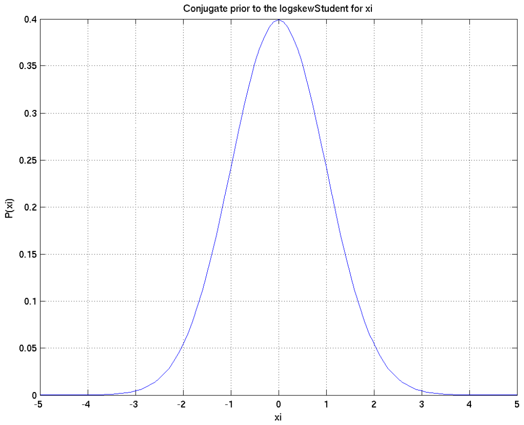

Figure 33: Prior distribution adopted for the parameter $\xi$ of the log-skew-Student distribution family

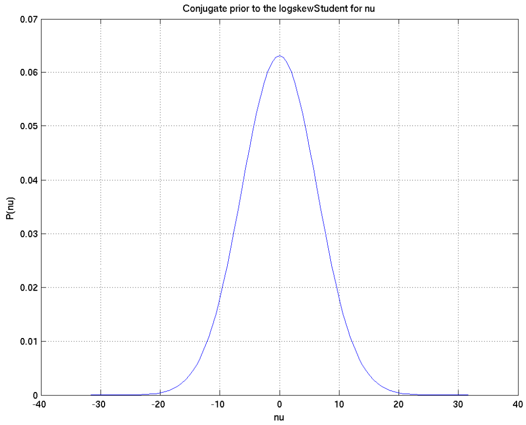

Figure 34: Prior distribution adopted for the parameter $\nu$ of the log-skew-Student distribution family

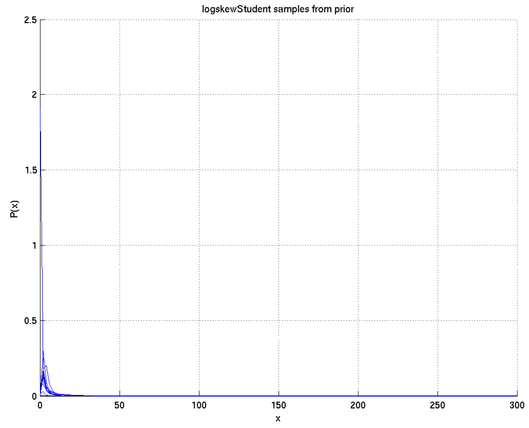

Figure 35: Samples of log-skew-Student distributions drawn from the prior

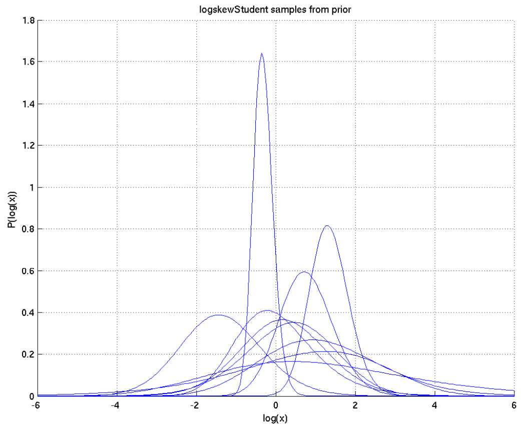

Figure 36: Samples of log-skew-Student distributions drawn from the prior

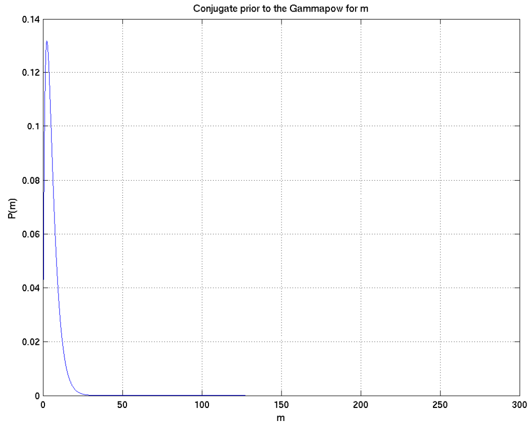

Figure 37: Prior distribution adopted for the parameter $m$ of the Gamma power distribution family

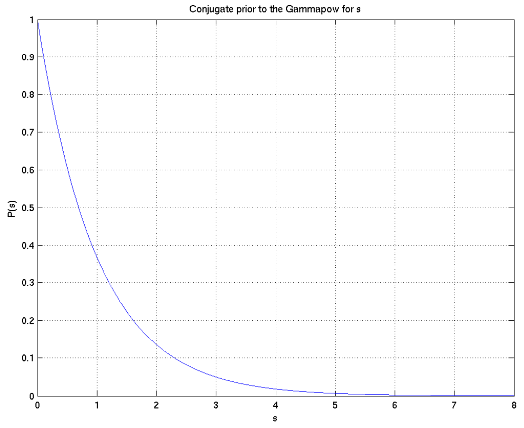

Figure 38: Prior distribution adopted for the parameter $s$ of the Gamma power distribution family

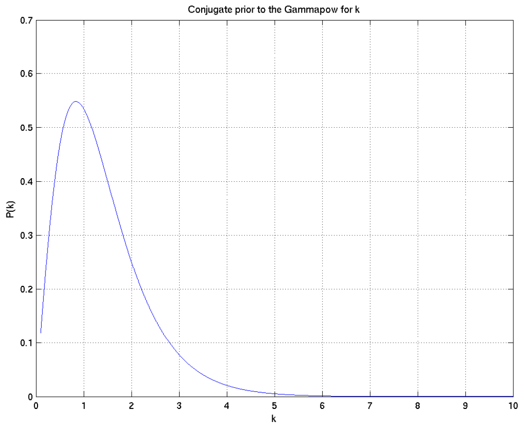

Figure 39: Prior distribution adopted for the parameter $k$ of the Gamma power distribution family

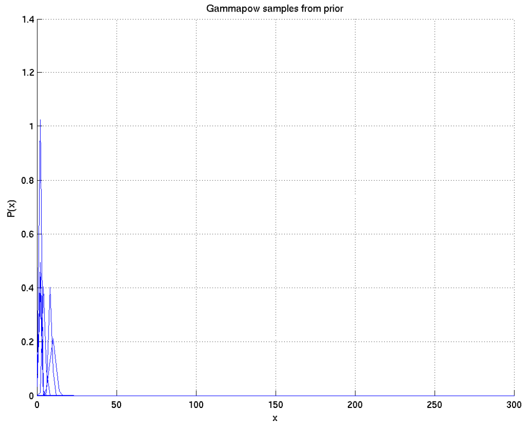

Figure 40: Samples of Gamma power distributions drawn from the prior

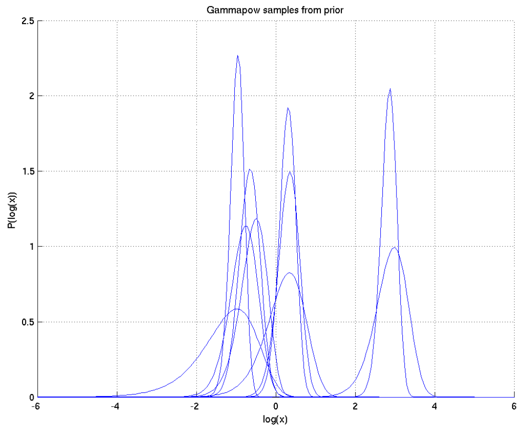

Figure 41: Samples of Gamma power distributions drawn from the prior

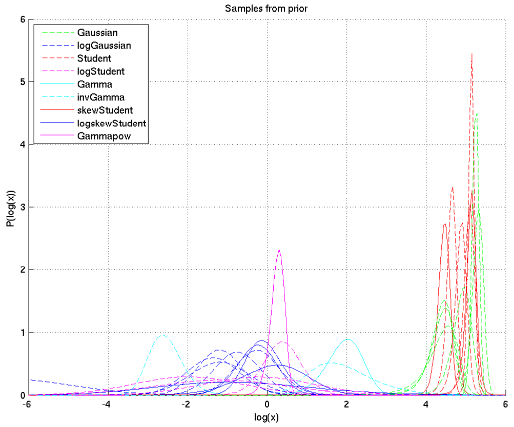

Figure 42: Samples of distributions drawn from the overall prior.

##### 5.11 Resulting priors on distributions

Overall, the priors defined above give us a prior on distributions (as opposed to families) from which samples are illustrated in Figures 42 and 43. For the alternative prior where only log-skew-Students are considered, see Figure 36.

#### 6 Choice of $Q$ - the restricted geometric mean

A further decision that needs to be made is which property of the distribution to use as $Q(f, \theta_f)$. Ordinarily one might expect the mean or median to be used. However, both have drawbacks in this particular situation.

In particular the median will be unaffected by e.g. the top 40% of the distribution's values being doubled, so is insensitive to changes that could be very important.

On the other hand, for five of the distributions being considered, including the one (log-skew-Student) which is will turn out to be by far the most likely single family to give rise to the distribution of all these cytokines, the mean is unfortunately ill-behaved. For the log-(skew)-Student, though the mode is

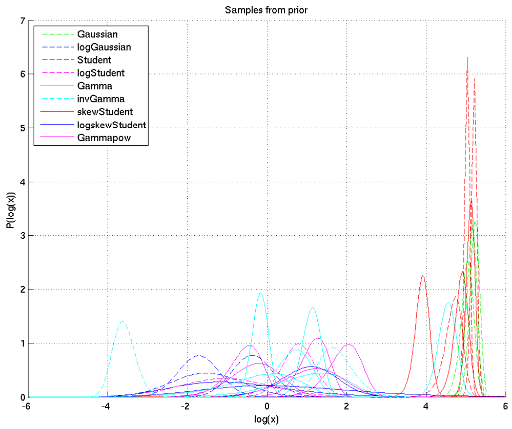

Figure 43: Samples of distributions drawn from the overall prior.

always zero, the mean is always infinite. For the (skew-)Student, the mean is infinite for values of the shape $m$ less than or equal to 0.5, and for the inverse-Gamma it is infinite for values of $m$ less than or equal to 1.

While the geometric mean in some ways behaves better, it does not yet avoid all these problems.

However, by considering instead the restricted geometric mean, i.e. the geometric mean of the part of the distribution between two fixed values $\check{x}$ and $\hat{x}$, the same for all groups being compared, and which are wide enough to encompass e.g. 99% of the probability of all the arising distributions under the posterior and all the observed data values, we obtain a well-behaved and consistent measurement that will capture any likely differences that would be of importance. This we therefore choose as our measurement $Q$, setting $\check{x} = e^{-6} \approx 0.0025$ times, and $\hat{x} = e^{6} \approx 400$ times the geometric mean of all the observed values of a particular cytokine, which choice turns out to satisfy the above criteria.

### 7 Carrying out the calculations

In order to carry out the calculations given in section 2 above, we carry out numerical integration on a grid. An automated algorithm was used to determine the grid values in such a way as to include all parts of the distribution with marginal 1-d density in any parameter at least 0.01 times the peak density, and with 40 grid points in each dimension other than that for $m$ for the (log-)skew-Student families, where 20 grid points were used.

The property $Q(f, \theta)$ used was the mean of that part of the distribution between $e^{-6}$ and $e^{+6}$ times the geometric mean of the data from all subgroups of the relevant cytokine; this was evaluated on the given grid for $\theta$ and a grid of 200 points log-uniformly spaced over this range.

Matlab was used as the main programming language, with C mex code written to speed up the inner loops (specifically the evaluation of the logarithm of the hypergeometric function).

### 8 Testing the software

#### 8.1 Adequacy of grid resolution

Adequacy of grid resolution was confirmed by visualising the posterior distributions of the parameters for each distribution family and dataset. We illustrate this by showing that for the log-skew-Student

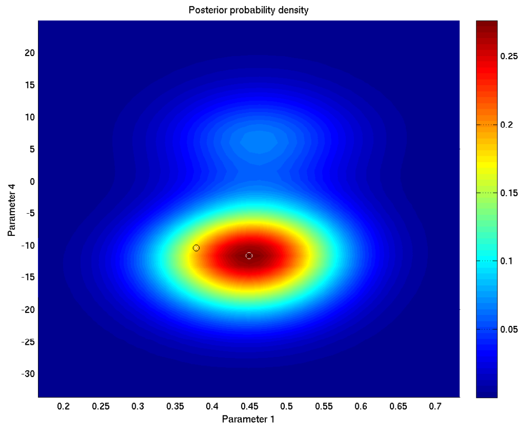

Figure 44: Joint 2-d marginal posterior on parameters $\xi$ and $\eta$ when inferring parameters of a log-skew-Student model for a synthetic truly log-skew-Student dataset, showing adequacy of grid resolution. The projection of the MAP value of the parameters $(\xi, m, s, \eta)$ is shown with a white circle and black x and happens to approximately coincide with the mode of this 2-d marginal, while the projection of the true parameter values is shown with a black circle and white x.

inference on a log-skew-Student test dataset in Figure 44. The only times when inadequate gridding was apparent were when the posterior on the (non-log-) skew Student parameters was evaluated for a logarithmically distributed dataset, as for example the data shown in Figures 46 and 47 which gives Figure 45; but it was nonetheless clear that the skew-Student family was very unlikely to be responsible for this dataset as its posterior probability was less than $10^{-48}$ times that of the log-skew-Student.

#### 8.2 Inference of distribution family and direction of comparison

In order to test the software, we generated two pairs of synthetic datasets from each distribution family, checking if it correctly identified both the family of each pair, and for each pair, which one of the two came from a distribution with higher restricted geometric mean. Each synthetic dataset consisted of 150 points, sampled from the the family, with a sample from the prior for that family determining the parameters.

The posterior probabilities inferred on distribution families, for each synthetic dataset, are shown in Figure 48. Note that we have two pairs for each of 9 families, giving 18 total.

Of the 18 pairs run, by *maximum a posteriori* (MAP) identification (multiple warnings about which are given in Appendix 1):

- one of the two log-Gaussian pairs was identified as log-Student;
- one of the two Student pairs was identified as Gaussian;
- one of the two log-Student pairs was identified as inverse-Gamma;
- one of the two skew-Student pairs was identified as Gaussian;
- one of the two Gamma-power pairs was identified as Gamma.

The remaining 13 pairs of groups were correctly MAP-identified. Of the MAP-misidentified pairs, the lowest probability attached to the correct identification was 0.08; four of the misidentifications were as families with fewer parameters (of which three as subfamilies) and one with more parameters.

As a basic sanity check of the posterior probabilities, we performed the following non-rigorous test:

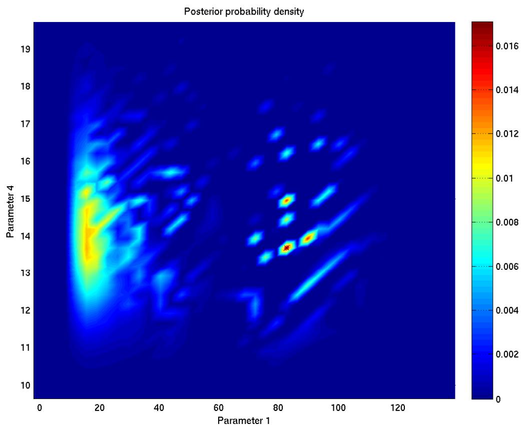

Figure 45: Joint 2-d marginal posterior on parameters $\xi$ and $\eta$ when inferring parameters of a (non-log-) skew-Student model for a log-skew-Student dataset, showing inadequacy of grid resolution. Nonetheless it was clear that the skew-Student could not account for this distribution, as its posterior probability was less than $10^{-48}$ times that of the log-skew-Student. The projection of the MAP parameter vector is marked.

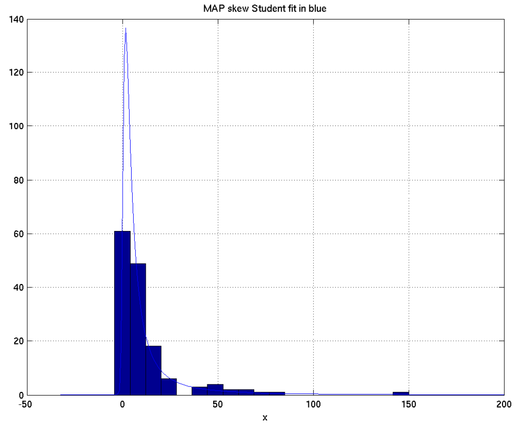

Figure 46: The data that gave rise to Figure 45 and the MAP skew-Student fit on a linear scale; see also Figure 47.

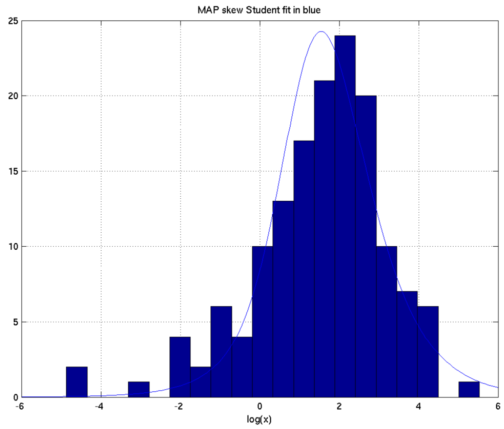

Figure 47: The data that gave rise to Figure 45 and the MAP skew-Student fit on a logarithmic scale, showing how bad the MAP skew-Student fit is to this synthetic log-skew-Student dataset.

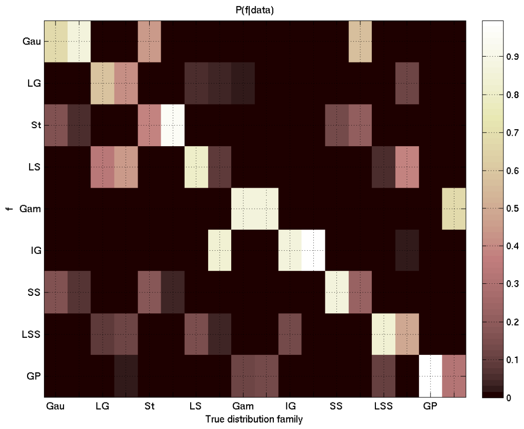

Figure 48: Posterior probabilities of the different possible distribution families for each of the test datasets.

1. Sort the 162 posterior probabilities into order;
2. For each initial segment and final segment of this sequence, count the number that represent the posterior probability of the true family, and apply the (approximate) inverse CDF of the distribution for this that we would expect given the posterior probabilities;
3. Check that all of these quantiles lie between 0.025 and 0.975.

The CDF and thus inverse CDF was approximated by an empirical CDF of 10000 samples from the Bernoulli distribution for each probability.

As seen in Figure 49, which shows these quantiles for each of the 162 possible initial and final segments, we don't observe any outside of our sanity range. If our posterior probabilties had a consistent large bias, or other major problem, we'd expect this simple test to pick up on it.

We observed no bias in favour of distribution families with more parameters, in contrast to maximum likelihood fitting; indeed all but one of the MAP errors observed were in the opposite direction, and all were far from definite misidentifications.

All calculations of $P(Q(f, \theta_{f,A}) > Q(f, \theta_{f,B})|A, B)$ inferred the correct sense of this inequality.

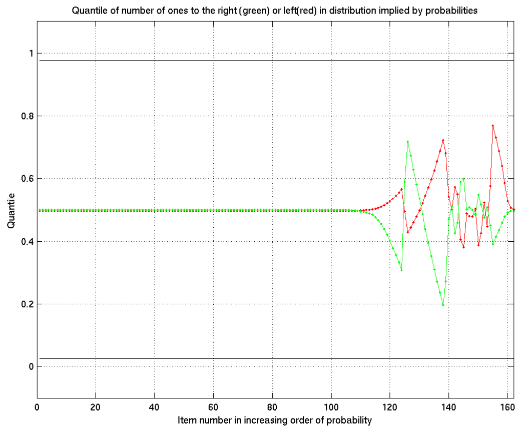

Figure 49: Results of sanity checking of inference software — see text for details.

It is immediately apparent that inference of the family of distributions that generated a particular dataset is based not only on the family definition, but also on the priors on its parameters; e.g. with these priors a dataset all of whose values lie between 100 and 150 is much more likely to be thought to be Gaussian or Student than one whose values all like between 0.001 and 0.1. This is, however, as it should be. As explained e.g. in section 5.2 above, the settings of the parameters are an essential part of practical use of each distribution family.

### 9 Comments on necessary practical details

In order to carry out these calculations, it is necessary to attend to some critical practical details.

#### 9.1 Calculation in the log domain

The probabilities and densities involved frequently lie outside the range of approximately $10^{-300}$ to $10^{300}$ representable in standard floating point arithmetic on a 64-bit computer. In order to ensure that overflow to 0 or $\infty$ does not occur, it is necessary to work entirely in the log-domain, without moving to the linear domain except for viewing the final answers. Thus e.g. one must use the Matlab `gammaln` function to evaluate $\log(\Gamma(m))$ and not e.g. call `log(gamma(m))`, and similarly for ${}_1F_1$.

#### 9.2 Running time and memory requirements

The implementation used takes approximately 3.5 hours to evaluate a single set of three subgroups of a cytokine on a total of around 400 patients, and requires around 20 Gigabytes of memory in its current implementation. Over 95% of the processing time and memory requirement is used in evaluating the log-skew-Student density, and most of that in evaluating the logarithm of the hypergeometric function ${}_1F_1$. That this is worth-while turns out to be apparent from the calculated posterior probability that these cytokines, if all distributed from the same distribution family, are more than $10^{64}$ times more likely to be from the log-skew-Student than from any other of the distribution families considered.

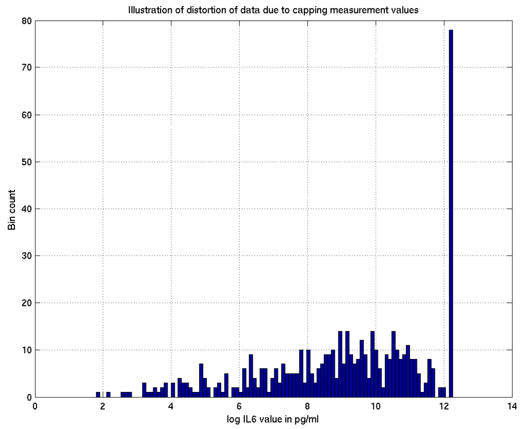

Figure 50: Histogram of samples of IL6, clipped at 210000 pg/ml, distorting the distribution

##### 9.3 Dealing with data at the extreme bounds of the experimental measurement range

Observed distributions can be greatly distorted by forcing all large values to a particular "top of measurement range" value, as illustrated in Figure 50. Similarly this can happen due to finite resolution, and due to clipping values at the bottom of the range.

In order that the results are not distorted by this, we did the following steps, for which process we use the common term "dithering":

1. We first inferred the distribution types as in part 1 of section 2.
2. For each value to be replaced, we:
   (a) Drew a family at random with probability equal to the posterior probability of that family;
   (b) Used the MAP sample of the parameters of that distribution to give a specific distribution from that family;
   (c) Replaced the clipped value with a random sample from that distribution, conditional on being within the range possible given the finite resolution and any clipping and the observed value.

Such random sampling was done by rejection sampling, easy to program though not efficient. We then reran the entire analysis using as input data the resampled clipped values.

(Note that we do not recommend MAP sampling in general; this was merely a quick and dirty expedient which was cheap in compute power and programming effort to execute. In an ideal world we would recommend Markov-chain Monte Carlo to infer the unclipped values, but in this context the workload would be too great. Indeed if we had used a random posterior sample of the parameters rather than the MAP value what we did would be precisely the first step of such an MCMC process.)

#### 10 The probability comparison problem

##### 10.1 Introduction

We now turn to a different question, namely assessment of whether a decision to collect cytokine measurements by the clinicians is influenced by the values of certain other variables. In particular we take

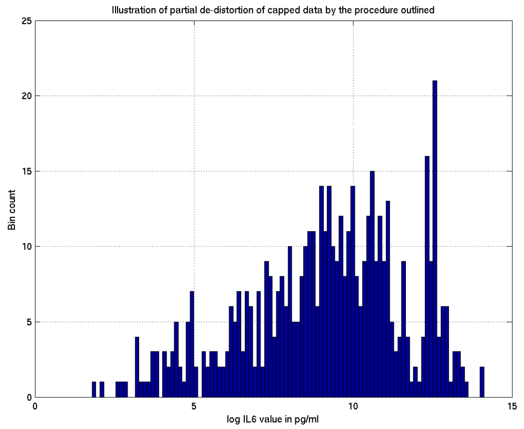

Figure 51: Histogram of samples of IL6, after resampling the clipped values from the MAP inferred distribution. Not perfect, but a great deal better than the unresampled version.

a subset $S$ of the patients, separate them into those without and with a particular variable ($S_0, S_1$ respectively), and we consider the respective probabilities $p_0$ and $p_1$ that a patient from $S_0$ or $S_1$ has their cytokines measured.

As data for this we have the values $N_0$ and $N_1$, the total numbers of patients in $S_0$ and $S_1$, and $n_0$ and $n_1$, the respective numbers who did have their cytokines measured. In the majority of cases a patient either had all or no cytokines measured; for the few for whom this was not true, we took measurement or not of TNF$\alpha$ as the defining datum.

#### 10.2 The questions asked

We then asked two questions:

1. Consider two models for what we think about $p_0, p_1$ before collecting the data, namely $M_1$ in which we are 100% sure that $p_0 = p_1$ and that this common value is uniformly distributed on $[0, 1]$, and $M_2$ in which we believe that $p_0$ and $p_1$ are independently uniformly distributed on $[0, 1]$ (so that the probability that they are exactly equal is zero). Now suppose we think the prior probabilities that $M_1$ holds and that $M_2$ holds are equal (and are both $\frac{1}{2}$). Now, in the light of the data, how likely is it that $M_2$ is the correct model ?

2. Suppose now that $M_2$ is the correct model. How likely is it that $p_0 > p_1$ ?

#### 10.3 Answering the first question

Similarly to the method of model choice used when considering different distribution families, we proceed as follows.

Applying Bayes' theorem we find

$$P(M_2|n_0, n_1, N_0, N_1) = \frac{P(M_2)P(n_0, n_1|M_2, N_0, N_1)}{\sum_{k=1}^{2} P(M_k)P(n_0, n_1|M_k, N_0, N_1)} = \frac{P(n_0, n_1|M_2, N_0, N_1)}{\sum_{k=1}^{2} P(n_0, n_1|M_k, N_0, N_1)}.$$

But now

$$
\begin{aligned}
P(n_0, n_1|M_1, N_0, N_1) &= \int P(n_0, n_1, p|M_1, N_0, N_1)\, dp \\
&= \int P(n_0, n_1|p, M_1, N_0, N_1) P(p|M_1, N_0, N_1)\, dp \\
&= \int P(n_0, n_1|p, M_1, N_0, N_1)\, dp \\
&= \int \frac{N_0!}{n_0!(N_0 - n_0)!} p^{n_0} (1-p)^{N_0 - n_0} \frac{N_1!}{n_1!(N_1 - n_1)!} p^{n_1} (1-p)^{N_1 - n_1}\, dp \\
&= \int \frac{N_0! N_1!}{n_0!(N_0 - n_0)! n_1!(N_1 - n_1)!} p^{n_0 + n_1} (1-p)^{N_0 - n_0 + N_1 - n_1}\, dp \\
&= \frac{N_0! N_1!}{n_0!(N_0 - n_0)! n_1!(N_1 - n_1)!} \frac{\Gamma(n_0 + n_1 + 1)\Gamma(N_0 - n_0 + N_1 - n_1 + 1)}{\Gamma(N_0 + N_1 + 2)}
\end{aligned}
$$

(where $\Gamma(m)$ denotes the Gamma function given by $\Gamma(m) = \int_0^\infty x^{m-1} e^{-x}\, dx$) while

$$
\begin{aligned}
P(n_0, n_1|M_2, N_0, N_1) &= \int P(n_0, n_1, p_0, p_1|M_2, N_0, N_1)\, d(p_0, p_1) \\
&= \int P(n_0, p_0|N_0)\, dp_0 \int P(n_1, p_1|N_1)\, dp_1
\end{aligned}
$$

and for $k = 0, 1$,

$$
\begin{aligned}
\int P(n_k, p_k|N_k)\, dp_k &= \int P(p_k|N_k) P(n_k|p_k, N_k)\, dp_k \\
&= \int P(n_k|p_k, N_k)\, dp_k \\
&= \int \frac{N_k!}{n_k!(N_k - n_k)!} p_k^{n_k} (1 - p_k)^{N_k - n_k}\, dp_k \\
&= \frac{N_k!}{n_k!(N_k - n_k)!} \frac{\Gamma(n_k + 1)\Gamma(N_k - n_k + 1)}{\Gamma(N_k + 2)}.
\end{aligned}
$$

Putting all this together we find that

$$
\begin{aligned}
P(M_2|n_0, n_1, N_0, N_1) &= \frac{\prod_{k=0}^{1} \frac{N_k!}{n_k!(N_k-n_k)!} \frac{\Gamma(n_k+1)\Gamma(N_k-n_k+1)}{\Gamma(N_k+2)}}{\prod_{k=0}^{1} \frac{N_k!}{n_k!(N_k-n_k)!} \frac{\Gamma(n_k+1)\Gamma(N_k-n_k+1)}{\Gamma(N_k+2)} + \frac{N_0! N_1!}{n_0!(N_0-n_0)! n_1!(N_1-n_1)!} \frac{\Gamma(n_0+n_1+1)\Gamma(N_0-n_0+N_1-n_1+1)}{\Gamma(N_0+N_1+2)}} \\
&= \frac{\prod_{k=0}^{1} \frac{\Gamma(n_k+1)\Gamma(N_k-n_k+1)}{\Gamma(N_k+2)}}{\prod_{k=0}^{1} \frac{\Gamma(n_k+1)\Gamma(N_k-n_k+1)}{\Gamma(N_k+2)} + \frac{\Gamma(n_0+n_1+1)\Gamma(N_0-n_0+N_1-n_1+1)}{\Gamma(N_0+N_1+2)}}
\end{aligned}
$$

which can now be easily evaluated e.g. in Matlab.

#### 10.4 Answering the second question

Here we have

$$
\begin{aligned}
P(p_0 > p_1|n_0, n_1, N_0, N_1) &= \int_{\{(p_0,p_1):p_0>p_1\}} P(p_0, p_1|n_0, n_1, N_0, N_1)\, d(p_0, p_1) \\
&= \int_{\{(p_0,p_1):p_0>p_1\}} P(p_0|n_0, N_0) P(p_1|n_1, N_1)\, d(p_0, p_1)
\end{aligned}
$$

while for $k = 0, 1$,

$$
\begin{aligned}
P(p_k|n_k, N_k) &= \frac{P(p_k)P(n_k|p_k, N_k)}{\int_0^1 P(p_k)P(n_k|p_k, N_k)\, dp_k} \\
&= \frac{P(n_k|p_k, N_k)}{\int_0^1 P(n_k|p_k, N_k)\, dp_k} \\
&= \frac{p_k^{n_k}(1-p_k)^{N_k-n_k}}{\int_0^1 p_k^{n_k}(1-p_k)^{N_k-n_k}\, dp_k} \\
&= \frac{\Gamma(N_k+2)}{\Gamma(n_k+1)\Gamma(N_k-n_k+1)} p_k^{n_k}(1-p_k)^{N_k-n_k},
\end{aligned}
$$

following which the required integral can be numerically evaluated on a grid.
