## Supplementary material for "Elevated cerebrospinal fluid cytokine levels in tuberculous meningitis predict survival in response to dexamethasone": Software: ars.m

```
function y = ars(logf, gradlogf, points, lbound, ubound)
% Usage: y = ars(logf, gradlogf, points, lbound, ubound)
% This function does adaptive rejection sampling of a 1-d 
%  log-concave probability density a constant multiple of whose 
%  logarithm is returned by the function logf, and 
%  the gradient of which is returned by gradlogf.
% See https://en.wikipedia.org/wiki/Rejection_sampling#Adaptive_rejection_sampling .
% lbound and ubound are the upper and lower ends of the 
%  distribution range (and may be -Inf and Inf respectively).
% points, a row vector, should ideally give a set of distinct points
%  in the range, at the first of which the gradient is positive
%  and at the second of which the gradient is negative.
% To some extent this function is capable of fixing some 
%  sets of points which don't meet this criterion.

% Change Log:
%
%     1.1          25:may:20    rfs      Taken from survival project ars.m v1.56 .
%     1.68         25:may:20    rfs      Version number harmonisation.

mytitl = ' /home/rfs/ramakrishnan/software/cytokines/rfs/SCCS/s.ars.m 1.102 20/09/03 08:54:05 ';

persistent mytitldone
if isempty(mytitldone),           
   mytitldone = titlfunction(mytitl);
end

global titls

% The variable m will accumulate the amount by which the numbers returned
%  by logf need reducing to avoid overflow on exponentiation.
% vals contains the log densities at points.
% grads contains the gradients of the log density at points.
% bounds contains lbound, the points of intersection of the 
%  tangents fitted to the log density, and ubound, in increasing order.

vals = logf(points);
m = max(vals);
vals = vals - m;
grads = gradlogf(points);

% RFS: There is a serious risk that this calculation will end up giving NaNs, 
%      either because of equal gradients or because of infinite gradients.
%      I suggest waiting to see how much of a problem this is in practice.
bounds = [lbound,((vals(2:end)-vals(1:end-1) + (grads(1:end-1).*points(1:end-1)) - (grads(2:end).*points(2:end)))./(grads(1:end-1) - grads(2:end))), ubound];

% RFS: The following is to make sure that bounds (the intersections of the 
%      pieces of the piecewise exponential 'upper bound') are correctly
%      related to points - necessary because the above calculation can 
%      violate that restriction due to numerical inaccuracy.
bounds(2 : end - 1) = min(points(2 : end), max(points(1 : end - 1), bounds(2 : end - 1)));

% RFS: The following is to check that this function has been correctly called,
%      and if not, to deal with the problem.
boundedlow = ~(isinf(bounds(1)) && grads(1) <= 0);
boundedhigh = ~(isinf(bounds(end)) && grads(end) >= 0);
while ~(boundedlow && boundedhigh),
   if ~boundedlow,
      points = [min(2 * points(1), -1), points];
   end
   if ~boundedhigh,
      points = [points, max(2 * points(end), 1)];
   end
   vals = logf(points) - m;
   maxvals = max(vals);
   m = m + maxvals;
   vals = vals - maxvals;
   grads = gradlogf(points);

   bounds = [lbound,((vals(2:end)-vals(1:end-1) + (grads(1:end-1).*points(1:end-1)) - (grads(2:end).*points(2:end)))./(grads(1:end-1) - grads(2:end))), ubound];

   boundedlow = ~(isinf(bounds(1)) && grads(1) <= 0);
   boundedhigh = ~(isinf(bounds(end)) && grads(end) >= 0);
end   

while true, % Do ARS steps, drawing from the current piecewise exponential fit
            % then if the rejection step rejects, fitting a new tangent and
            % updating bounds.

%    if any(isnan(points)) || any(isnan(vals)) || any(isnan(grads)) || any(isnan(bounds)),
%        keyboard;
%    end

    if length(points) > 2000,
       keyboard;
    end

    Nrejects = 0;
    done = 0;
    while ~done,
       p = piecewise_exponential(points, vals, grads, bounds);
       k = p(1);
       if ismember(k, points),
          Nrejects = Nrejects + 1;
          if Nrejects > 10,
             keyboard; % We shouldn't end up here because piecewise_exponential
                       % contains measures designed to prevent return of its endpoints,
                       % but this is to catch and allow debugging of any failure of 
                       % those measures.
          end
       else
          done = 1;
       end
    end

    u = rand;

    z1 = logf(k) - m;
    maxvals = max([z1, vals]);
    m = m + maxvals;
    vals = vals - maxvals;
    z1 = z1 - maxvals;

    l = exp(z1 - (grads(p(2))*(k-points(p(2))))-vals(p(2)));
    if l > 1.001,
        warning('Function is not log-concave.');
    elseif u < l
        break
    end
    
    if k > points(p(2))
        loc = p(2);
    else
        loc = p(2)-1;
    end
    points = [points(1:loc), k, points(loc+1:end)];

    if any(diff(points) <= 0),
       keyboard;
    end

    vals = [vals(1:loc), z1, vals(loc+1:end)];
    grads = [grads(1:loc), gradlogf(k), grads(loc+1:end)];

    % RFS: Repeat above warning.    
    bounds = [lbound,((vals(2:end)-vals(1:end-1) + (grads(1:end-1).*points(1:end-1)) - (grads(2:end).*points(2:end)))./(grads(1:end-1) - grads(2:end))), ubound];

    % RFS: As before.
    bounds(2 : end - 1) = min(points(2 : end), max(points(1 : end - 1), bounds(2 : end - 1)));

    % RFS: Don't do this, as it easily exceeds recursion limits.
    %     nlogf = @(x) logf(x) - m;
    %     logf = nlogf;

    maxvals = max(vals);
    m = m + maxvals;
    vals = vals - maxvals;
end
y = k;
end

% Local Variables: 
% indent-line-function: indent-relative
% eval: (auto-fill-mode 0)
% End:
```
