## Supplementary material for "Elevated cerebrospinal fluid cytokine levels in tuberculous meningitis predict survival in response to dexamethasone": Software: calccompprobs.m

```
function compprob = calccompprobs(Pxs, xs);
% Usage: compprob = calccompprobs(Pxs, xs);
% Pxs and xs should each be cell arrays of size {2, 1}.
% For each cell, Pxs gives a column vector of probabilities that a random variable 
%  takes the corresponding value in the column vector in the corresponding cell
%  of xs.
% This function then calculates the probability that the value
%  in the first cell is greater than the value in the second cell.
% HOWEVER, where there are equal values, it counts the first cell as 
%  less than the second. The easiest way of fixing this, though it is rather inefficient,
%  is to call it twice giving the cells both ways round, then to normalise.

% Change Log:
%
%     1.1          06:may:20    rfs      First version.
%     1.10         06:may:20    rfs      Version number harmonisation.

mytitl = ' /home/rfs/ramakrishnan/software/cytokines/rfs/SCCS/s.calccompprobs.m 1.102 20/09/03 08:54:05 ';

persistent mytitldone
if isempty(mytitldone),           
   mytitldone = titlfunction(mytitl);
end

global titls

if any(cellfun(@isempty, xs)),
   compprob = 0.5;
   return;
end

x = cat(1, xs{:});
Px = [[Pxs{1}; zeros(size(Pxs{2}))], [zeros(size(Pxs{1})); Pxs{2}]];

if size(x, 2) ~= 1 || size(Px, 2) ~= 2 || size(Px, 1) ~= length(x),
   error('Input arrays are not of the correct sizes');
end

[x, ind] = sort(x, 1, 'ascend');
Px = Px(ind, :);
cumPx = cumsum(Px, 1);

compprob = sum(Px(:, 1) .* cumPx(:, 2));

return;


% Local Variables: 
% indent-line-function: indent-relative
% eval: (auto-fill-mode 0)
% End:
```
