## Supplementary material for "Elevated cerebrospinal fluid cytokine levels in tuberculous meningitis predict survival in response to dexamethasone": Software: checkdither.m

```
function checkdither(cyksold, cyksnew, tcount, hiv, mins, maxs);
% Usage: checkdither(cyksold, cyksnew, tcount, hiv, mins, maxs);
% Compares histograms of cyksold and cyksnew for each combination
%  of ncyk, tcount, and hiv values.
% Here tcount indexes groups starting at 0 and hiv gives subset
%  selection starting at 0.

% Change Log:
%
%     1.1          12:jun:20    rfs      First version.
%     1.89         12:jun:20    rfs      Version number harmonisation; now working.
%     1.98         08:jul:20    rfs      Added mins and maxs and marked their positions.

mytitl = ' /home/rfs/ramakrishnan/software/cytokines/rfs/SCCS/s.checkdither.m 1.102 20/09/03 08:54:05 ';

persistent mytitldone
if isempty(mytitldone),           
   mytitldone = titlfunction(mytitl);
end

global titls

Ncyks = size(cyksold, 2);
Ntcounts = length(unique(tcount));
Nhivs = length(unique(hiv));

for ncyk = 1 : Ncyks,
   for thistcount = 0 : Ntcounts - 1,
      for thishiv = 0 : Nhivs - 1,

         clf;
      
         ind = tcount == thistcount & hiv == thishiv;

         cold = log(cyksold(ind, ncyk));
         cnew = log(cyksnew(ind, ncyk));

         cmin = min([cold; cnew]);
         cmax = max([cold; cnew]);

         Nbins = 30;
         cc = cmin + (cmax - cmin) * [0 : Nbins] / Nbins;

         ho = hist(cold, cc);
         hn = hist(cnew, cc);
         bar(cc, [ho(:), hn(:)]);
         colormap([1, 0, 0; 0, 1, 0]);
         shading('flat');
         title(sprintf('ncyk %d, tcount %d, hiv %d', ncyk, thistcount, thishiv));

         hold on;
         ax = axis;
         plot(log([mins(ncyk), mins(ncyk)]), ax(3:4), 'k');
         plot(log([maxs(ncyk), maxs(ncyk)]), ax(3:4), 'k');
         hold off;

         keyboard;
      end
   end
end

return;


% Local Variables: 
% indent-line-function: indent-relative
% eval: (auto-fill-mode 0)
% End:
```
