## Supplementary material for "Elevated cerebrospinal fluid cytokine levels in tuberculous meningitis predict survival in response to dexamethasone": Software: comparegroups.m

```
function [Etypes, compprobslin, compprobslog, compprobsgeom, ...
          allcompprobslin, allcompprobslog, allcompprobsgeom, Etot] ...
            = comparegroups(xs, priors, xmeangridlin, xmeangridlog, parmstrue, nfig, ...
                            allcompprobslin, allcompprobslog, allcompprobsgeom, Etot);
% Usage: [Etypes, compprobslin, compprobslog, compprobsgeom, ...
%         allcompprobslin, allcompprobslog, allcompprobsgeom, Etot] ...
%           = comparegroups(xs, priors, xmeangridlin, xmeangridlog, parmstrue, nfig, ...
%                           allcompprobslin, allcompprobslog, allcompprobsgeom, Etot);
% This function runs gridassess on each of the groups of data using each of the
%  distribution types given in priors, and reports the posterior probabilities that the 
%  linear and geometric means of the different groups have certain ordering relationships.
% xs should be a cell array of size {Ngroups, 1}, 
%  each cell of which contains a vector of values of x; 
%  the different groups can be of different sizes.
% priors is as output by setpriors.m .
% xmeangridlin is a grid of x values over which the various possible 
%  distributions will have their mean numerically measured with linear weighting;
%  it should be uniformly spaced.
% xmeangridlog is a grid of x values over which the various possible 
%  distributions will have their arithmetic mean numerically measured with a logarithmic grid;
%  log(xmeangridlog) should be uniformly spaced.
% parmstrue, if passed, is as output by makesynthdata.m .
% nfig, if non-empty, gives the first figure number to be used;
%  if empty, plotting is omitted.
% compprobslin(ngroup1, ngroup2) gives the probability that group ngroup1
%  has mean greater than that of group ngroup2 (with equality counting as half),
%  where we mean arithmetic mean calculated on a linear grid.
% compprobslog is analogous for arithmetic mean calculated on a logarithmically spaced grid.
% compprobsgeom is analogous for geometric mean calculated on a logarithmically spaced grid.
% Etypes is minus the log posterior probability that the various groups are all of the relevant type.
% Etot is as Etypes but kept separate for the different groups, of size [Ntypes, Ngroups].
% allcompprobslin is the array of size [Ngroups, Ngroups, Ntypes] found
%  before weighting by the type probabilities.
% If Etot, allcompprobslin, allcompprobslog, and allcompprobsgeom are passed in, then if necessary
%  the Etypes dimension will be extended if necessary, then anything with non-NaN in
%  any of them for a given ntype will be skipped (the input value just being copied to the output)
%  while those with NaN everywhere will be computed from scratch.


% Change Log:
%
%     1.1          06:may:20    rfs      First version.
%     1.10         06:may:20    rfs      Appears to be working.
%     1.11         06:may:20    rfs      Now ignores zero-probability distribution types.
%     1.12         07:may:20    rfs      Comparisons with different numbers of parameters now working.
%     1.13         08:may:20    rfs      Now traps posonly distribution types if data is non-positive.
%     1.31         11:may:20    rfs      Moved setting Ngridmax and shrinklimit to setpriors.m .
%     1.60         16:may:20    rfs      Saves allcompprobslin and allcompprobslog as well.
%     1.67         21:may:20    rfs      Returns Etot also.
%     1.70         25:may:20    rfs      Now allows shortcuts by inputting values precomputed for 
%                                        some of the distribution types.
%     1.85         03:jun:20    rfs      Makes geometric means also.
%     1.86         04:jun:20    rfs      Fixed typos in above.
%     1.97         06:jul:20    rfs      Fixed bug that only matters when there is non-zero probability
%                                        of equality.

mytitl = ' /home/rfs/ramakrishnan/software/cytokines/rfs/SCCS/s.comparegroups.m 1.102 20/09/03 08:54:05 ';

persistent mytitldone
if isempty(mytitldone),           
   mytitldone = titlfunction(mytitl);
end

global titls

if nargin < 4,
   xmeangridlog = [];
end
if nargin < 5,
   parmstrue = [];
end
if nargin < 6,
   nfig = [];
end
if nargin < 7,
   allcompprobslin = [];
end
if nargin < 8,
   allcompprobslog = [];
end
if nargin < 9,
   allcompprobsgeom = [];
end
if nargin < 10,
   Etot = [];
end

Ngroups = length(xs);

if isempty(parmstrue),
   parmstrue = cell(Ngroups, 1);
end

if ~iscell(xs),
   error('xs should be a cell array with one group in each cell');
end

Ntypes = length(priors.typelist);

parmmaxes = cell(Ntypes, Ngroups);
xmeanvalslin = cell(Ntypes, Ngroups);
xmeanvalslog = cell(Ntypes, Ngroups);
xmeanvalsgeom = cell(Ntypes, Ngroups);
Pxmeanvals = cell(Ntypes, Ngroups);

if isempty(Etot),
   Etot = NaN(Ntypes, Ngroups);
end
if isempty(allcompprobslin),
   allcompprobslin = NaN(Ngroups, Ngroups, Ntypes);
end
if isempty(allcompprobslog),
   allcompprobslog = NaN(Ngroups, Ngroups, Ntypes);
end
if isempty(allcompprobsgeom),
   allcompprobsgeom = NaN(Ngroups, Ngroups, Ntypes);
end

if size(Etot, 1) < Ntypes,
   Etot = [Etot; NaN(Ntypes - size(Etot, 1), Ngroups)];
end
if size(allcompprobslin, 3) < Ntypes,
   allcompprobslin = cat(3, allcompprobslin, NaN(Ngroups, Ngroups, Ntypes - size(allcompprobslin, 3)));
end
if size(allcompprobslog, 3) < Ntypes,
   allcompprobslog = cat(3, allcompprobslog, NaN(Ngroups, Ngroups, Ntypes - size(allcompprobslog, 3)));
end
if size(allcompprobsgeom, 3) < Ntypes,
   allcompprobsgeom = cat(3, allcompprobsgeom, NaN(Ngroups, Ngroups, Ntypes - size(allcompprobsgeom, 3)));
end

compprobslin = allcompprobslin;
compprobslog = allcompprobslog;
compprobsgeom = allcompprobsgeom;

% Apply gridassess to each group and distribution type.

for ngroup = 1 : Ngroups,
   for ntype = 1 : Ntypes,

      if (~isnan(Etot(ntype, ngroup)) ...
          && all(~isnan(compprobslin(:, ngroup, ntype))) ...
          && all(~isnan(compprobslin(ngroup, :, ntype))) ...
          && all(~isnan(compprobslog(:, ngroup, ntype))) ...
          && all(~isnan(compprobslog(ngroup, :, ntype))) ...
          && all(~isnan(compprobsgeom(:, ngroup, ntype))) ...
          && all(~isnan(compprobsgeom(ngroup, :, ntype)))),
         continue;
      end

      fprintf('Starting ngroup %d, test distribution type %d\n', ngroup, ntype);

      if priors.Etypes(ntype) < Inf && (all(xs{ngroup} > 0) || ~priors.posonly(ntype)),
   
         fprintf('Running gridassess on ngroup %d, ntype %d\n', ngroup, ntype);
   
         [thisEtot, thisparmmaxes, thisPxmeanvals, thisxmeanvalslin, thisxmeanvalslog, thisxmeanvalsgeom, nfig] ...
               = gridassess(xs{ngroup}, ntype, priors, parmstrue{ngroup}, ...
                            nfig, xmeangridlin, xmeangridlog);
   
         Etot(ntype, ngroup) = thisEtot;
         parmmaxes{ntype, ngroup} = thisparmmaxes;
         Pxmeanvals{ntype, ngroup} = thisPxmeanvals;
         xmeanvalslin{ntype, ngroup} = thisxmeanvalslin;
         xmeanvalslog{ntype, ngroup} = thisxmeanvalslog;
         xmeanvalsgeom{ntype, ngroup} = thisxmeanvalsgeom;

      else % i.e. if this type is not possible

         Etot(ntype, ngroup) = Inf;

      end

   end % for ntype = 1 : Ntypes
end % for ngroup = 1 : Ngroups

Etypes = sum(Etot, 2);
Etypes = Etypes - min(Etypes);
Etypes = Etypes - Esum(Etypes, 1);
Ptypes = exp(-Etypes);
Ptypes = Ptypes / sum(Ptypes);

% Do the comparisons of the means.

for ntype = 1 : Ntypes,
   if priors.Etypes(ntype) < Inf && (all(xs{ngroup} > 0) || ~priors.posonly(ntype)),
      for ngroup1 = 1 : Ngroups,
         for ngroup2 = 1 : Ngroups,

            if isnan(compprobslin(ngroup1, ngroup2, ntype)),
               compprobslin(ngroup1, ngroup2, ntype) ...
                  = calccompprobs([Pxmeanvals(ntype, ngroup1); Pxmeanvals(ntype, ngroup2)], ...
                                  [xmeanvalslin(ntype, ngroup1); xmeanvalslin(ntype, ngroup2)]);
            end

            if isnan(compprobslog(ngroup1, ngroup2, ntype)),
               compprobslog(ngroup1, ngroup2, ntype) ...
                  = calccompprobs([Pxmeanvals(ntype, ngroup1); Pxmeanvals(ntype, ngroup2)], ...
                                  [xmeanvalslog(ntype, ngroup1); xmeanvalslog(ntype, ngroup2)]);
            end

            if isnan(compprobsgeom(ngroup1, ngroup2, ntype)),
               compprobsgeom(ngroup1, ngroup2, ntype) ...
                  = calccompprobs([Pxmeanvals(ntype, ngroup1); Pxmeanvals(ntype, ngroup2)], ...
                                  [xmeanvalsgeom(ntype, ngroup1); xmeanvalsgeom(ntype, ngroup2)]);
            end

         end
      end

      % Compensate for mishandling of equality.
      if 1,
         % This method works whichever way equality is handled and however much is equal.
         compprobslin(:, :, ntype) = (compprobslin(:, :, ntype) + (1 - compprobslin(:, :, ntype).')) / 2;
         compprobslog(:, :, ntype) = (compprobslog(:, :, ntype) + (1 - compprobslog(:, :, ntype).')) / 2;
         compprobsgeom(:, :, ntype) = (compprobsgeom(:, :, ntype) + (1 - compprobsgeom(:, :, ntype).')) / 2;
      else
         % This was the previous way of calculating the compensation.
         % It works only if the probability of equality is zero,
         %  as in theory it always is, except because of finite grid resolution.
         compprobslin(:, :, ntype) = compprobslin(:, :, ntype) ...
                                     ./ (compprobslin(:, :, ntype) + compprobslin(:, :, ntype).');
         compprobslog(:, :, ntype) = compprobslog(:, :, ntype) ...
                                     ./ (compprobslog(:, :, ntype) + compprobslog(:, :, ntype).');
         compprobsgeom(:, :, ntype) = compprobsgeom(:, :, ntype) ...
                                     ./ (compprobsgeom(:, :, ntype) + compprobsgeom(:, :, ntype).');
      end
   else % i.e. if this type is not possible
      compprobslin(:, :, ntype) = 0.5;
      compprobslog(:, :, ntype) = 0.5;
      compprobsgeom(:, :, ntype) = 0.5;
   end
end

% Save all the compprobs before weighting as well.

allcompprobslin = compprobslin;
allcompprobslog = compprobslog;
allcompprobsgeom = compprobsgeom;

Ptypesreshaped = reshape(Ptypes, [1, 1, Ntypes]);
compprobslin = sum(compprobslin .* Ptypesreshaped(ones(Ngroups, 1), ...
                                                  ones(1, Ngroups), ...
                                                  :), 3);
compprobslog = sum(compprobslog .* Ptypesreshaped(ones(Ngroups, 1), ...
                                                  ones(1, Ngroups), ...
                                                  :), 3);
compprobsgeom = sum(compprobsgeom .* Ptypesreshaped(ones(Ngroups, 1), ...
                                                    ones(1, Ngroups), ...
                                                    :), 3);

return;


% Local Variables: 
% indent-line-function: indent-relative
% eval: (auto-fill-mode 0)
% End:
```
