## Supplementary material for "Elevated cerebrospinal fluid cytokine levels in tuberculous meningitis predict survival in response to dexamethasone": Software: ditherdo.m

```
function [cyks] = ditherdo(cyks, mins, maxs, ncyks, tcount, hiv, savesetids, savedir);
% Usage: [cyks] = ditherdo(cyks, mins, maxs, ncyks, tcount, hiv, savesetids, savedir);
% Inputs: cyks is a [Npatients, Ncyks] array of cytokine values,
%  which have not been previously dithered, some of which may
%  be jammed at their upper or lower limits of reading.
% This function dithers the cytokine values that are so jammed,
%  doing so over a wide range, with the dither distributed
%  according to the MAP logskewStudent or Gammapow or other distribution whose
%  parameters are extracted from the figures indexed by 
%  savesetids{1 + hiv} and ncyk, subfigures [familyoffset:familyoffset+familywidth] + tcount * groupstep 
%  for each family.
% Doing this is far from ideal (MAP rather than iterated random sample
%  (i.e. MCMC), but a great deal better than using the undithered values.
% All others are dithered log-uniformly over original value +/- 5e-3 .
% Lower limit of reading is assumed to be 1e-2,
%  and upper limit of reading is assumed to be the maximum value of 
%  the relevant cytokine observed.
% ditherdist should be the string representing the distribution type to be used for dithering.
% As the only most likely distributions observed are log-skew-Student and Gammapow, only
%  these have been implemented.

% Change Log:
%
%     1.1          19:may:20    rfs      First version.
%     1.63         19:may:20    rfs      Appears to be working.
%     1.65         21:may:20    rfs      Prevented going below 1e-30 to avoid underflows with inverse Gamma.
%     1.83         30:may:20    rfs      Added Gammapow dithering option.
%     1.88         12:jun:20    rfs      Made to follow posterior probabilities of families;
%                                        before splitting off into individual functions or debugging.
%     1.89         12:jun:20    rfs      Fixed new NaNs appearing; now working, just needs 
%                                        subfunctions moving to their own files.
%     1.90         12:jun:20    rfs      Subfunctions now all in their own files.
%     1.91         19:jun:20    rfs      Incorporating lower bounds from Thuong 2017 supplementary.
%     1.94         28:jun:20    rfs      Introduced absmax to exclude silly large values.

mytitl = ' /home/rfs/ramakrishnan/software/cytokines/rfs/SCCS/s.ditherdo.m 1.102 20/09/03 08:54:05 ';

persistent mytitldone
if isempty(mytitldone),           
   mytitldone = titlfunction(mytitl);
end

global titls

Nncyks = length(ncyks);
Nhivs = length(unique(hiv));
Ntcounts = length(unique(tcount));

absmin = 1e-30;
absmax = 2e6;
Ntrieslong = 1e5;
Ntriesshort = 1e4;

priors = setpriors(1); % Note we use nothing dependent on scale here.

Ntypes = length(priors.typelist);
groupstep = priors.groupstep;
familyoffsets = priors.familyoffsets;
Nparams = priors.Nparams;

for nncyk = 1 : Nncyks,
   ncyk = ncyks(nncyk);

   thismax = maxs(ncyk);
   thismin = mins(ncyk);

   for thistcount = 0 : Ntcounts - 1,
      for thishiv = 0 : Nhivs - 1,

         fprintf('Starting ncyk %d, tcount %d, hiv %d:\n', ncyk, thistcount, thishiv);

         % Collect the MAP parameters for each distribution type.
         mapparms = cell(Ntypes, 1);
         for ntype = 1 : Ntypes,

            thisparams = NaN(Nparams(ntype), 1);

            % Set up the figures from which to extract the MAP parameters
            %  for this distribution type.
            savesetid = [savesetids{thishiv + 1}, '.', sprintf('%d', ncyk)];
            nfig = familyoffsets(ntype) + thistcount * groupstep;

            firstfig = sprintf('%s/Fig%s.fig%d.fig', savedir, savesetid, nfig);
         
            if ~exist(firstfig, 'file'),
               % Then this distribution type was excluded in the priors.
               continue;
            end

            hgload(firstfig);
            ch = get(gca, 'Children');
            thisparams(1) = get(ch(1), 'XData');

            for nparam = 2 : Nparams(ntype),
         
               hgload(sprintf('%s/Fig%s.fig%d.fig', savedir, savesetid, nfig + nparam - 2));
               ch = get(gca, 'Children');
               thisparams(nparam) = get(ch(1), 'YData');

            end % for nparam = 2 : Nparams
         
            close all

            mapparms{ntype} = thisparams;
         end % for ntype = 1 : Ntypes

         % Collect the posterior probabilities of each distribution type.
         load(sprintf('%s/save.%s.mat', savedir, savesetids{thishiv + 1}), ...
              'Etypes');
         Etypes = Etypes(:, ncyk);
         Etypes = Etypes - min(Etypes);
         Etypes = Etypes - Esum(Etypes, 1);

         lowtop = thismax - 5e-3;
         highbot = thismin + 5e-3;
            
         ind = ~isnan(cyks(:, ncyk)) ...
               & tcount == thistcount ...
               & hiv == thishiv;
            
         cold = cyks(ind, ncyk);
         cnew = cold;
                          
         fprintf('ncyk = %d, thistcount = %d, thishiv = %d:\n', ncyk, thistcount, thishiv);
         for n = 1 : length(cnew),
            thisc = cnew(n);
            oldc = cold(n);
            
            if thisc >= thismax,
               thisc = dithersample(priors, lowtop, absmax, Etypes, mapparms, Ntrieslong);
            elseif thisc <= thismin,
               thisc = dithersample(priors, absmin, highbot, Etypes, mapparms, Ntrieslong);
            else
               thisc = dithersample(priors, thisc - 5e-3, thisc + 5e-3, Etypes, mapparms, Ntriesshort);
            end

            % Catchall in case it took too long to find a non-rejected sample,
            % in which case draw log-uniformly from the desired range.
            if isnan(thisc),
               thisc = exp(log(oldc - 5e-3) + (log(oldc + 5e-3) - log(oldc - 5e-3)) * rand(1, 1));
            end
            
            if isnan(thisc) & ~isnan(cold(n)),
               fprintf('New NaN appeared !\n');
               keyboard;
            end

            cnew(n) = thisc;
         end % for n = 1 : length(cnew)
         
         cyks(ind, ncyk) = cnew;
      end
   end
end

return;


function out = dithersample(priors, low, high, Etypes, mapparms, Ntries);
% Usage: out = dithersample(priors, low, high, Etypes, mapparms, Ntries);
% Draws a single sample from a distribution family indicated by the probabilities represented 
%  by Etypes at the MAP parameters in the corresponding entry of mapparms.
% Uses rejection sampling, and if Ntries doesn't get a sample, returns NaN.

Ntypes = length(mapparms);

ntype = randdiscreteE([1, 1], Etypes);

ditherfns = priors.ditherfns;

parms = num2cell(mapparms{ntype});

out = feval(ditherfns{ntype}, low, high, Ntries, parms{:});

return;


% Local Variables: 
% indent-line-function: indent-relative
% eval: (auto-fill-mode 0)
% End:
```
