## Supplementary material for "Elevated cerebrospinal fluid cytokine levels in tuberculous meningitis predict survival in response to dexamethasone": Software: dithergau.m

```
function out = dithergau(low, high, Ntries, mu, s);
% Usage: out = dithergau(low, high, Ntries, mu, s);
% Draws a sample from the Gaussian distribution with the given parameters,
%  conditional on it being between low and high.
% Does this by the dumbest least efficient way available to save programming time,
%  i.e. it draws random samples until it finds one in the given range.

% Change Log:
%
%     1.1          12:jun:20    rfs      First version.

mytitl = ' /home/rfs/ramakrishnan/software/cytokines/rfs/SCCS/s.dithergau.m 1.102 20/09/03 08:54:05 ';

persistent mytitldone
if isempty(mytitldone),           
   mytitldone = titlfunction(mytitl);
end

global titls

ntried = 0;
fprintf('low = %g, high = %g, mu = %g, s = %g:\n', low, high, mu, s);
found = 0;
while ~found && ntried < Ntries,

   out = mu + randn(1, 1) ./ sqrt(s);

   if low <= out && out <= high,
      found = 1;
   end

   ntried = ntried + 1;
   fprintf('\rTried %d times ...         ', ntried);
end
if found,
   fprintf('...found, out = %g                 \n', out);
else
   fprintf('...not found, resorting to log-uniform.\n');
   out = NaN;
end

return;


% Local Variables: 
% indent-line-function: indent-relative
% eval: (auto-fill-mode 0)
% End:
```
