## Supplementary material for "Elevated cerebrospinal fluid cytokine levels in tuberculous meningitis predict survival in response to dexamethasone": Software: Eadd.m

```
function [y] = Eadd(x1, x2);
% Usage: [y] = Eadd(x1, x2);
% This function calculates
%  y = -log(exp(-x1) + exp(-x2))
% without ever exponentiating either -x1 or -x2.

% Change Log:
%
%     1.1          04:jun:20    rfs      First version.
%     1.86         04:jun:20    rfs      Version number harmonisation.

mytitl = ' /home/rfs/ramakrishnan/software/cytokines/rfs/SCCS/s.Eadd.m 1.102 20/09/03 08:54:05 ';

persistent mytitldone
if isempty(mytitldone),           
   mytitldone = titlfunction(mytitl);
end

global titls

sz1 = size(x1);
sz2 = size(x2);

if length(sz1) ~= length(sz2),
   error('Eadd called with arrays of different dimensionality');
end

if any(sz1 ~= sz2),
   error('Eadd called with unequal sizes');
end

y = zeros(sz1);

ind1 = find(x1 < x2);
y(ind1) = x1(ind1) - log(1 + exp(x1(ind1) - x2(ind1)));
ind2 = find(x1 >= x2);
y(ind2) = x2(ind2) - log(1 + exp(x2(ind2) - x1(ind2)));
ind3 = find(x1 == x2 & isinf(x1));
y(ind3) = x1(ind3);

return;


% Local Variables: 
% indent-line-function: indent-relative
% eval: (auto-fill-mode 0)
% End:
```
