## Supplementary material for "Elevated cerebrospinal fluid cytokine levels in tuberculous meningitis predict survival in response to dexamethasone": Software: Egamma.m

```
function E = Egamma(parmgrid, priors, nowarn);
% Usage: E = Egamma(parmgrid, priors, nowarn);
% E is minus log probability density of seeing each of the 1-d variables 
%  in the array x = parmgrid{3} at the parameters m = parmgrid{1}, s = parmgrid{2}, r = m * s,
%  each of which is an array of the same dimension, using the prior parameters
%  in priors.

% Change Log:
%
%     1.1          07:may:20    rfs      First version.

mytitl = ' /home/rfs/ramakrishnan/software/cytokines/rfs/SCCS/s.Egamma.m 1.102 20/09/03 08:54:05 ';

persistent mytitldone
if isempty(mytitldone),           
   mytitldone = titlfunction(mytitl);
end

global titls

persistent warned

if isempty(warned),
   warned = 0;
end

if nargin < 3,
   nowarn = [];
end

if isempty(nowarn),
   nowarn = 0;
end

if isempty(priors) && warned == 0 && ~nowarn,
   warning('Have passed empty priors to Egamma');
   warned = 1;
end

Nparams = 2;

if length(parmgrid) ~= Nparams + 1;
   error('Wrong number of cells in parmgrid in Egamma');
end

m = parmgrid{1};
s = parmgrid{2};
x = parmgrid{3};
r = m .* s;

Elike = - m .* log(r) + gammaln(m) - (m - 1) .* log(x) + r .* x;

if ~isempty(priors),
   prior = priors.Gamma;
   ms = prior.ms;
   ss = prior.ss;
   rs = ms .* ss;
   am = prior.am;
   bm = prior.bm;
   Km = prior.Km;

   sz = size(m);
   szwanted = sz(1 : 2);
   m = reshape(m(1 : prod(szwanted)), szwanted);
   s = reshape(s(1 : prod(szwanted)), szwanted);

   Epriors = - ms * log(rs) + gammaln(ms) - (ms - 1) * log(s) + rs * s;
   Epriorm = - log(Km) + (am + bm) .* m - bm .* m .* log(m) + bm .* gammaln(m);

   Eprior = Epriors + Epriorm;

   E = cat(Nparams + 1, Elike, Eprior);
else
   E = Elike;
end

return;


% Local Variables: 
% indent-line-function: indent-relative
% eval: (auto-fill-mode 0)
% End:
```
