## Supplementary material for "Elevated cerebrospinal fluid cytokine levels in tuberculous meningitis predict survival in response to dexamethasone": Software: Egammapow.m

```
function E = Egammapow(parmgrid, priors, nowarn);
% Usage: E = Egammapow(parmgrid, priors, nowarn);
% E is minus log probability density of seeing each of the 1-d variables 
%  in the array x = parmgrid{3} at the parameters m = parmgrid{1}, 
%  s = parmgrid{2}, k = parmgrid{3}, r = m * s ^ k,
%  each of which is an array of the same dimension, 
%  using the prior parameters in priors,
%  where the density is such that x^k is distributed Gamma(m,r).

% Change Log:
%
%     1.1          07:may:20    rfs      First version.
%     1.69         25:may:20    rfs      Normalisation constant remembered; two silly typos fixed.

mytitl = ' /home/rfs/ramakrishnan/software/cytokines/rfs/SCCS/s.Egammapow.m 1.102 20/09/03 08:54:05 ';

persistent mytitldone
if isempty(mytitldone),           
   mytitldone = titlfunction(mytitl);
end

global titls

persistent warned

if isempty(warned),
   warned = 0;
end

if nargin < 3,
   nowarn = [];
end

if isempty(nowarn),
   nowarn = 0;
end

if isempty(priors) && warned == 0 && ~nowarn,
   warning('Have passed empty priors to Egammapow');
   warned = 1;
end

Nparams = 3;

if length(parmgrid) ~= Nparams + 1;
   error('Wrong number of cells in parmgrid in Egamma');
end

m = parmgrid{1};
s = parmgrid{2};
k = parmgrid{3};
x = parmgrid{4};
r = m .* s .^ k;

Elike = -log(k) - m .* log(r) + gammaln(m) - (m .* k - 1) .* log(x) + r .* x .^ k;

if ~isempty(priors),
   prior = priors.Gammapow;
   ms = prior.ms;
   ss = prior.ss;
   rs = ms .* ss;
   am = prior.am;
   bm = prior.bm;
   Km = prior.Km;
   Nk = prior.Nk;
   ak = prior.ak;
   bk = prior.bk;
   ck = prior.ck;
   Kk = prior.Kk;

   sz = [size(m), ones(1, Nparams)];
   szwanted = sz(1 : Nparams);
   sz = sz(1 : Nparams + 1);
   m = reshape(m(1 : prod(szwanted)), szwanted);
   s = reshape(s(1 : prod(szwanted)), szwanted);
   k = reshape(k(1 : prod(szwanted)), szwanted);

   Epriors = - ms * log(rs) + gammaln(ms) - (ms - 1) * log(s) + rs * s;
   Epriorm = - log(Km) + (am + bm) .* m - bm .* m .* log(m) + bm .* gammaln(m);
   Epriork = - log(Kk) - ak * log(k);
   for n = 1 : Nk,
      Epriork = Epriork - k .* ck(n) .* log(bk(n)) + ck(n) .* bk(n) .^ k;
   end

   Eprior = Epriors + Epriorm + Epriork;

   E = cat(Nparams + 1, Elike, Eprior);
else
   E = Elike;
end

return;


% Local Variables: 
% indent-line-function: indent-relative
% eval: (auto-fill-mode 0)
% End:
```
