## Supplementary material for "Elevated cerebrospinal fluid cytokine levels in tuberculous meningitis predict survival in response to dexamethasone": Software: Egauss.m

```
function E = Egauss(parmgrid, priors, nowarn);
% Usage: E = Egauss(parmgrid, priors, nowarn);
% E is minus log probability density of seeing each of the 1-d variables 
%  in the array x = parmgrid{3} at the parameters mu = parmgrid{1,} s parmgrid{2},
%  each of which is an array of the same dimension, using the prior parameters
%  in priors.

% Change Log:
%
%     1.1          01:may:20    rfs      First version, with priors not yet implemented.
%     1.2          01:may:20    rfs      Fixed forgetting to subtract mean.
%     1.3          01:may:20    rfs      Now handles priors.

mytitl = ' /home/rfs/ramakrishnan/software/cytokines/rfs/SCCS/s.Egauss.m 1.102 20/09/03 08:54:05 ';

persistent mytitldone
if isempty(mytitldone),           
   mytitldone = titlfunction(mytitl);
end

global titls

persistent warned

if isempty(warned),
   warned = 0;
end

if nargin < 3,
   nowarn = [];
end

if isempty(nowarn),
   nowarn = 0;
end

if isempty(priors) && warned == 0 && ~nowarn,
   warning('Have passed empty priors to Egauss');
   warned = 1;
end

Nparams = 2;

if length(parmgrid) ~= Nparams + 1;
   error('Wrong number of cells in parmgrid in Egauss');
end

mu = parmgrid{1};
s = parmgrid{2};
x = parmgrid{3};

Elike = - 0.5 * log(s / (2 * pi)) + 0.5 * s .* (x - mu) .^ 2;

if ~isempty(priors),
   prior = priors.Gaussian;
   ms = prior.ms;
   ss = prior.ss;
   mumu = prior.mumu;
   Kmu = prior.Kmu;

   sz = size(mu);
   szwanted = sz(1 : 2);
   mu = reshape(mu(1 : prod(szwanted)), szwanted);
   s = reshape(s(1 : prod(szwanted)), szwanted);

   Eprior = - ms * log(ms * ss) + gammaln(ms) - (ms - 1) * log(s) + ms * ss * s ...
            - 0.5 * log(Kmu * s / (2 * pi)) + 0.5 * Kmu * s .* (mu - mumu) .^ 2;

   E = cat(Nparams + 1, Elike, Eprior);
else
   E = Elike;
end

return;


% Local Variables: 
% indent-line-function: indent-relative
% eval: (auto-fill-mode 0)
% End:
```
