## Supplementary material for "Elevated cerebrospinal fluid cytokine levels in tuberculous meningitis predict survival in response to dexamethasone": Software: Elogskewstudent.m

```
function E = Elogskewstudent(parmgrid, priors, nowarn);
% Usage: E = Elogskewstudent(parmgrid, priors, nowarn);
% E is minus log probability density of seeing each of the 1-d variables 
%  in the array x = parmgrid{5} at the parameters m = parmgrid{3}, s = parmgrid{2}, 
%  xi = parmgrid{1}, nu = nutrans(parmgrid{4}), 
%  each of which is an array of the same dimension, 
%  where r = m * s, nu = nutrans(nucr), and mu = xi - nu * sqrt(r) * Gamma(m - 0.5) / Gamma(m), and
%  P(x) = (1/x) * (r^m / Gamma(m)) * sqrt(1 / (2 * pi)) * (r + 0.5 * (log(x) - mu) .^ 2) ^ -(m + 0.5) ...
%         * exp(-0.5 * nu ^ 2) * ...
%         * (Gamma(m + 0.5) * 1F1(m + 0.5, 0.5, ((log(x) - mu)' * nu) ^ 2 / (4 * (r + 0.5 * (log(x) - mu) ^ 2))) ...
%            + (((log(x) - mu)' * nu / sqrt(r + 0.5 * (log(x) - mu) .^ 2)) ...
%               * Gamma(m + 1) * 1F1(m + 1, 1.5, ((log(x) - mu)' * nu) .^ 2 / (4 * (r + 0.5 * (log(x) - mu) ^ 2)))))
%   where 1F1 is the hypergeometric function hypergeom, 
% using the prior parameters in priors.
% Note that we must have m > 0.5 .

% Change Log:
%
%     1.1          07:may:20    rfs      First version.
%     1.17         07:may:20    rfs      Version number harmonisation.
%     1.27         09:may:20    rfs      Trapped NaNs and Infs (not ideal - we really need a loghypergeom function).
%     1.41         12:may:20    rfs      Now uses loghypergeom1F1.m .
%     1.46         13:may:20    rfs      Reparameterised using nucr.
%     1.47         13:may:20    rfs      Trivial bug fixed.
%     1.48         13:may:20    rfs      Remembered abs before log.
%     1.49         13:may:20    rfs      Reparameterised nu again.
%     1.51         14:may:20    rfs      Switched to v2 of loghypergeom1F1.
%     1.54         15:may:20    rfs      Fixed horrid bug omitting snu.
%     1.78         29:may:20    rfs      Adjusted recursion to reduce memory consumption.

mytitl = ' /home/rfs/ramakrishnan/software/cytokines/rfs/SCCS/s.Elogskewstudent.m 1.102 20/09/03 08:54:05 ';

persistent mytitldone
if isempty(mytitldone),           
   mytitldone = titlfunction(mytitl);
end

global titls

persistent warned

if isempty(warned),
   warned = 0;
end

if nargin < 3,
   nowarn = [];
end

if isempty(nowarn),
   nowarn = 0;
end

if isempty(priors) && warned == 0 && ~nowarn,
   warning('Have passed empty priors to Elogskewstudent');
   warned = 1;
end

Nparams = 4;

if length(parmgrid) ~= Nparams + 1;
   error('Wrong number of cells in parmgrid in Elogskewstudent');
end

xi = parmgrid{1};
m = parmgrid{2};
s = parmgrid{3};
nucr = parmgrid{4};
x = parmgrid{5};

nu = nutrans(nucr);

% To reduce total memory requirement, we now consider whether to split
%  the evaluation along the x dimension.
if size(x, Nparams + 1) > 1 && size(x, 1) > 5,

   % In this case we evaluate for one nucr value and one x value at a time,
   % by recursively calling this same function on a subarray.

   Elike = NaN(size(x));
   Nx = size(x, Nparams + 1) * size(x, Nparams);
   for nx = 1 : Nx,
      Elike(:, :, :, nx) = Elogskewstudent({xi(:, :, :, nx), ...
                                            m(:, :, :, nx), ...
                                            s(:, :, :, nx), ...
                                            nucr(:, :, :, nx), ...
                                            x(:, :, :, nx)}, ...
                                           [], 1);
   end % for nx = 1 : Nx

else % i.e. if size(x, Nparams + 1) = 1 or size(x, 1) <= 5
   
   r = m .* s;
   mu = xi - nu .* sqrt(r) .* exp(gammaln(m - 0.5) - gammaln(m));
   
   y = log(x) - mu;
   z = r + 0.5 * y .^ 2;
   w = y .* nu;
   
   Eterm1 = - gammaln(m + 0.5) - loghypergeom1F1v2(m + 0.5, 0.5, w .^ 2 ./ (4 * z));
   
   Eterm2 = - log(abs(w)) + 0.5 * log(z) - gammaln(m + 1) - loghypergeom1F1v2(m + 1, 1.5, w .^ 2 ./ (4 * z));
   
   ind = w > 0;
   Eterms = NaN(size(Eterm2));
   Eterms(ind) = Eadd(Eterm1(ind), Eterm2(ind));
   
   Eterm1neg = Eterm1(~ind);
   Eterm2neg = Eterm2(~ind);
   Eterm2neg = max(Eterm2neg, Eterm1neg); % Prevent negative results
   Eterms(~ind) = Esub(Eterm1neg, Eterm2neg);
   
   % Trap NaN values arising from overflow of hypergeom.
   Eterms(isnan(Eterms)) = Inf;
   
   % Trap -Inf values arising from overflow of hypergeom.
   % This is obviously not ideal.
   Eterms(isinf(Eterms)) = Inf;
   
   Elike = (- m .* log(r) + gammaln(m)) + 0.5 * log(2 * pi) ...
           + (m + 0.5) .* log(z) + 0.5 * nu .^ 2 ...
           + Eterms + log(x);

end % if size(x, Nparams + 1) > 1 and size(x, 1) > 5 or not

if ~isempty(priors),
   prior = priors.logskewStudent;
   ms = prior.ms;
   ss = prior.ss;
   rs = ms * ss;
   muxi = prior.muxi;
   sxi = prior.sxi;
   am = prior.am;
   bm = prior.bm;
   Km = prior.Km;
   snu = prior.snu;

   sz = [size(m), ones(1, Nparams)];
   szwanted = sz(1 : Nparams);
   sz = sz(1 : Nparams + 1);
   xi = reshape(xi(1 : prod(szwanted)), szwanted);
   s = reshape(s(1 : prod(szwanted)), szwanted);
   m = reshape(m(1 : prod(szwanted)), szwanted);
   nu = reshape(nu(1 : prod(szwanted)), szwanted);
   nucr = reshape(nucr(1 : prod(szwanted)), szwanted);

   Epriorxi = -0.5 * log(sxi / (2 * pi)) + 0.5 * sxi * (xi - muxi) .^ 2;
   Epriors = - ms * log(rs) + gammaln(ms) - (ms - 1) * log(s) + rs * s;
   Epriorm = - log(Km) + (am + bm) .* m - bm .* m .* log(m) + bm .* gammaln(m);
   Epriornucr = -0.5 * log(snu / (2 * pi)) + 0.5 * snu .* nu .^ 2 - log(nutransderiv(nucr));

   Eprior = Epriorm + Epriorxi + Epriors + Epriornucr;

   E = cat(Nparams + 1, Elike, Eprior);
else
   E = Elike;
end

return;


% Local Variables: 
% indent-line-function: indent-relative
% eval: (auto-fill-mode 0)
% End:
```
