## Supplementary material for "Elevated cerebrospinal fluid cytokine levels in tuberculous meningitis predict survival in response to dexamethasone": Software: Elogstudent.m

```
function E = Elogstudent(parmgrid, priors, nowarn);
% Usage: E = Elogstudent(parmgrid, priors, nowarn);
% E is minus log probability density of seeing each of the 1-d variables 
%  in the array x = parmgrid{4} at the parameters m = parmgrid{3}, s = parmgrid{2}, mu = parmgrid{1},
%  each of which is an array of the same dimension, 
%  where r = m * s, and 
%  P(x) = (1/sqrt(2 * pi)) * (gamma(m + 0.5) / gamma(m)) * r ^ m / (r + 0.5 * (log(x) - mu) ^ 2) ^ (m + 0.5) / x.
% using the prior parameters in priors.

% Change Log:
%
%     1.1          06:may:20    rfs      First version.
%     1.11         06:may:20    rfs      Version number harmonisation.

mytitl = ' /home/rfs/ramakrishnan/software/cytokines/rfs/SCCS/s.Elogstudent.m 1.102 20/09/03 08:54:05 ';

persistent mytitldone
if isempty(mytitldone),           
   mytitldone = titlfunction(mytitl);
end

global titls

persistent warned

if isempty(warned),
   warned = 0;
end

if nargin < 3,
   nowarn = [];
end

if isempty(nowarn),
   nowarn = 0;
end

if isempty(priors) && warned == 0 && ~nowarn,
   warning('Have passed empty priors to Elogstudent');
   warned = 1;
end

Nparams = 3;

if length(parmgrid) ~= Nparams + 1;
   error('Wrong number of cells in parmgrid in Elogstudent');
end

mu = parmgrid{1};
m = parmgrid{2};
s = parmgrid{3};
x = parmgrid{4};

r = m .* s;

Elike = 0.5 * log(2 * pi) - gammaln(m + 0.5) + gammaln(m) - m .* log(r) ...
        + (m + 0.5) .* log(r + 0.5 * (log(x) - mu) .^ 2) + log(x);

if ~isempty(priors),
   prior = priors.logStudent;
   ms = prior.ms;
   ss = prior.ss;
   rs = ms * ss;
   mumu = prior.mumu;
   smu = prior.smu;
   am = prior.am;
   bm = prior.bm;
   Km = prior.Km;

   sz = [size(mu), ones(1, Nparams)];
   szwanted = sz(1 : Nparams);
   sz = sz(1 : Nparams + 1);
   mu = reshape(mu(1 : prod(szwanted)), szwanted);
   s = reshape(s(1 : prod(szwanted)), szwanted);
   m = reshape(m(1 : prod(szwanted)), szwanted);

   Epriormu = -0.5 * log(smu / (2 * pi)) + 0.5 * smu * (mu - mumu) .^ 2;
   Epriors = - ms * log(rs) + gammaln(ms) - (ms - 1) * log(s) + rs * s;
   Epriorm = - log(Km) + (am + bm) .* m - bm .* m .* log(m) + bm .* gammaln(m);

   Eprior = Epriorm + Epriormu + Epriors;

   E = cat(Nparams + 1, Elike, Eprior);
else
   E = Elike;
end

return;


% Local Variables: 
% indent-line-function: indent-relative
% eval: (auto-fill-mode 0)
% End:
```
