## Supplementary material for "Elevated cerebrospinal fluid cytokine levels in tuberculous meningitis predict survival in response to dexamethasone": Software: Esub.m

```
function [y] = Esub(x1, x2);
% Usage: [y] = Esub(x1, x2);
% This function calculates
%  y = -log(exp(-x1) - exp(-x2))
% without ever exponentiating -x1 or -x2.
% It is required that all elements of x1 are less than or equal to the
%  corresponding elements of x2.

% Change Log:
%
%     1.1          04:jun:20    rfs      First version.
%     1.87         04:jun:20    rfs      Version number harmonisation.

mytitl = ' /home/rfs/ramakrishnan/software/cytokines/rfs/SCCS/s.Esub.m 1.102 20/09/03 08:54:05 ';

persistent mytitldone
if isempty(mytitldone),           
   mytitldone = titlfunction(mytitl);
end

global titls

sz1 = size(x1);
sz2 = size(x2);

if length(sz1) ~= length(sz2),
   error('Esub called with arrays of different dimensionality');
end

if any(sz1 ~= sz2),
   error('Esub called with unequal sizes');
end

if any(x1(:) > x2(:)),
   error('Esub called with some of first argument greater than corresponding elements of second argument');
end

y = zeros(size(x1));

ind1 = find(x1 < x2);
y(ind1) = x1(ind1) - log(1 - exp(x1(ind1) - x2(ind1)));
ind3 = find(x1 == x2);
y(ind3) = Inf;

return;


% Local Variables: 
% indent-line-function: indent-relative
% eval: (auto-fill-mode 0)
% End:
```
