## Supplementary material for "Elevated cerebrospinal fluid cytokine levels in tuberculous meningitis predict survival in response to dexamethasone": Software: Esum.m

```
function [y] = Esum(x, n);
% Usage: [y] = Esum(x, n);
% This function calculates
%  y = -log(sum(exp(-x), n))
% without every exponentiating -x .

% Change Log:
%
%     1.1          04:jun:20    rfs      First version.
%     1.86         04:jun:20    rfs      Version number harmonisation.

mytitl = ' /home/rfs/ramakrishnan/software/cytokines/rfs/SCCS/s.Esum.m 1.102 20/09/03 08:54:05 ';

persistent mytitldone
if isempty(mytitldone),           
   mytitldone = titlfunction(mytitl);
end

global titls

shift = min(x, [], n);
ind = find(isinf(shift) | isnan(shift));
shift(ind) = 0;

sz = size(x);
sz = [sz, ones(n - length(sz))];
repper = ones(size(sz));
repper(n) = sz(n);

x = x - repmat(shift, repper);

y = -lognw(sum(exp(-x), n));

y = y + shift;

return;


% Local Variables: 
% indent-line-function: indent-relative
% eval: (auto-fill-mode 0)
% End:
```
