## Supplementary material for "Elevated cerebrospinal fluid cytokine levels in tuberculous meningitis predict survival in response to dexamethasone": Software: gridassess.m

```
function [Etot, parmmaxes, Pxmeanvals, xmeanvalslin, xmeanvalslog, xmeanvalsgeom, nfig] ...
            = gridassess(x, ntype, priors, parmstrue, nfig, xmeangridlin, xmeangridlog);
% Usage: [Etot, parmmaxes, Pxmeanvals, xmeanvalslin, xmeanvalslog, xmeanvalsgeom, nfig] ...
%           = gridassess(x, ntype, priors, parmstrue, nfig, xmeangridlin, xmeangridlog);
% This function takes some data x, 
%  maximum number of grid points to be used priors.Ngridmax,
%  the index number of the distribution type to be assessed as defined by setpriors.m,
%  and a structure priors of parameters for the priors relevant to fn.
% priors contains a field Efns containing a list of function handles corresponding
%  to distribution types, which are called using
%   E = feval(fn, parmgrid, priors, nowarn)
%  where parmgrid is a cell array of values of the parameters,
%   of size {Nparams + 1, 1}, of which cell nparam is the ndgrid output
%   corresponding to parameter nparam, and cell nparam+1 is the ndgrid
%   output corresponding to x,
%  and priors is the set of prior parameters for that evaluation function,
%  and E is minus the log probability density of x at those parameters,
%   of same size as parmgrid{1}, except that the last dimension is one longer, with Eprior being last,
%  and nowarn should be 1 to avoid a warning if passing empty priors.
% It then experiments to find an appropriate grid over which to evaluate the density,
%  shrinking a range until the density at the ends is roughly shrinklimit times the 
%  maximum, then increasing the grid resolution.
% It outputs Etot, which is minus the log probability density of x integrated
%  over the parameter range, and parmmaxes, the vector of values of the parameters
%  at which the value of E is minimised.
% If parmstrue is passed and ntype matches parmstrue.ntype, then 
%  all the 2-d marginals are plotted with the truth overlaying them.
% nfig gives the figure number of the first to be plotted; if empty then plotting is omitted.
% If xmeangridlin is passed then it should be a vector of uniformly spaced 
%  values encompassing the total relevant range of the random variable x,
%  over which the mean will be calculated at each value of the parameters.
%  xmeanvalslin will then be a vector of values of the mean of x, one for 
%  each of the many combinations of parameter values, with the posterior probability
%  (NOT probability density) of each in the corresponding vector Pxmeanvals.
% If xmeangridlog is passed, then it should be a vector of values of the random
%  variable x whose logarithms are uniformly spaced, and the values of 
%  xmeanvalslog and Pxmeanvals will be returned correspondingly;
%  note that xmeanvalslog does not contain logarithms, but arithmetic means that have 
%  been evaluated on a logarithmically spaced grid.
% If xmeangridlog is passed as above, then xmeanvalsgeom will be the geometric means
%  also evaluated on that grid.

% Change Log:
%
%     1.1          01:may:20    rfs      First version.
%     1.2          01:may:20    rfs      Appears to be working.
%     1.3          01:may:20    rfs      Now calculates joint rather than marginal parmmaxes.
%     1.5          05:may:20    rfs      Also plots fit versus true versus histogram, and takes
%                                        ntype in rather than fn.
%     1.6          05:may:20    rfs      Fixed bug to do with potentially different number of parameters
%                                        in true distribution and distribution being assessed.
%     1.7          05:may:20    rfs      bounds and absbounds put in priors.
%     1.8          06:may:20    rfs      Added computations of means, not yet debugged.
%     1.9          06:may:20    rfs      Apparently now working.
%     1.10         06:may:20    rfs      Output parameter order changed and multiple other improvements.
%     1.12         07:may:20    rfs      Comparisons with different numbers of parameters now working.
%     1.13         07:may:20    rfs      Now handles distributions that have positive support better.
%     1.31         11:may:20    rfs      Now only allows one Ngrid value to advance per round,
%                                        and it can only get one step ahead of the other Ngrid values.
%     1.32         11:may:20    rfs      Now has a different Ngridmax value for each parameter.
%     1.33         11:may:20    rfs      Typo fixed.
%     1.35         11:may:20    rfs      Termination conditions adjusted.
%     1.37         11:may:20    rfs      And again.
%     1.75         26:may:20    rfs      Removed trap to stop types 7 and 8 being tried while testing.
%     1.81         30:may:20    rfs      Removed unnecessary calculation of priors when calculating means.
%     1.82         30:may:20    rfs      Fixed silly bug in above.
%     1.85         03:jun:20    rfs      Geometric means also made available.
%     1.95         04:jul:20    rfs      Added catches for empty x.

mytitl = ' /home/rfs/ramakrishnan/software/cytokines/rfs/SCCS/s.gridassess.m 1.102 20/09/03 08:54:05 ';

persistent mytitldone
if isempty(mytitldone),           
   mytitldone = titlfunction(mytitl);
end

global titls

if nargin < 4,
   parmstrue = [];
end
if nargin < 5,
   nfig = [];
end
if nargin < 6,
   xmeangridlin = [];
end
if nargin < 7,
   xmeangridlog = [];
end

if 0,
   % Temporary trap to avoid crashing other things while this is being tested.
   if ntype == 7 || ntype == 8,
      error('gridassess has been called with ntype 7 or 8 which is trapped during testing');
   else
      warning('gridassess currently contains a trap for types 7 and 8 which needs removing');
   end
end

Ngridmax = priors.Ngridmax{ntype};
shrinklimit = priors.shrinklimit(ntype);
typename = priors.typelist{ntype};
prior = getfield(priors, typename);
bounds = prior.bounds;
absbounds = prior.absbounds;

if isempty(absbounds),
   absbounds = bounds;
end

Nparams = size(bounds, 2);
if size(bounds, 1) ~= 2,
   error('Must give lower and upper bounds for each parameter');
end
if any(bounds(1, :) >= bounds(2, :)) || any(isinf(bounds(:))) || any(isnan(bounds(:))),
   error('Cannot have infinite or NaN bounds and bounds(1, :) must be less than bounds(2, :)');
end
if any(bounds(1, :) < absbounds(1, :)) || any(bounds(2, :) > absbounds(2, :)),
   error('Given bounds are already illegal according to absbounds');
end

if ~isempty(parmstrue),
   if isstruct(parmstrue),
      parmstruevals = struct2cell(parmstrue);
      parmstruevals = parmstruevals{2};
      if isstruct(parmstruevals),
         parmstruevals = struct2cell(parmstruevals);
         parmstruevals = cat(1, parmstruevals{:});
      else
         error('parmstrue should be a structure with the distribution type and parameters, the latter in turn also a structure');
      end
   else
      error('parmstrue should be a structure with the distribution type and parameters, the latter in turn also a structure');
   end
end

% Initialise grid resolution.
Ngrid = repmat(5, [Nparams, 1]);

if any(Ngrid > Ngridmax),
   error(sprintf('Ngridmax is infeasibly low for %d parameters', Nparams));
end

niter = 1;
while all(Ngrid <= Ngridmax),

   fprintf('Starting grid iteration %d, bounds and grid sizes are:\n', niter);
   [bounds; Ngrid.']

   parmgrid = cell(Nparams, 1);
   parmspacing = NaN(Nparams, 1);
   for nparam = 1 : Nparams,
      parmspacing(nparam) = (bounds(2, nparam) - bounds(1, nparam)) / Ngrid(nparam);
      parmgrid{nparam} = bounds(1, nparam) + [0 : Ngrid(nparam)] * parmspacing(nparam);
   end

   % Slight complication here as we have to cope with the possibility that x is empty.
   if ~isempty(x),
      parmgrid{Nparams + 1} = x;
      parmndgrid = cell(Nparams + 1, 1);
      [parmndgrid{:}] = ndgrid(parmgrid{:});
   
      E = feval(priors.Efns{ntype}, parmndgrid, priors);
   else
      parmgrid{Nparams + 1} = 1; % Dummy value.
      parmndgrid = cell(Nparams + 1, 1);
      [parmndgrid{:}] = ndgrid(parmgrid{:});
      
      E = feval(priors.Efns{ntype}, parmndgrid, priors);

      % Remove the dummy values again.
      parmgrid{Nparams + 1} = x;
      sz = size(E);
      sz(Nparams + 1) = 1;
      E = reshape(E(prod(sz) + 1 : 2 * prod(sz)), sz);
   end

   % Combine the densities at all the x values
   %  and the prior (which is at the last coord of the last dimension).
   E = sum(E, Nparams + 1);
   
   % Now, for each parameter, assess whether the bounds are appropriate.
   oldbounds = bounds;
   newbounds = bounds;
   oldNgrid = Ngrid;
   newNgrid = Ngrid;
   Etot = NaN(Nparams, 1);
   for nparam = 1 : Nparams,

      % Construct the marginal density.
      thisE = E;
      for ntosum = 1 : Nparams,
         if ntosum ~= nparam,
            thisE = Esum(thisE, ntosum) - log(parmspacing(ntosum));
         end
      end
      Etot(nparam) = Esum(thisE, nparam) - log(parmspacing(nparam));

      % Find the maximum of the marginal density.
      thisE = thisE(:);
      [minE, argmin(nparam)] = min(thisE);

      % Find the first and last points which exceed shrinklimit times the maximum density.
      thresh = minE - log(shrinklimit);
      indfirst = find(thisE <= thresh, 1, 'first') - 2;
      indlast = find(thisE <= thresh, 1, 'last') + 2;

      % Provisionally set the new bounds to the found range plus 2 in each direction.
      newbounds(:, nparam) = [bounds(1, nparam) + (indfirst - 1) * parmspacing(nparam); ...
                              bounds(1, nparam) + (indlast - 1) * parmspacing(nparam)];

      % ...capping at the absolute limits if needed.
      newbounds(1, nparam) = max(newbounds(1, nparam), absbounds(1, nparam));
      newbounds(2, nparam) = min(newbounds(2, nparam), absbounds(2, nparam));

      % If the bounds haven't changed much, then increase Ngrid(nparam), but never
      %  allow more than one such to increase at a time, and never 
      %  allow it to be more than one step ahead of the others.

      if (abs(newbounds(1, nparam) - oldbounds(1, nparam)) < 0.5 * parmspacing(nparam) ...
          && abs(newbounds(2, nparam) - oldbounds(2, nparam)) < 0.5 * parmspacing(nparam) ...
          && all(Ngrid == newNgrid) ...
          && all(Ngrid >= Ngrid(nparam) | Ngrid == Ngridmax) ...
          && (Ngrid(nparam) < Ngridmax(nparam) || all(Ngrid == Ngridmax))),
          newNgrid(nparam) = 2 * Ngrid(nparam);
      end

   end % for nparam = 1 : Nparams

   fprintf('Completed grid iteration %d.\n', niter);
   fprintf('Etot values are currently (should all be roughly the same):\n');
   Etot.'

   bounds = newbounds;
   Ngrid = newNgrid;
   niter = niter + 1;
end % while all(Ngrid <= Ngridmax)

% We exit here when the new grid is too memory-consumptive, so step back from it.
bounds = oldbounds;
Ngrid = oldNgrid;

% Calculate the joint parmmaxes.
[Epeak, argpeak] = min(E(:));
indmaxes = cell(Nparams, 1);
[indmaxes{:}] = ind2sub(size(E), argpeak);
indmaxes = cat(1, indmaxes{:});
parmmaxes = NaN(Nparams, 1);
for nparam = 1 : Nparams,
   parmmaxes(nparam) = bounds(1, nparam) + (indmaxes(nparam) - 1) * parmspacing(nparam);
end

Etot = mean(Etot);

if ~isempty(nfig),
   nfig = nfig - 1;
   if 1,
      % We plot the 2-d marginals of the posterior.
   
      for nparam1 = 1 : Nparams,
         for nparam2 = nparam1 + 1 : Nparams,
         
            % Make the 2-d marginal.
            thisE = E;
            for ntosum = 1 : Nparams,
               if ntosum ~= nparam1 && ntosum ~= nparam2,
                  thisE = Esum(thisE, ntosum) - log(parmspacing(ntosum));
               end
            end
            thisE = squeeze(thisE);
   
            minthisE = min(thisE(:));
            thisE = thisE - minthisE;
            thisP = exp(-thisE);
            thisP = thisP / (sum(thisP(:)) * prod(parmspacing([nparam1; nparam2])));
   
            nfig = nfig + 1;
            figure(nfig);
            clf;
            [theseparms1, theseparms2] = ndgrid(parmgrid{nparam1}, parmgrid{nparam2});
            surf(theseparms1, theseparms2, thisP);
            shading('interp');
            view(0, 90);
            colorbar;
            ax = bounds(:, [nparam1, nparam2]);
            axis(ax(:).');
            hold on;
            plot3(parmmaxes(nparam1), parmmaxes(nparam2), 2 * abs(max(thisP(:))), 'wo');
            plot3(parmmaxes(nparam1), parmmaxes(nparam2), 2 * abs(max(thisP(:))), 'kx');
            if ~isempty(parmstrue) && ntype == parmstrue.ntype,
               plot3(parmstruevals(nparam1), parmstruevals(nparam2), 2 * abs(max(thisP(:))), 'wx');
               plot3(parmstruevals(nparam1), parmstruevals(nparam2), 2 * abs(max(thisP(:))), 'ko');
            end
            hold off;
            xlabel(sprintf('Parameter %d', nparam1));
            ylabel(sprintf('Parameter %d', nparam2));
            title('Posterior probability density');
         end
      end
   end
   
   if ~isempty(x),
      Nx = round(length(x) * 0.5);
      minx = min(x);
      maxx = max(x);
   else
      Nx = 1;
      minx = 1e-2;
      maxx = 400;
   end
   deltax = (maxx - minx) / (Nx / 4);
   xx = [minx - 4 * deltax : deltax : maxx + 4 * deltax];

   if priors.posonly(ntype) || (~isempty(parmstrue) && priors.posonly(parmstrue.ntype)),
      xx = xx(xx > 0);
   end

   Nxcurve = 10 * Nx;
   deltaxcurve = deltax / 10;
   xxcurve = min(xx) + [0 : Nxcurve].' * deltaxcurve;

   paramsmax = cell(Nparams + 1, 1);
   parmsmaxcell = mat2cell(parmmaxes, ones(Nparams, 1));
   [paramsmax{:}] = ndgrid(parmsmaxcell{:}, xxcurve);
   Emax = feval(priors.Efns{ntype}, paramsmax, [], 1);
   
   if ~isempty(parmstrue),
      Nparamstrue = length(parmstruevals);
      paramstrue = cell(Nparamstrue + 1, 1);
      parmstruecell = mat2cell(parmstruevals, ones(Nparamstrue, 1));
      [paramstrue{:}] = ndgrid(parmstruecell{:}, xxcurve);
      Etrue = feval(priors.Efns{parmstrue.ntype}, paramstrue, [], 1);
   end
   
   nfig = nfig + 1;
   figure(nfig);
   clf;
   hist(x, xx);
   hold on;
   plot(xxcurve, length(x) * deltax * exp(-Emax(:)), 'b');
   if ~isempty(parmstrue),
      plot(xxcurve, length(x) * deltax * exp(-Etrue(:)), 'g');
   end
   hold off;
   xlabel('x');
   if ~isempty(parmstrue)
      title(sprintf('Best %s fit in blue, true %s in green', ...
                    priors.typelist{ntype}, priors.typelist{parmstrue.ntype}));
   else
      title(sprintf('Best %s fit in blue', priors.typelist{ntype}));
   end

   if all(x > 0),

      lx = log(x);
   
      if length(x) > 0,
         Nx = round(length(lx) * 0.5);
         maxlx = max(lx);
         minlx = min(lx);
      else
         Nx = 1;
         maxlx = log(400);
         minlx = log(1e-2);
      end
      deltalx = (maxlx - minlx) / (Nx / 4);
      lxx = [minlx - 4 * deltalx : deltalx : maxlx + 4 * deltalx].';
      xxl = exp(lxx);

      Nxcurve = Nx * 10;
      deltalxcurve = deltalx / 10;
      lxxcurve = min(lxx) + [0 : Nxcurve].' * deltalxcurve;
      xxlcurve = exp(lxxcurve);

      paramsmax = cell(Nparams + 1, 1);
      parmsmaxcell = mat2cell(parmmaxes, ones(Nparams, 1));
      [paramsmax{:}] = ndgrid(parmsmaxcell{:}, xxlcurve);
      Emax = feval(priors.Efns{ntype}, paramsmax, [], 1) - log(paramsmax{Nparams + 1});
      
      if ~isempty(parmstrue),
         Nparamstrue = length(parmstruevals);
         paramstrue = cell(Nparamstrue + 1, 1);
         parmstruecell = mat2cell(parmstruevals, ones(Nparamstrue, 1));
         [paramstrue{:}] = ndgrid(parmstruecell{:}, xxlcurve);
         Etrue = feval(priors.Efns{parmstrue.ntype}, paramstrue, [], 1) - log(paramstrue{Nparamstrue + 1});
      end
      
      nfig = nfig + 1;
      figure(nfig);
      clf;
      hist(lx, lxx);
      hold on;
      plot(lxxcurve, length(lx) * deltalx * exp(-Emax(:)), 'b');
      if ~isempty(parmstrue),
         plot(lxxcurve, length(lx) * deltalx * exp(-Etrue(:)), 'g');
      end
      hold off;
      xlabel('log(x)');
      if ~isempty(parmstrue)
         title(sprintf('Best %s fit in blue, true %s in green', ...
                       priors.typelist{ntype}, priors.typelist{parmstrue.ntype}));
      else
         title(sprintf('Best %s fit in blue', priors.typelist{ntype}));
      end

   end % if all(x > 0)

end % if ~isempty(nfig)

% Now time to evaluate the means at each gridpoint.

%  Default returns.
xmeanvalslin = [];
xmeanvalslog = [];
xmeanvalsgeom = [];
Pxmeanvals = [];
   
if nargout > 2,
   % First work out the posterior probability at each gridpoint.
   minE = min(E(:));
   E = E - minE;
   P = exp(-E);
   P = P(:);
   sumP = sum(P);
   Pxmeanvals = P / sumP;
end
   
% First the mean of x.

if ~isempty(xmeangridlin) && nargout > 3,
   % We use parmgrid and xmeangridlin to evaluate the mean at
   %  each point of the parameter grid.

   fprintf('Calculating means on linear grid...');

   parmgrid{Nparams + 1} = xmeangridlin(:);
   parmndgrid = cell(Nparams + 1, 1);
   [parmndgrid{:}] = ndgrid(parmgrid{:});

   Exmeanlin = feval(priors.Efns{ntype}, parmndgrid, [], 1);

   % Reshape to only leave x in the second dimension,
   %  remembering that the last entry in that dimension is the prior, which needs removing.
   sz = size(parmndgrid{1});
   Exmeanlin = reshape(Exmeanlin, [prod(sz(1 : Nparams)), sz(Nparams + 1)]);

   % Normalise along the x direction.
   minExmeanlin = min(Exmeanlin, [], 2);
   Exmeanlin = Exmeanlin - minExmeanlin(:, ones(1, size(Exmeanlin, 2)));
   Pxmeanlin = exp(- Exmeanlin);
   sumPxmeanlin = sum(Pxmeanlin, 2);
   Pxmeanlin = Pxmeanlin ./ sumPxmeanlin(:, ones(1, size(Exmeanlin, 2)));
   % making Pxmeanlin an array of probabilities, not of densities.

   % Calculate the means.
   xmeangridlin = xmeangridlin(:).';
   xmeanvalslin = sum(Pxmeanlin .* xmeangridlin(ones(size(Pxmeanlin, 1), 1), :), 2);

   fprintf('done.\n');
end % if ~isempty(xmeangridlin) && nargout > 2 or not

% Second the arithmetic mean of x on a log-spaced grid.

if ~isempty(xmeangridlog) && nargout > 4,
   % We use parmgrid and xmeangridlog to evaluate the mean at
   %  each point of the parameter grid.

   fprintf('Calculating means on log grid...');

   parmgrid{Nparams + 1} = xmeangridlog(:);
   parmndgrid = cell(Nparams + 1, 1);
   [parmndgrid{:}] = ndgrid(parmgrid{:});

   Exmeanlog = feval(priors.Efns{ntype}, parmndgrid, [], 1);
   % Will need correcting for being density of log later.

   % Reshape to only leave x in the second dimension.
   sz = size(parmndgrid{1});
   Exmeanlog = reshape(Exmeanlog, [prod(sz(1 : Nparams)), sz(Nparams + 1)]);
   Exmeanlog = Exmeanlog - log(reshape(parmndgrid{Nparams + 1}, size(Exmeanlog)));

   % Normalise along the x direction.
   minExmeanlog = min(Exmeanlog, [], 2);
   Exmeanlog = Exmeanlog - minExmeanlog(:, ones(1, size(Exmeanlog, 2)));
   Pxmeanlog = exp(- Exmeanlog);
   sumPxmeanlog = sum(Pxmeanlog, 2);
   Pxmeanlog = Pxmeanlog ./ sumPxmeanlog(:, ones(1, size(Exmeanlog, 2)));
   % making Pxmeanlog an array of probabilities, not of densities.

   % Calculate the means.
   xmeangridlog = xmeangridlog(:).';
   xmeanvalslog = sum(Pxmeanlog .* xmeangridlog(ones(size(Pxmeanlog, 1), 1), :), 2);
   if nargout > 5,
      xmeanvalsgeom = exp(sum(Pxmeanlog .* log(xmeangridlog(ones(size(Pxmeanlog, 1), 1), :)), 2));
   end

   fprintf('done.\n');
end % if ~isempty(xmeangridlog) && nargout > 4 or not

nfig = nfig + 1;
return;


% Local Variables: 
% indent-line-function: indent-relative
% eval: (auto-fill-mode 0)
% End:
```
