## Supplementary material for "Elevated cerebrospinal fluid cytokine levels in tuberculous meningitis predict survival in response to dexamethasone": Software: ksample.m

```
function y = ksample(a_k, b_k, c_k, N_k)
% Usage: y = ksample(a_k, b_k, c_k, N_k)
% Samples k from the prior for the Gamma-power distribution.

% Change Log:
%
%     1.1          25:may:20    rfs      Taken from survival project k_prior_sample.m v1.56 and adapted.

mytitl = ' /home/rfs/ramakrishnan/software/cytokines/rfs/SCCS/s.ksample.m 1.102 20/09/03 08:54:05 ';

persistent mytitldone
if isempty(mytitldone),           
   mytitldone = titlfunction(mytitl);
end

global titls

logf = @(x) log_k_density(x, a_k, b_k, c_k, N_k);
gradlogf = @(x) log_k_derivative(x, a_k, b_k, c_k, N_k);

% Come up with some suitable points to initiate ARS with
% (see ars.m).

p_points = [1,2];

while log_k_derivative(p_points(1), a_k, b_k, c_k, N_k) <= 0
    p_points = [p_points(1)/2, p_points];
end
while log_k_derivative(p_points(end), a_k, b_k, c_k, N_k) >= 0
    p_points = [p_points, 2*p_points(end)];
end

k = 2*rand - 1;

y = ars(logf, gradlogf, p_points, 0, Inf);

return;

% Local Variables: 
% indent-line-function: indent-relative
% eval: (auto-fill-mode 0)
% End:
```
