## Supplementary material for "Elevated cerebrospinal fluid cytokine levels in tuberculous meningitis predict survival in response to dexamethasone": Software: lhgmex2.c

```
/* This file contains code that is intended to speed up the calculation
 * of the log of the hypergeometric function 1F1 for positive arguments
 * of which the second is scalar and the others are matrices of the same size.
 *
 * It is many many times faster than doing the same thing in Matlab, but
 * for arguments where it cannot first be calculated in the linear domain,
 * it is 20 times slower than for those where it can.
 *
 * It is intended only to be called from loghypergeom1F1v2.m .
 */

/* To compile:
 *  /opt/MATLAB/R2012b/bin/mex -f /home/rfs/matlab/gccopts.sh -largeArrayDims CFLAGS='-ansi -fno-fast-math -Wall -fexceptions -fPIC -DARCH64BIT' -output lhgmex2.mexa64 lhgmex2.c -lm
 *
 * You can ignore a warning that
 *   You are using gcc version "8.3.1".  The version currently supported with MEX is "4.4.6".
 *
 */

/* Change Log:
 *
 *    1.1          14:may:20    rfs  First version, not yet completely working.
 *    1.50         14:may:20    rfs  Version number harmonisation.
 *    1.51         14:may:20    rfs  Added version in the -log domain, seems to be working.
 *
 *  /home/rfs/ramakrishnan/software/cytokines/rfs/SCCS/s.lhgmex2.c 1.102 20/09/03 08:54:05
 */

#include <stdlib.h>
#include <stdarg.h>
#include <math.h>
#include <string.h>
#include <sys/types.h>
#include <limits.h>
#include <unistd.h>

/* Matlab-specific header files. */
#include "mex.h"
#include "matrix.h"


double NaN;
double Inf;

void
initNaNInf(void)
{
  NaN = mxGetNaN();
  Inf = mxGetInf();
}


int
isnand(double x)
{
  return(mxIsNaN(x));
}


int
isinfd(double x)
{
  return(mxIsInf(x));
}


/* Die, giving an error message.
 * The arguments are exactly like those of printf.
 */
void
Die(char *format, ...)
{
  char s[1000];
  int nused = 0;
  va_list argp;

  /* format the error mesg */

  nused = sprintf(s, "FATAL: ");
  va_start(argp, format);
  nused += vsprintf(s + nused, format, argp);
  va_end(argp);
  sprintf(s + nused, "\n");
  mexErrMsgTxt(s); /* Exit. */

  return;	
}


/* As for Die, but just gives a warning. */
void
Warn(char *format, ...)
{
  va_list  argp;
  char s[1000];
  int nused = 0;

  nused = sprintf(s, "WARNING: ");
  va_start(argp, format);
  nused += vsprintf(s + nused, format, argp);
  va_end(argp);
  sprintf(s + nused, "\n");

  mexWarnMsgTxt(s);

  return;
}


/* Return the int nearest to in. */
int
makeintfromdouble(double in)
{
  static int warned = 0;
  if (!warned && (in > 0.9 * INT_MAX || in < -0.9 * INT_MAX))
    {
      Warn("Conversion to int of a double dangerously close to LONG_MAX");
      warned = 1;
    }
  if (in >= 0)
    return (long int) (in + 0.5);
  else
    return (long int) (in - 0.5);
}


static int
scalarcheck (const mxArray *A)
{
  if (mxGetNumberOfElements (A) != 1)
    return 0;
  else
    return 1;
}


/* This function adds two numbers in the -log domain. */
double Eadd (double x1, double x2)
{
  double out = NaN;
  
  if (x1 < x2)
    out = x1 - log (1 + exp (x1 - x2));
  if (x1 >= x2)
    out = x2 - log (1 + exp (x2 - x1));
  if (x1 == x2 && isinfd (x1))
    out = x1;
  
  return(out);
}


/* Function: checkdims()
 *
 * Purpose:  Checks whether the passed array has the specified size
 *           to within the ambiguity of [A, B, C, 1, 1] is equivalent to [A, B, C].
 *
 * Args:     A - pointer to the mxArray to be checked
 *           sz - pointer to vector of ints containing the appropriate dimensions.
 *           Ndims - length of sz
 *
 * Returns:  1 if good, 0 if bad.
 */
int
checkdims(const mxArray *A,
          const size_t *sz,
          const int Ndims)
{
  int NdimsA = mxGetNumberOfDimensions(A);
  int ndim;
  int bad = 0;
  const size_t *dimsA = mxGetDimensions(A);

  for (ndim = 0; ndim < Ndims && ndim < NdimsA; ndim++)
    if (dimsA[ndim] != sz[ndim])
      bad = 1;
  for (; ndim < Ndims; ndim++)
    if (sz[ndim] != 1)
      bad = 1;
  for (; ndim < NdimsA; ndim++)
    if (dimsA[ndim] != 1)
      bad = 1;

  return(!bad);
}

/* This is the core function of this file.
 *
 * This function calculates the logarithm of the hypergeometric function
 * log(1F1(alpha, gamma, z)) while avoiding overflow of 1F1 itself.
 *
 * For this function, alpha, gamma, and z are all positive real scalars.
 *
 * The general case is that starting from zero, the magnitudes of the
 * terms of the defining series may:
 *  - decrease, then increase, then decrease again; or
 *  - increase, then decrease again; or
 *  - just decrease from the start.
 * So there may be local maxima either at zero and one other place,
 * just at zero, or just at one other place.
 *
 * Call by:
 *   out = lhg1F1(alpha, gamma, z, n1, n3, Eterm1, Eterm3);
 * where out is a double return, and alpha, gamma, and z are doubles,
 * and n1 and n3 are the locations of the two possible biggest terms,
 * and Eterm and Eterm2 are minus the logs of their values
 * If n1 is negative then there is no local maximum at zero,
 * in which case Eterm1 will be Inf; but it must be passed in negative
 * to ensure it doesn't prevent term zero being used.
 *
 * The basic plan is to start with the biggest terms, then work away from
 * them until they are too small to make any further difference to the answer.
 */
double
lhg1F1(double alpha, double gamma, double z, int n1, int n3, double Eterm1, double Eterm3)
{
  double outval = 0;
  int n2;
  double ratio;
  double lastoutval = 0;
  int outputchanged = 1;
  double term1 = exp(-Eterm1);
  double Eterm2 = Eterm3;
  double term2 = 0;
  double term3 = exp(-Eterm3);
  const unsigned long int basicN = 10000;
  const unsigned long int stageN = (n3 > basicN ? n3 : basicN);
  static int warned = 0;

  /* A absolute upper limit on terms to be used. */  
  const unsigned long int N = (2 * fabs(z) > stageN ? 2 * fabs(z) : stageN);

  if ((2 * fabs(z) > ULONG_MAX) && !warned)
    {
      Warn("Parameters too big for safety in lhgmex2");
      warned = 1;
    }

  /* It has been checked by the Matlab calling function that alpha, gamma, and z are real and positive,
   * except that z may be zero.
   */

  /* We first try calculating in the linear domain as it is quicker.
   * If that fails, we will resort to the -log domain.
   */

  /*  |  set this to 1 to force the -log domain method to be used. */
  if (0 || ((alpha >= gamma && z > 709.0) /* 709.0 is roughly the largest real whose exponential is not infinite */
            || Eterm1 < -709.0 || Eterm3 < -709.0))
    {
      outval = Inf; /* Not worth trying the linear domain in this case. */
    }
  else
    {
      n2 = (n3 > 0 ? n3 - 1 : -1);
      term2 = (n2 >= 0 ? term3 * ((gamma + n2) * (1 + n2)) / ((alpha + n2) * z) : 0);

      while (n3 < N && outputchanged)
        {
          outputchanged = 0;
          if (term3 > 0)
            {
              lastoutval = outval;
              outval += term3;
              
              ratio = (alpha + n3) * z / ((gamma + n3) * (1 + n3));
          
              term3 *= ratio;

              if (outval == lastoutval)
                term3 = 0;
              else
                outputchanged = 1;
          
              n3++;
            }

          if (term2 > 0 && n2 >= n1 && n2 >= 0)
            {
              lastoutval = outval;
              outval += term2;

              n2--;

              ratio = ((gamma + n2) * (1 + n2) / ((alpha + n2) * z));

              term2 *= ratio;

              if (outval == lastoutval)
                term2 = 0;
              else
                outputchanged = 1;
            }
          else
            term2 = 0;

          if (term1 > 0 && n1 >= 0 && n1 <= n2)
            {
              lastoutval = outval;
              outval += term1;

              ratio = ((alpha + n1) * z) / ((gamma + n1) * (1 + n1));

              term1 *= ratio;

              if (outval == lastoutval)
                term1 = 0;
              else
                outputchanged = 1;

              n1++;
            }
          else
            term1 = 0;

#if 0
          mexPrintf("n1=%d, n2=%d, n3=%d, term1=%g, term2=%g, term3=%g, outval=%g\n",
                    n1, n2, n3, term1, term2, term3, outval);
          
          mexCallMATLAB(0, NULL, 0, NULL, "pause");
#endif
        }
      outval = log(outval);
    }

  if (!isinfd(outval) && !isnand(outval))
    return(outval);

  /* Otherwise we need to do the calculation over again in the -log domain. */

  /* Reinitialise. */
  outval = Inf;
  lastoutval = Inf;
  outputchanged = 1;

  n2 = (n3 > 0 ? n3 - 1 : -1);
  Eterm2 = (n2 >= 0 ? Eterm3 - log(((gamma + n2) * (1 + n2)) / ((alpha + n2) * z)) : Inf);

  while (n3 < N && outputchanged)
    {
      outputchanged = 0;
      if (Eterm3 != Inf)
        {
          lastoutval = outval;
          outval = Eadd(outval, Eterm3);
              
          ratio = (alpha + n3) * z / ((gamma + n3) * (1 + n3));
          
          Eterm3 -= log(ratio);

          if (outval == lastoutval)
            Eterm3 = Inf;
          else
            outputchanged = 1;
          
          n3++;
        }

      if (Eterm2 != Inf && n2 >= n1 && n2 >= 0)
        {
          lastoutval = outval;
          outval = Eadd(outval, Eterm2);

          n2--;

          ratio = ((gamma + n2) * (1 + n2) / ((alpha + n2) * z));

          Eterm2 -= log(ratio);

          if (outval == lastoutval)
            Eterm2 = Inf;
          else
            outputchanged = 1;
        }
      else
        Eterm2 = Inf;

      if (Eterm1 != Inf && n1 >= 0 && n1 <= n2)
        {
          lastoutval = outval;
          outval = Eadd(outval, Eterm1);

          ratio = ((alpha + n1) * z) / ((gamma + n1) * (1 + n1));

          Eterm1 -= log(ratio);

          if (outval == lastoutval)
            Eterm1 = Inf;
          else
            outputchanged = 1;

          n1++;
        }
      else
        Eterm1 = Inf;

#if 0
      mexPrintf("n1=%d, n2=%d, n3=%d, term1=%g, term2=%g, term3=%g, outval=%g\n",
                n1, n2, n3, Eterm1, Eterm2, Eterm3, outval);
      
      mexCallMATLAB(0, NULL, 0, NULL, "pause");
#endif
    }

  return(-outval);
}


/* This is the mex wrapper function which collects the arguments
 * and checks the sizes of the arrays.
 */
void
mexFunction(int nlhs,
            mxArray *plhs[],
            int nrhs,
            const mxArray *prhs[])
{
  int Ndims;
  int ndim;
  unsigned long int Nelements;
  unsigned long int nelement;
  static int infchecked = 0;
  double *outptr;
  double *alphaptr;
  double *gammaptr;
  double *zptr;
  double *n1ptr;
  double *n3ptr;
  double *Eterm1ptr;
  double *Eterm3ptr;

  static int initialised = 0;
  if (!initialised)
    {
      initNaNInf();
      initialised = 1;
    }

  /* check NaN and Inf */
  if (!infchecked)
    {
      if (NaN == NaN) 
        Warn ("NaN is equal to itself on this platform (it should not be)");
      if (Inf != Inf) 
        Warn ("Inf is not equal to itself on this platform (it should be)");
      if (!isnand (NaN))
        Warn ("isnand () not working on this platform");
      if (!isinfd (Inf))
        Warn ("isinfd () not working on this platform");

      infchecked = 1;
    }

  if (nlhs != 1 || nrhs != 7)
    Die("Usage: out = lhgmex2(alpha, gamma, z, n1, n3, Eterm1, Eterm3)");

  /* Block that checks the six non-scalar inputs have the same size,
   * then allocates the output array of that size.
   */
  {
    const size_t *dims = mxGetDimensions(prhs[0]);
    Ndims = mxGetNumberOfDimensions(prhs[0]);

    if (!checkdims(prhs[2], dims, Ndims)
        || !scalarcheck(prhs[1])
        || !checkdims(prhs[3], dims, Ndims)
        || !checkdims(prhs[4], dims, Ndims)
        || !checkdims(prhs[5], dims, Ndims)
        || !checkdims(prhs[6], dims, Ndims))
      Die("Non-scalar inputs do not have the same size or gamma is not a scalar");

    /* Create the output array. */
    plhs[0] = mxCreateNumericArray(Ndims, dims, mxDOUBLE_CLASS, mxREAL);

    Nelements = 1;
    for (ndim = 0; ndim < Ndims; ndim++)
      Nelements *= dims[ndim];
  }

  /* Initialise the pointers. */
  outptr = mxGetPr(plhs[0]);
  alphaptr = mxGetPr(prhs[0]);
  gammaptr = mxGetPr(prhs[1]);
  zptr = mxGetPr(prhs[2]);
  n1ptr = mxGetPr(prhs[3]);
  n3ptr = mxGetPr(prhs[4]);
  Eterm1ptr = mxGetPr(prhs[5]);
  Eterm3ptr = mxGetPr(prhs[6]);

  /* Loop over elements. */
  for (nelement = 0; nelement < Nelements; nelement++)
    *outptr++ = lhg1F1(*alphaptr++, *gammaptr, *zptr++,
                       makeintfromdouble(*n1ptr++), makeintfromdouble(*n3ptr++),
                       *Eterm1ptr++, *Eterm3ptr++);

  return;
}
```
