## Supplementary material for "Elevated cerebrospinal fluid cytokine levels in tuberculous meningitis predict survival in response to dexamethasone": Software: lhgmex.c

```
/* This file contains code that is intended to speed up the calculation
 * of the log of the hypergeometric function 1F1 for positive arguments
 * of which the second is scalar and the others are matrices of the same size.
 *
 * It is many many times faster than doing the same thing in Matlab, but
 * for arguments where it cannot first be calculated in the linear domain,
 * it is 20 times slower than for those where it can.
 */

/* To compile:
 *  /opt/MATLAB/R2012b/bin/mex -f /home/rfs/matlab/gccopts.sh -largeArrayDims CFLAGS='-ansi -fno-fast-math -Wall -fexceptions -fPIC -DARCH64BIT' -output loghypergeom1F1.mexa64 lhgmex.c -lm
 *
 * You can ignore a warning that
 *   You are using gcc version "8.3.1".  The version currently supported with MEX is "4.4.6".
 *
 */

/* Change Log:
 *
 *    1.1          13:may:20    rfs  First version.
 *    1.44         13:may:20    rfs  Compiles OK now, version number harmonisation.
 *
 *  /home/rfs/ramakrishnan/software/cytokines/rfs/SCCS/s.lhgmex.c 1.102 20/09/03 08:54:05
 */

#include <stdio.h>
#include <stdlib.h>
#include <stdarg.h>
#include <math.h>
#include <string.h>
#include <sys/types.h>
#include <time.h>   

/* Matlab-specific header files. */
#include "mex.h"
#include "matrix.h"


double NaN;
double Inf;

void
initNaNInf(void)
{
  NaN = mxGetNaN();
  Inf = mxGetInf();
}


int
isnand(double x)
{
  return(mxIsNaN(x));
}


int
isinfd(double x)
{
  return(mxIsInf(x));
}


/* Die, giving an error message.
 * The arguments are exactly like those of printf.
 */
void
Die(char *format, ...)
{
  char s[1000];
  int nused = 0;
  va_list argp;

  fflush(stdout);
  fflush(stderr);

  /* format the error mesg */

  nused = sprintf(s, "FATAL: ");
  va_start(argp, format);
  nused += vsprintf(s + nused, format, argp);
  va_end(argp);
  sprintf(s + nused, "\n");
  mexErrMsgTxt(s); /* Exit. */

  return;	
}


/* As for Die, but just gives a warning. */
void
Warn(char *format, ...)
{
  va_list  argp;
  char s[1000];
  int nused = 0;

  fflush(stdout);
  fflush(stderr);

  nused = sprintf(s, "WARNING: ");
  va_start(argp, format);
  nused += vsprintf(s + nused, format, argp);
  va_end(argp);
  sprintf(s + nused, "\n");

  mexWarnMsgTxt(s);

  return;
}


static int
scalarcheck (const mxArray *A)
{
  if (mxGetNumberOfElements (A) != 1)
    return 0;
  else
    return 1;
}


/* This function adds two numbers in the -log domain. */
double Eadd (double x1, double x2)
{
  double out = NaN;
  
  if (x1 < x2)
    out = x1 - log (1 + exp (x1 - x2));
  if (x1 >= x2)
    out = x2 - log (1 + exp (x2 - x1));
  if (x1 == x2 && isinfd (x1))
    out = x1;
  
  return(out);
}


/* Function: checkdims()
 *
 * Purpose:  Checks whether the passed array has the specified size
 *           to within the ambiguity of [A, B, C, 1, 1] is equivalent to [A, B, C].
 *
 * Args:     A - pointer to the mxArray to be checked
 *           sz - pointer to vector of ints containing the appropriate dimensions.
 *           Ndims - length of sz
 *
 * Returns:  1 if good, 0 if bad.
 */
int
checkdims(const mxArray *A,
          const size_t *sz,
          const int Ndims)
{
  int NdimsA = mxGetNumberOfDimensions(A);
  int ndim;
  int bad = 0;
  const size_t *dimsA = mxGetDimensions(A);

  for (ndim = 0; ndim < Ndims && ndim < NdimsA; ndim++)
    if (dimsA[ndim] != sz[ndim])
      bad = 1;
  for (; ndim < Ndims; ndim++)
    if (sz[ndim] != 1)
      bad = 1;
  for (; ndim < NdimsA; ndim++)
    if (dimsA[ndim] != 1)
      bad = 1;

  return(!bad);
}


/* This is the core function of this file.
 *
 * This function calculates the logarithm of the hypergeometric function
 * log(1F1(alpha, gamma, z)) while avoiding overflow of 1F1 itself.
 *
 * For this function, alpha, gamma, and z are all positive real scalars.
 *
 * Call by:
 *   out = lhg1F1(alpha, gamma, z);
 * where out is a double return, and alpha, gamma, and z are doubles.
 */
double
lhg1F1(double alpha, double gamma, double z)
{
  double outval = 0;
  double term = 1;
  double n = 0;
  double ratio;
  int update = 1;
  double lastoutval = 0;
  int outputchanged = 1;
  const unsigned long int basicN = 10000;
  const unsigned long int N = (2 * fabs(z) > basicN ? 2 * fabs(z) : basicN);

  if (z <= 0 || alpha <= 0 || gamma <= 0)
    {
      Die("Arguments to loghypergeom1F1 must at present be positive");
    }

  /* We first try calculating in the linear domain as it is quicker.
   * If that fails, we will resort to the -log domain.
   */

  if (alpha >= gamma && z > 709.0) /* 709.0 is roughly the largest real whose exponential is not infinite */
    {
      outval = Inf; /* Not worth trying in this case. */
    }
  else
    {

      while (n < N && (n < 3 || update || outputchanged))
        {
          lastoutval = outval;
          outval += term;
          
          ratio = (alpha + n) * z / ((gamma + n) * (1 + n));
          
          term *= ratio;
          
          if (update)
            {
              update = fabs(ratio) > 1;
            }
          
          outputchanged = (outval != lastoutval);
          
          n++;
        }
      outval = log(outval);
    }

  if (!isinfd(outval) && !isnand(outval))
    return(outval);

  /* Otherwise we need to do the calculation over again in the -log domain. */

  /* Reinitialise. */
  outval = Inf;
  term = 0;
  n = 0;
  update = 1;
  lastoutval = Inf;
  outputchanged = 1;

  while (n < N && (n < 3 || update || outputchanged))
    {
      lastoutval = outval;
      outval = Eadd(outval, term);
      ratio = log((alpha + n) * z / ((gamma + n) * (1 + n)));
      term = term - ratio;

      update = update & (ratio > 0);

      outputchanged = (outval != lastoutval);

      n++;
    }
              
  return(-outval);
}


/* This is the mex wrapper function which collects the arguments
 * and checks the sizes of the arrays.
 */
void
mexFunction(int nlhs,
            mxArray *plhs[],
            int nrhs,
            const mxArray *prhs[])
{
  int Ndims;
  int ndim;
  unsigned long int Nelements;
  unsigned long int nelement;
  static int infchecked = 0;
  double *outptr;
  double *alphaptr;
  double *gammaptr;
  double *zptr;

  static int initialised = 0;
  if (!initialised)
    {
      initNaNInf();
      initialised = 1;
    }

  /* check NaN and Inf */
  if (!infchecked)
    {
      if (NaN == NaN) 
        Warn ("NaN is equal to itself on this platform (it should not be)");
      if (Inf != Inf) 
        Warn ("Inf is not equal to itself on this platform (it should be)");
      if (!isnand (NaN))
        Warn ("isnand () not working on this platform");
      if (!isinfd (Inf))
        Warn ("isinfd () not working on this platform");

      infchecked = 1;
    }

  if (nlhs != 1 || nrhs != 3)
    Die("Usage: out = loghypergeom1F1(alpha, gamma, z)");

  /* Block that checks the two non-scalar inputs have the same size,
   * then allocates the output array of that size.
   */
  {
    const size_t *dims = mxGetDimensions(prhs[0]);
    Ndims = mxGetNumberOfDimensions(prhs[0]);

    if (!checkdims(prhs[2], dims, Ndims) || !scalarcheck(prhs[1]))
      Die("alpha and z do not have the same size or gamma is not a scalar");

    /* Create the output array. */
    plhs[0] = mxCreateNumericArray(Ndims, dims, mxDOUBLE_CLASS, mxREAL);

    Nelements = 1;
    for (ndim = 0; ndim < Ndims; ndim++)
      Nelements *= dims[ndim];
  }

  /* Initialise the pointers. */
  outptr = mxGetPr(plhs[0]);
  alphaptr = mxGetPr(prhs[0]);
  gammaptr = mxGetPr(prhs[1]);
  zptr = mxGetPr(prhs[2]);

  /* Loop over elements. */
  for (nelement = 0; nelement < Nelements; nelement++)
    *outptr++ = lhg1F1(*alphaptr++, *gammaptr, *zptr++);

  return;
}
```
