## Supplementary material for "Elevated cerebrospinal fluid cytokine levels in tuberculous meningitis predict survival in response to dexamethasone": Software: log_k_density.m

```
function y = log_k_density(k, a, b, c, N)
% Usage: y = log_k_density(k, a, b, c, N)
% Returns the log density of k at the parameter values a,b,c,
%  where N is the length of b and c, unnormalised.
% Prior for k of the form:
%  P(scalar k) proportional to 
%   abs(k) .^ a .* prod(b .^ (c .* k) .* exp(- c .* b .^ k))
%  for positive scalar integer a, and vectors b and c 
%  of positive reals of the same length as each other.
% Note that zero entries in b (and the corresponding entries in c)
%  are ignored.

% Change Log:
%
%     1.1          25:may:20    rfs      Taken from survival project log_k_density.m v1.56 and adapted.
%     1.69         25:may:20    rfs      Various buggy transpositions fixed.

mytitl = ' /home/rfs/ramakrishnan/software/cytokines/rfs/SCCS/s.log_k_density.m 1.102 20/09/03 08:54:05 ';

persistent mytitldone
if isempty(mytitldone),           
   mytitldone = titlfunction(mytitl);
end

global titls

b = b(:);
c = c(:);

sz = size(k);
k = k(:).';
M = length(k);

krep = repmat(k, [N, 1]);
b = repmat(b, [1, M]);
c = repmat(c, [1, M]);

d = b;
d(b == 0) = 1;
dd = d .^ krep;
dd(b == 0) = 0;

y = log(abs(k)) .* a + sum((log(d) .* (c .* krep) - (c .* dd)), 1);

y = reshape(y, sz);

return;

% Local Variables: 
% indent-line-function: indent-relative
% eval: (auto-fill-mode 0)
% End:
```
