## Supplementary material for "Elevated cerebrospinal fluid cytokine levels in tuberculous meningitis predict survival in response to dexamethasone": Software: log_k_derivative.m

```
function y = log_k_derivative(k, a, b, c, N)
% Usage: y = log_k_derivative(k, a, b, c, N)
% Returns the gradient of the log density returned by
%  log_k_density (unnormalised).
% Note that zero entries in b (and the corresponding entries in c)
%  are ignored.

% Change Log:
%
%     1.1          25:may:20    rfs      Taken from survival project log_k_derivative.m v1.56 and adapted.
%     1.69         25:may:20    rfs      Various buggy transpositions fixed.

mytitl = ' /home/rfs/ramakrishnan/software/cytokines/rfs/SCCS/s.log_k_derivative.m 1.102 20/09/03 08:54:05 ';

persistent mytitldone
if isempty(mytitldone),           
   mytitldone = titlfunction(mytitl);
end

global titls

b = b(:);
c = c(:);

sz = size(k);
k = k(:).';

M = length(k);

kN = repmat(k, [N, 1]);
b = repmat(b, [1, M]);
c = repmat(c, [1, M]);

dd = log(b) .* (b .^ kN);
dd(b==0) = 0;
ee = c .* log(b);
ee(b==0) = 0;

y = (a ./ k) - sum(c .* dd, 1) + sum(ee, 1);

y = reshape(y, sz);

return;

% Local Variables: 
% indent-line-function: indent-relative
% eval: (auto-fill-mode 0)
% End:
```
