## Supplementary material for "Elevated cerebrospinal fluid cytokine levels in tuberculous meningitis predict survival in response to dexamethasone": Software: log_m_density.m

```
function y = log_m_density(m,a,b)
% Usage: y = log_m_density(m,a,b)
% Returns the logarithm of the prior probability density of m,
%  given by
%  P(m) proportional to m ^ (b * m) * exp(-(a + b) * m) / Gamma(m) ^ b.

% Change Log:
%
%     1.1          04:jun:20    jc2062   Taken from survival code log_m_density.m v1.56 .
%     1.86         04:jun:20    rfs34    Version number harmonisation.

mytitl = ' /home/rfs/ramakrishnan/software/cytokines/rfs/SCCS/s.log_m_density.m 1.102 20/09/03 08:54:05 ';

persistent mytitldone
if isempty(mytitldone),           
   mytitldone = titlfunction(mytitl);
end

global titls

y = (m.*b.*log(m)) - (b .* gammaln(m)) - (m.*(a+b));
end

% Local Variables: 
% indent-line-function: indent-relative
% eval: (auto-fill-mode 0)
% End:
```
