## Supplementary material for "Elevated cerebrospinal fluid cytokine levels in tuberculous meningitis predict survival in response to dexamethasone": Software: log_m_derivative.m

```
function y = log_m_derivative(m, a, b)
% Usage: y = log_m_derivative(m, a, b)
% Returns the gradient of the log density returned by log_m_density.

% Change Log:
%
%     1.1          04:jun:20    rfs34    Taken from survival code v1.86 .
%     1.86         04:jun:20    rfs      Version number harmonisation.

mytitl = ' /home/rfs/ramakrishnan/software/cytokines/rfs/SCCS/s.log_m_derivative.m 1.102 20/09/03 08:54:05 ';

persistent mytitldone
if isempty(mytitldone),           
   mytitldone = titlfunction(mytitl);
end

global titls

y = (b*log(m)) + b - (b*psi(m)) - (a+b);
end

% Local Variables: 
% indent-line-function: indent-relative
% eval: (auto-fill-mode 0)
% End:
```
