## Supplementary material for "Elevated cerebrospinal fluid cytokine levels in tuberculous meningitis predict survival in response to dexamethasone": Software: loghypergeom1F1v2.m

```
function out = loghypergeom1F1v2(alpha, gamma, z);
% Usage: out = loghypergeom1F1v2(alpha, gamma, z);
% Calculates the log of Kummer's confluent hypergeometric function 1F1.
% Currently only implemented for all arguments positive
%  and gamma scalar and alpha and z the same size.
% (Now invokes a mex function after doing some preliminary calculations.)

% Change Log:
%
%     1.1          14:may:20    rfs      First version.
%     1.2          14:may:20    rfs      Fixed bug calculating Eterm3.
%     1.52         14:may:20    rfs      Cleared some variables after use to conserve memory.
%     1.58         15:may:20    rfs      Removed the objection to z being zero which should
%                                        not have been there in the first place.

mytitl = ' /home/rfs/ramakrishnan/software/cytokines/rfs/SCCS/s.loghypergeom1F1v2.m 1.102 20/09/03 08:54:05 ';

persistent mytitldone
if isempty(mytitldone),           
   mytitldone = titlfunction(mytitl);
end

global titls

debugging = 0;

if ~all(size(alpha) == size(z)) || ~isscalar(gamma),
   error('loghypergeom1F1v2 requires alpha and z to be the same size and gamma to be a scalar');
end

if any(alpha(:) <= 0) || any(gamma(:) <= 0) || any(z(:) < 0) || any(~isreal(z(:))),
   error('loghypergeom1F1 is currently not implemented for non-positive arguments except for zero z being allowed');
end

% Find the potential positions of the biggest terms in the defining series.

w = (z - gamma - 1) / 2;
discrim = w .^ 2 - gamma + alpha .* z;

sdiscrim = NaN(size(discrim));
high = zeros(size(alpha));
low = zeros(size(alpha));

ind = discrim >= 0;
sdiscrim(ind) = sqrt(discrim(ind));
high(ind) = round(w(ind) + sdiscrim(ind));
low(ind) = round(w(ind) - sdiscrim(ind));

clear sdiscrim discrim w 

% The potential biggest terms are now roughly at positions high and zero in the series,
% with low indicating a potential local minimum.

n1 = -ones(size(alpha));
n3 = zeros(size(alpha));
Eterm1 = Inf(size(alpha));
Eterm3 = zeros(size(alpha));

% Where discrim is <= 0 we stick with the default values.

highthresh = 10;

% Where high is <= highthresh we also stick with the default values,
%  reckoning that using the first highthresh terms (which might have been needed anyhow)
%  is not too wasteful.

% Where high > highthresh:
%  If low < 0, then we leave term1 dead and set up term3 at high.
%  If low >= 0 and low < high - 1, then we also set up term1.

indforterm3 = ind & (high > highthresh);
n3(indforterm3) = high(indforterm3);
Eterm3(indforterm3) = - gammaln(alpha(indforterm3) + n3(indforterm3)) ...
                      + gammaln(alpha(indforterm3)) ...
                      + gammaln(gamma + n3(indforterm3)) ...
                      - gammaln(gamma) ...
                      - n3(indforterm3) .* log(z(indforterm3)) ...
                      + gammaln(n3(indforterm3) + 1);

indforterm1 = indforterm3 & (low >= 0) & (low < high - 1);
n1(indforterm1) = 0;
Eterm1(indforterm1) = 0;

if 0, % if debugging
   % Indicate which cases have been covered in this call.
   fprintf('Cases %d%d%d%d have been covered in this call.\n', ...
           sum(ind(:) == 0) > 0, ...
           sum(ind(:) == 1 & indforterm3(:) == 0) > 0, ...
           sum(indforterm3(:) == 1 & indforterm1(:) == 0) > 0, ...
           sum(indforterm1(:) == 1) > 0);
end

% Having identified the positions and values of the biggest terms in the series to be summed,
%  we finally call the mex function.
out = reshape(lhgmex2(alpha, gamma, z, n1, n3, Eterm1, Eterm3), size(alpha));

return;

% Local Variables: 
% indent-line-function: indent-relative
% eval: (auto-fill-mode 0)
% End:
```
