## Supplementary material for "Elevated cerebrospinal fluid cytokine levels in tuberculous meningitis predict survival in response to dexamethasone": Software: m_sample.m

```
function y = m_sample(m_a, m_b);
% Usage: y = m_sample(m_a, m_b);
% Original Usage: y = m_sample(J, r, k, x, m_a, m_b)
% This function resamples m from the prior, given the other parameters,
%  of which m_a and m_b are the parameters of the prior
%  (for the form of which see log_m_density.m).
% Original usage sampled from the posterior.
% Note that only x that are not Inf are considered.
% Method is by ARS (see ars.m).

% Change Log:
%
%     1.1          04:jun:20    rfs      Taken from survival code m_sample.m v1.86 and adapted.
%     1.86         04:jun:20    rfs      Version number harmonisation.

mytitl = ' /home/rfs/ramakrishnan/software/cytokines/rfs/SCCS/s.m_sample.m 1.102 20/09/03 08:54:05 ';

persistent mytitldone
if isempty(mytitldone),           
   mytitldone = titlfunction(mytitl);
end

global titls

% Parameters for abbreviated form.
J = 1;
r = 1;
k = 1;
x = zeros(0, 1);

    % Conjugate prior for m of the form detailed in log_m_density.m
    
    M = size(x,1);
    y1 = x.^repmat(k,M,1);
    
    N = sum(x ~= Inf, 1);
    
    z1 = log(y1);
    z1(x == Inf) = 0;
    X1 = sum(z1, 1);
    
    z2 = repmat(r.^k, M, 1).*y1;
    z2(x == Inf) = 0;
    X2 = sum(z2, 1);
    
    z3 = repmat(log(r).*k, M, 1);
    z3(x == Inf) = 0;
    X3 = sum(z3, 1);
    
    
    b = m_b + N;
    a = m_a + m_b - X1 + X2 - X3 -b;
    y = zeros(1,J);
    
    for i = 1:J
       if isinf(a(i)), % Then the distribution is a all at infinity.
           y(i) = 1e100;
        else
           % Construct suitable input parameters for ars.
           points = [b(i)/(5*a(i)),b(i)/(2*a(i)), b(i)/a(i)];
           logf = @(x) log_m_density(x, a(i), b(i));
           gradlogf = @(x) log_m_derivative(x,a(i),b(i));
           y(i) = ars(logf, gradlogf, points, 0, Inf);
        end
    end
end

% Local Variables: 
% indent-line-function: indent-relative
% eval: (auto-fill-mode 0)
% End:
```
