## Supplementary material for "Elevated cerebrospinal fluid cytokine levels in tuberculous meningitis predict survival in response to dexamethasone": Software: makesynthdata.m

```
function [x, parmstrue] = makesynthdata(priors, Nx, ntype);
% Usage: [x, parmstrue] = makesynthdata(priors, Nx, ntype);
% Makes synthetic data.
% parmstrue is a cell array of size {size(Nx)} of which each cell contains a structure 
%  containing the fields ntype, indexing priors.typelist, and
%  e.g. parsmtrue.Gaussian if the type is Gaussian,
%  with parmstrue.Gaussian being a structure as defined in setpriors.m .
% It is assumed that seeding of random generators has already been done.
% The data output is a cell array of size {size(Nx)},
%  with each cell containing a column vector of length equal to the corresponding
%  element of Nx; each comes from the same distribution type.
% If ntype is passed and non-empty then it forces the choice of distribution type.

% Change Log:
%
%     1.1          01:may:20    rfs      First version.
%     1.3          01:may:20    rfs      Now appears to be working.
%     1.5          05:may:20    rfs      Added log-Gaussian.
%     1.7          05:may:20    rfs      Added Student.
%     1.10         06:may:20    rfs      Made able to make multiple groups of the same distribution type.
%     1.11         06:may:20    rfs      Added log-Student.
%     1.13         07:may:20    rfs      Added Gamma.
%     1.14         07:may:20    rfs      Added invGamma.
%     1.15         07:may:20    rfs      Added skewStudent.
%     1.16         07:may:20    rfs      Reparameterised the skewStudent.
%     1.17         07:may:20    rfs      Added logskewStudent.
%     1.18         08:may:20    rfs      ntype made a parameter to enable forcing it; remembered to 
%                                        consider mlow and mhigh.
%     1.23         09:may:20    rfs      Edited out entries that are out of permissible range 
%                                        according to the prior.
%     1.55         15:may:20    rfs      Fixed wrong inversion of nutrans.
%     1.68         25:may:20    rfs      Added Gammapow.
%     1.69         25:may:20    rfs      Fixed two nasty bugs generating data.
%     1.87         04:jun:20    rfs      Modified to make self-contained.

mytitl = ' /home/rfs/ramakrishnan/software/cytokines/rfs/SCCS/s.makesynthdata.m 1.102 20/09/03 08:54:05 ';

persistent mytitldone
if isempty(mytitldone),           
   mytitldone = titlfunction(mytitl);
end

global titls

if nargin < 3,
   ntype = [];
end

Ntypes = length(priors.typelist);

sz = size(Nx);
Nx = Nx(:);
Ngroups = length(Nx);

if isempty(ntype),
   ntype = randdiscreteE([1, 1], priors.Etypes, 1);
end

type = priors.typelist{ntype};

parmstrue = cell(Ngroups, 1);

for ngroup = 1 : Ngroups,
   
   switch type,
   
      case 'Gaussian',
         prior = priors.Gaussian;
         ss = prior.ss;
         ms = prior.ms;
         rs = ms * ss;
         Kmu = prior.Kmu;
         mumu = prior.mumu;
   
         strue = gamrnd(ms, 1 / rs, [1, 1]);
         mutrue = randn(1, 1) / sqrt(Kmu * strue) + mumu;

         x{ngroup} = randn(Nx(ngroup), 1) ./ sqrt(strue) + mutrue;

         parmstrueGaussian = struct('mutrue', mutrue, ...
                                    'strue', strue);
         parmstrue{ngroup} = struct('ntype', ntype, ...
                                    'Gaussian', parmstrueGaussian);      
   
      case 'logGaussian',
         prior = priors.logGaussian;
         ss = prior.ss;
         ms = prior.ms;
         rs = ms * ss;
         Kmu = prior.Kmu;
         mumu = prior.mumu;
   
         strue = gamrnd(ms, 1 / rs, [1, 1]);
         mutrue = randn(1, 1) / sqrt(Kmu * strue) + mumu;
   
         x{ngroup} = exp(randn(Nx(ngroup), 1) ./ sqrt(strue) + mutrue);
   
         parmstruelogGaussian = struct('mutrue', mutrue, ...
                                       'strue', strue);
         parmstrue{ngroup} = struct('ntype', ntype, ...
                                    'logGaussian', parmstruelogGaussian);      
   
      case 'Student',
         prior = priors.Student;
         mumu = prior.mumu;
         smu = prior.smu;
         am = prior.am;
         bm = prior.bm;
         mlow = prior.mlow;
         mhigh = prior.mhigh;
         ms = prior.ms;
         ss = prior.ss;
   
         mutrue = randn(1, 1) ./ sqrt(smu) + mumu;
         mtrue = m_sample(am, bm);
         mtrue = max(mlow, min(mtrue, mhigh));
         strue = gamrnd(ms, 1 ./ (ms * ss), [1, 1]);
         rtrue = mtrue * strue;
   
         x{ngroup} = mutrue + randn(Nx(ngroup), 1) ./ sqrt(gamrnd(mtrue, 1 ./ rtrue, [Nx(ngroup), 1]));
   
         parmstrueStudent = struct('mutrue', mutrue, ...
                                   'mtrue', mtrue, ...
                                   'strue', strue);
                                   
         parmstrue{ngroup} = struct('ntype', ntype, ...
                            'Student', parmstrueStudent);
   
      case 'logStudent',
         prior = priors.logStudent;
         mumu = prior.mumu;
         smu = prior.smu;
         am = prior.am;
         bm = prior.bm;
         mlow = prior.mlow;
         mhigh = prior.mhigh;
         ms = prior.ms;
         ss = prior.ss;
   
         mutrue = randn(1, 1) ./ sqrt(smu) + mumu;
         mtrue = m_sample(am, bm);
         mtrue = max(mlow, min(mtrue, mhigh));
         strue = gamrnd(ms, 1 ./ (ms * ss), [1, 1]);
         rtrue = mtrue * strue;
   
         x{ngroup} = exp(mutrue + randn(Nx(ngroup), 1) ./ sqrt(gamrnd(mtrue, 1 ./ rtrue, [Nx(ngroup), 1])));
   
         parmstruelogStudent = struct('mutrue', mutrue, ...
                                   'mtrue', mtrue, ...
                                   'strue', strue);
                                   
         parmstrue{ngroup} = struct('ntype', ntype, ...
                            'logStudent', parmstruelogStudent);
   
      case 'Gamma',
         prior = priors.Gamma;
         am = prior.am;
         bm = prior.bm;
         mlow = prior.mlow;
         mhigh = prior.mhigh;
         Km = prior.Km;
         ms = prior.ms;
         ss = prior.ss;
         rs = ms .* ss;
   
         mtrue = m_sample(am, bm);
         mtrue = max(mlow, min(mtrue, mhigh));
         strue = gamrnd(ms, 1 ./ rs, [1, 1]);
         rtrue = mtrue * strue;
   
         x{ngroup} = gamrnd(mtrue, 1 ./ rtrue, [Nx(ngroup), 1]);
   
         parmstrueGamma = struct('mtrue', mtrue, ...
                                 'strue', strue);
                                   
         parmstrue{ngroup} = struct('ntype', ntype, ...
                                    'Gamma', parmstrueGamma);
   
      case 'invGamma',
         prior = priors.invGamma;
         am = prior.am;
         bm = prior.bm;
         mlow = prior.mlow;
         mhigh = prior.mhigh;
         Km = prior.Km;
         ms = prior.ms;
         ss = prior.ss;
         rs = ms .* ss;
   
         mtrue = m_sample(am, bm);
         mtrue = max(mlow, min(mtrue, mhigh));
         strue = gamrnd(ms, 1 ./ rs, [1, 1]);
         rtrue = mtrue * strue;
   
         x{ngroup} = 1 ./ gamrnd(mtrue, 1 ./ rtrue, [Nx(ngroup), 1]);
   
         parmstrueinvGamma = struct('mtrue', mtrue, ...
                                    'strue', strue);
                                   
         parmstrue{ngroup} = struct('ntype', ntype, ...
                                    'invGamma', parmstrueinvGamma);
   
      case 'skewStudent',
         prior = priors.skewStudent;
         muxi = prior.muxi;
         sxi = prior.sxi;
         am = prior.am;
         bm = prior.bm;
         mlow = prior.mlow;
         mhigh = prior.mhigh;
         ms = prior.ms;
         ss = prior.ss;
         snu = prior.snu;
   
         xitrue = randn(1, 1) ./ sqrt(sxi) + muxi;
         mtrue = m_sample(am, bm);
         mtrue = max(mlow, min(mtrue, mhigh));
         strue = gamrnd(ms, 1 ./ (ms * ss), [1, 1]);
         rtrue = mtrue * strue;
         nutrue = randn(1, 1) ./ sqrt(snu);
         nucrtrue = nutransinv(nutrue);
         mutrue = xitrue - nutrue .* sqrt(rtrue) .* exp(gammaln(mtrue - 0.5) - gammaln(mtrue));
   
         x{ngroup} = mutrue + (randn(Nx(ngroup), 1) + nutrue) ./ sqrt(gamrnd(mtrue, 1 ./ rtrue, [Nx(ngroup), 1]));
   
         parmstrueskewStudent = struct('xitrue', xitrue, ...
                                       'mtrue', mtrue, ...
                                       'strue', strue, ...
                                       'nucrtrue', nucrtrue);
                                   
         parmstrue{ngroup} = struct('ntype', ntype, ...
                                    'skewStudent', parmstrueskewStudent);
   
      case 'logskewStudent',
         prior = priors.logskewStudent;
         muxi = prior.muxi;
         sxi = prior.sxi;
         am = prior.am;
         bm = prior.bm;
         mlow = prior.mlow;
         mhigh = prior.mhigh;
         ms = prior.ms;
         ss = prior.ss;
         snu = prior.snu;
   
         xitrue = randn(1, 1) ./ sqrt(sxi) + muxi;
         mtrue = m_sample(am, bm);
         mtrue = max(mlow, min(mtrue, mhigh));
         strue = gamrnd(ms, 1 ./ (ms * ss), [1, 1]);
         rtrue = mtrue * strue;
         nutrue = randn(1, 1) ./ sqrt(snu);
         nucrtrue = nutransinv(nutrue);
         mutrue = xitrue - nutrue .* sqrt(rtrue) .* exp(gammaln(mtrue - 0.5) - gammaln(mtrue));
   
         x{ngroup} = exp(mutrue + (randn(Nx(ngroup), 1) + nutrue) ./ sqrt(gamrnd(mtrue, 1 ./ rtrue, [Nx(ngroup), 1])));
   
         parmstruelogskewStudent = struct('xitrue', xitrue, ...
                                          'mtrue', mtrue, ...
                                          'strue', strue, ...
                                          'nucrtrue', nucrtrue);
                                   
         parmstrue{ngroup} = struct('ntype', ntype, ...
                                    'logskewStudent', parmstruelogskewStudent);
   
      case 'Gammapow',
         prior = priors.Gammapow;
         am = prior.am;
         bm = prior.bm;
         mlow = prior.mlow;
         mhigh = prior.mhigh;
         Km = prior.Km;
         ms = prior.ms;
         ss = prior.ss;
         rs = ms .* ss;
         Kk = prior.Kk;
         Nk = prior.Nk;
         ak = prior.Nk;
         bk = prior.bk;
         ck = prior.ck;

         mtrue = m_sample(am, bm);
         mtrue = max(mlow, min(mtrue, mhigh));
         strue = gamrnd(ms, 1 ./ rs, [1, 1]);
         ktrue = ksample(ak, bk, ck, Nk);
         rtrue = mtrue * strue ^ ktrue;

         x{ngroup} = gamrnd(mtrue, 1 ./ rtrue, [Nx(ngroup), 1]) .^ (1 ./ ktrue);
   
         parmstrueGammapow = struct('mtrue', mtrue, ...
                                    'strue', strue, ...
                                    'ktrue', ktrue);
                                   
         parmstrue{ngroup} = struct('ntype', ntype, ...
                                    'Gammapow', parmstrueGammapow);
   
      otherwise,
         error('Undefined type in makesynthdata');
   end

   ind = x{ngroup} < priors.xrange(1) | x{ngroup} > priors.xrange(2);
   if sum(ind) > 0,
      warning(sprintf('Of the desired %d entries for group %d, %d were out of range', ...
                      length(ind), ngroup, sum(ind)));
   end

   x{ngroup}(ind) = [];

   if Nx(ngroup) > 0 && isempty(x{ngroup}),
      warning(sprintf('There were no entries in range in group %d', ngroup));
   end

end % for ngroup = 1 : Ngroups

x = reshape(x, sz);
parmstrue = reshape(parmstrue, sz);

return;


% Local Variables: 
% indent-line-function: indent-relative
% eval: (auto-fill-mode 0)
% End:
```
