## Supplementary material for "Elevated cerebrospinal fluid cytokine levels in tuberculous meningitis predict survival in response to dexamethasone": Software: nutrans.m

```
function nu = nutrans(nucr);
% Usage: nu = nutrans(nucr);
% This function applies a transformation to nu.
% At present this is f(x) = (x ^ 3 + alpha * x) / (x ^ 2 + beta),
%  for internally set parameters alpha and beta.

% Change Log:
%
%     1.1          13:may:20    rfs      First version.

mytitl = ' /home/rfs/ramakrishnan/software/cytokines/rfs/SCCS/s.nutrans.m 1.102 20/09/03 08:54:05 ';

persistent mytitldone
if isempty(mytitldone),           
   mytitldone = titlfunction(mytitl);
end

global titls

% If you change these, you MUST also change the corresponding
%  settings in nutransderiv.m and nutransinv.m !
alpha = 100;
beta = 1000;

nu = (nucr .^ 3 + alpha * nucr) ./ (nucr .^ 2 + beta);

return;


% Local Variables: 
% indent-line-function: indent-relative
% eval: (auto-fill-mode 0)
% End:
```
