## Supplementary material for "Elevated cerebrospinal fluid cytokine levels in tuberculous meningitis predict survival in response to dexamethasone": Software: nutransinv.m

```
function nucr = nutransinv(nu);
% Usage: nucr = nutransinv(nu);
% This function applies the transformation to nu inverse to that in nutrans.m;
%  at present the latter is f(x) = (x ^ 3 + alpha * x) / (x ^ 2 + beta),
%  so this function simply uses the cubic equation formula to solve
%  for x given f(x).

% Change Log:
%
%     1.1          15:may:20    rfs      First version.
%     1.55         15:may:20    rfs      Version number harmonisation.

mytitl = ' /home/rfs/ramakrishnan/software/cytokines/rfs/SCCS/s.nutransinv.m 1.102 20/09/03 08:54:05 ';

persistent mytitldone
if isempty(mytitldone),           
   mytitldone = titlfunction(mytitl);
end

global titls

% If you change these, you MUST also change the corresponding
%  settings in nutrans.m and nutransinv.m !
alpha = 100;
beta = 1000;

% Solve the equation x ^ 3 - y * x ^ 2 + alpha * x - beta * y = 0 for x,
%  where x is nucr and nu is y.

p = -nu;
q = alpha;
r = - beta * nu;

% The equation is now x ^ 3 + p * x ^ 2 + q * x + r = 0;
%  see the Rubber Book for how to solve this.

a = q - p .^ 2 / 3;
b = (2 / 27) * p .^ 3 - p * q / 3 + r;

A = cbrt(- b / 2 + sqrt(b .^ 2 / 4 + a .^ 3 / 27));
B = cbrt(- b / 2 - sqrt(b .^ 2 / 4 + a .^ 3 / 27));

nucr = A + B - p / 3;

return;


function y = cbrt(x);
% Usage: y = cbrt(x);
% Calculates the real cube root of the real number x, ensuring that 
%  we don't get a complex number resulting.

y = sign(x) .* abs(x) .^ (1/3);

return;


% Local Variables: 
% indent-line-function: indent-relative
% eval: (auto-fill-mode 0)
% End:
```
