## Supplementary material for "Elevated cerebrospinal fluid cytokine levels in tuberculous meningitis predict survival in response to dexamethasone": Software: piecewise_exponential.m

```
function y = piecewise_exponential(points,vals,grads,bounds)
% Usage: y = piecewise_exponential(points,vals,grads,bounds)
% Samples from piecewise exponential (exponential components defined by
%  the equations of their log). 
% bounds: array of endpoints of the components.
% points, vals, grads: (points(i),vals(i)) lie on the logarithm of the ith
%  component, which has gradient grads(i).

% Change Log:
%
%     1.1          25:may:20    rfs      Taken from survival project piecewise_exponential.m v1.56 .
%     1.68         25:may:20    rfs      Version number harmonisation and adapted to use Esum and randdiscreteE.

mytitl = ' /home/rfs/ramakrishnan/software/cytokines/rfs/SCCS/s.piecewise_exponential.m 1.102 20/09/03 08:54:05 ';

persistent mytitldone
if isempty(mytitldone),           
   mytitldone = titlfunction(mytitl);
end

global titls

maxvals = max(vals);
if isinf(maxvals),
   if maxvals == -Inf,
      warning('All exp(vals) are zero in piecewise_exponential');
   else
      error('Infinite positive vals in piecewise_exponential');
   end
end
vals = vals - maxvals;

shifted_bounds = bounds(2:end);
bounds = bounds(1:end-1);
index = find(grads);
weights = zeros([1 length(bounds)]);
weights(index) = (1./grads(index)) .* (exp(grads(index).*(shifted_bounds(index)-points(index))+vals(index))-exp(grads(index).*(bounds(index)-points(index))+vals(index)));
zindex = find(~grads);
weights(zindex) = exp(vals(zindex)) .* (shifted_bounds(zindex) - bounds(zindex));

% RFS: Added this because we were getting NaN weights getting picked.
weights(isnan(weights)) = 0;

z = randdiscreteE([1, 1], -log(weights(:)));
u = rand;

bz = bounds(z);
sbz = shifted_bounds(z);

if grads(z) == Inf,
   y = [sbz, z];
elseif grads(z) == -Inf,
   y = [bz, z];
elseif isinf(bz) && grads(z) <= 0,
   % RFS: This shouldn't be able to happen, and should have been checked for in calling function.
   warning('Piecewise exponential called with unbounded density at left end');
   y = [sbz * (1 - 0.1 * sign(sbz)), z];
elseif isinf(sbz) && grads(z) >= 0,
   warning('Piecewise exponential called with unbounded density at right end');
   y = [bz * (1 + 0.1 * sign(bz)), z];
elseif grads(z) == 0,
   y = [bz.*u + sbz.*(1-u), z];
else
    a = [log(u) + (grads(z)*sbz), log(1-u)+grads(z)*bz];
    y = [(1/grads(z))*(-Esum(-a, 2)), z];
end

% RFS: I added this because otherwise numerical inaccuracy in the last two lines above
%      was leading to these conditions being violated.
if y(1) <= bz,
   y(1) = min([0.001 * sbz + 0.999 * bz, ...
               bz * (1 + sign(bz) * 0.001)]);
end
if y(1) >= sbz,
   y(1) = max([0.001 * bz + 0.999 * sbz, ...
              sbz * (1 - sign(sbz) * 0.001)]);
end

if any(isinf(y)) || any(isnan(y)) || y(1) < bounds(y(2)) || y(1) > shifted_bounds(y(2)),
   keyboard;
end

end

% Local Variables: 
% indent-line-function: indent-relative
% eval: (auto-fill-mode 0)
% End:
```
