## Supplementary material for "Elevated cerebrospinal fluid cytokine levels in tuberculous meningitis predict survival in response to dexamethasone": Software: piecewise_exponential_density.m

```
function y = piecewise_exponential_density(xx, points, vals, grads, bounds);
% Usage: y = piecewise_exponential_density(xx, points, vals, grads, bounds);
% Returns the unnormalised density of the piecewise exponential distribution
%  at the points xx, where:
%   bounds gives the changeover points from each piece to the next;
%   points is a list of x values one in each piece;
%   vals are the unnormalised log-densities at points;
%   grads are the gradients of log-densities at points.

% Change Log:
%
%     1.1          25:may:20    rfs      Taken from survival project piecewise_exponential_density.m v1.56 .
%     1.68         25:may:20    rfs      Version number harmonisation.

mytitl = ' /home/rfs/ramakrishnan/software/cytokines/rfs/SCCS/s.piecewise_exponential_density.m 1.102 20/09/03 08:54:05 ';

persistent mytitldone
if isempty(mytitldone),           
   mytitldone = titlfunction(mytitl);
end

global titls

Npieces = length(bounds) - 1;

vals = vals - max(vals);

for npiece = 1 : Npieces,
   inpiece = (bounds(npiece) <= xx) & (xx <= bounds(npiece + 1));
   y(inpiece) = exp(vals(npiece) + grads(npiece) * (xx(inpiece) - points(npiece)));
end

return;


% Local Variables: 
% indent-line-function: indent-relative
% eval: (auto-fill-mode 0)
% End:
```
