## Supplementary material for "Elevated cerebrospinal fluid cytokine levels in tuberculous meningitis predict survival in response to dexamethasone": Software: plotpriors.m

```
function plotpriors(priors);
% Usage: plotpriors(priors);

% Change Log:
%
%     1.1          01:may:20    rfs      First version.
%     1.3          01:may:20    rfs      Now appears to be working for Gaussian only.
%     1.7          05:may:20    rfs      Added Student.
%     1.11         06:may:20    rfs      Added logStudent.
%     1.14         07:may:20    rfs      Added Gamma.
%     1.30         11:may:20    rfs      Broadened plotting range in some cases.
%     1.68         25:may:20    rfs      Added Gammapow.
%     1.69         25:may:20    rfs      Various silly typos fixed.

mytitl = ' /home/rfs/ramakrishnan/software/cytokines/rfs/SCCS/s.plotpriors.m 1.102 20/09/03 08:54:05 ';

persistent mytitldone
if isempty(mytitldone),           
   mytitldone = titlfunction(mytitl);
end

global titls

% First plot the conjugate prior to the Gaussian.

prior = priors.Gaussian;
ss = prior.ss;
ms = prior.ms;
Kmu = prior.Kmu;
mumu = prior.mumu;

Nplot = 200;

muax = mumu - 5 * sqrt(ss) + [0 : Nplot] * 10 * sqrt(ss) / Nplot;
sax = (1 / ss) / 10 + [0 : Nplot] * (3 - 1/10) * (1 / ss) / Nplot;
[muax2, sax2] = ndgrid(muax, sax);

Egaussconj = - ms * log(ms * ss) + gammaln(ms) - (ms - 1) * log(sax2) + ms * ss * sax2 ...
             - 0.5 * log(Kmu * sax2 / (2 * pi)) + 0.5 * Kmu * sax2 .* (muax2 - mumu) .^ 2;

Pgaussconj = exp(- Egaussconj);

nfig = 0;

nfig = nfig + 1;
figure(nfig);
clf;
surf(muax2, sax2, Pgaussconj);
shading('interp');
view(0, 90);
colorbar;
xlabel('mu');
ylabel('s');
title('Conjugate prior to Gaussian');
axis([muax([1, end]), sax([1, end])]);

% Next plot the conjugate prior to the log-Gaussian.

prior = priors.logGaussian;
ss = prior.ss;
ms = prior.ms;
Kmu = prior.Kmu;
mumu = prior.mumu;

Nplot = 200;

muax = mumu - 5 * sqrt(ss) + [0 : Nplot] * 10 * sqrt(ss) / Nplot;
sax = (1 / ss) / 10 + [0 : Nplot] * (3 - 1/10) * (1 / ss) / Nplot;
[muax2, sax2] = ndgrid(muax, sax);

Eloggaussconj = - ms * log(ms * ss) + gammaln(ms) - (ms - 1) * log(sax2) + ms * ss * sax2 ...
             - 0.5 * log(Kmu * sax2 / (2 * pi)) + 0.5 * Kmu * sax2 .* (muax2 - mumu) .^ 2;

Ploggaussconj = exp(- Eloggaussconj);

nfig = nfig + 1;
figure(nfig);
clf;
surf(muax2, sax2, Ploggaussconj);
shading('interp');
view(0, 90);
colorbar;
xlabel('mu');
ylabel('s');
title('Conjugate prior to log-Gaussian');
axis([muax([1, end]), sax([1, end])]);

% Next plot the conjugate priors to the Student.

prior = priors.Student;

% First m.

mlow = prior.mlow;
mhigh = prior.mhigh;
deltam = 0.001;
mm = [mlow : deltam : mhigh].';
am = prior.am;
bm = prior.bm;
Km = prior.Km;
Emm = - log(Km) + (am + bm) .* mm - bm .* mm .* log(mm) + bm .* gammaln(mm);
Pmm = exp(- Emm);

nfig = nfig + 1;
figure(nfig);
plot(mm, Pmm);
xlabel('m');
ylabel('P(m)');
title('Conjugate prior to the Student for m');

% Then s.

ms = prior.ms;
ss = prior.ss;
s = [0.01 : 0.01 : 8 ./ ss].';
Es = - ms .* log(ms .* ss) + gammaln(ms) - (ms - 1) * log(s) + s .* (ms * ss);
Ps = exp(-Es);

nfig = nfig + 1;
figure(nfig);
plot(s, Ps);
xlabel('s');
ylabel('P(s)');
title('Conjugate prior to the Student for s');

% Then mu.

mumu = prior.mumu;
smu = prior.smu;
xxlow = mumu - 5 / sqrt(smu);
xxhigh = mumu + 5 / sqrt(smu);
deltax = (xxhigh - xxlow) / 100;
xx = [xxlow : deltax : xxhigh].';
Exx = - 0.5 * log(smu / (2 * pi)) + 0.5 * smu * (xx - mumu) .^ 2;
Pxx = exp(- Exx);

nfig = nfig + 1;
figure(nfig);
plot(xx, Pxx);
xlabel('mu');
ylabel('P(mu)');
title('Conjugate prior to the Student for mu');

% Next plot the conjugate priors to the logStudent.

prior = priors.logStudent;

% First m.

mlow = prior.mlow;
mhigh = prior.mhigh;
deltam = 0.001;
mm = [mlow : deltam : mhigh].';
am = prior.am;
bm = prior.bm;
Km = prior.Km;
Emm = - log(Km) + (am + bm) .* mm - bm .* mm .* log(mm) + bm .* gammaln(mm);
Pmm = exp(- Emm);

nfig = nfig + 1;
figure(nfig);
plot(mm, Pmm);
xlabel('m');
ylabel('P(m)');
title('Conjugate prior to the logStudent for m');

% Then s.

ms = prior.ms;
ss = prior.ss;
s = [0.001 : 0.001 : 8 ./ ss].';
Es = - ms .* log(ms .* ss) + gammaln(ms) - (ms - 1) * log(s) + s .* (ms * ss);
Ps = exp(-Es);

nfig = nfig + 1;
figure(nfig);
plot(s, Ps);
xlabel('s');
ylabel('P(s)');
title('Conjugate prior to the logStudent for s');

% Then mu.

mumu = prior.mumu;
smu = prior.smu;
xxlow = mumu - 5 / sqrt(smu);
xxhigh = mumu + 5 / sqrt(smu);
deltax = (xxhigh - xxlow) / 100;
xx = [xxlow : deltax : xxhigh].';
Exx = - 0.5 * log(smu / (2 * pi)) + 0.5 * smu * (xx - mumu) .^ 2;
Pxx = exp(- Exx);

nfig = nfig + 1;
figure(nfig);
plot(xx, Pxx);
xlabel('mu');
ylabel('P(mu)');
title('Conjugate prior to the logStudent for mu');

% Next plot the conjugate priors to the Gamma.

prior = priors.Gamma;

% First m.

mlow = prior.mlow;
mhigh = prior.mhigh;
deltam = 0.001;
mm = [mlow : deltam : mhigh].';
am = prior.am;
bm = prior.bm;
Km = prior.Km;
Emm = - log(Km) + (am + bm) .* mm - bm .* mm .* log(mm) + bm .* gammaln(mm);
Pmm = exp(- Emm);

nfig = nfig + 1;
figure(nfig);
plot(mm, Pmm);
xlabel('m');
ylabel('P(m)');
title('Conjugate prior to the Gamma for m');

% Then s.

ms = prior.ms;
ss = prior.ss;
s = [0.001 : 0.001 : 8 ./ ss].';
Es = - ms .* log(ms .* ss) + gammaln(ms) - (ms - 1) * log(s) + s .* (ms * ss);
Ps = exp(-Es);

nfig = nfig + 1;
figure(nfig);
plot(s, Ps);
xlabel('s');
ylabel('P(s)');
title('Conjugate prior to the Gamma for s');

% Next plot the conjugate priors to the invGamma.

prior = priors.invGamma;

% First m.

mlow = prior.mlow;
mhigh = prior.mhigh;
deltam = 0.001;
mm = [mlow : deltam : mhigh].';
am = prior.am;
bm = prior.bm;
Km = prior.Km;
Emm = - log(Km) + (am + bm) .* mm - bm .* mm .* log(mm) + bm .* gammaln(mm);
Pmm = exp(- Emm);

nfig = nfig + 1;
figure(nfig);
plot(mm, Pmm);
xlabel('m');
ylabel('P(m)');
title('Conjugate prior to the invGamma for m');

% Then s.

ms = prior.ms;
ss = prior.ss;
s = [0.01 : 0.01 : 8 ./ ss].';
Es = - ms .* log(ms .* ss) + gammaln(ms) - (ms - 1) * log(s) + s .* (ms * ss);
Ps = exp(-Es);

nfig = nfig + 1;
figure(nfig);
plot(s, Ps);
xlabel('s');
ylabel('P(s)');
title('Conjugate prior to the invGamma for s');

% Next plot the conjugate priors to the skewStudent.

prior = priors.skewStudent;

% First m.

mlow = prior.mlow;
mhigh = prior.mhigh;
deltam = 0.001;
mm = [mlow : deltam : mhigh].';
am = prior.am;
bm = prior.bm;
Km = prior.Km;
Emm = - log(Km) + (am + bm) .* mm - bm .* mm .* log(mm) + bm .* gammaln(mm);
Pmm = exp(- Emm);

nfig = nfig + 1;
figure(nfig);
plot(mm, Pmm);
xlabel('m');
ylabel('P(m)');
title('Conjugate prior to the skewStudent for m');

% Then s.

ms = prior.ms;
ss = prior.ss;
s = [0.01 : 0.01 : 8 ./ ss].';
Es = - ms .* log(ms .* ss) + gammaln(ms) - (ms - 1) * log(s) + s .* (ms * ss);
Ps = exp(-Es);

nfig = nfig + 1;
figure(nfig);
plot(s, Ps);
xlabel('s');
ylabel('P(s)');
title('Conjugate prior to the skewStudent for s');

% Then xi.

muxi = prior.muxi;
sxi = prior.sxi;
xxlow = muxi - 5 / sqrt(sxi);
xxhigh = muxi + 5 / sqrt(sxi);
deltax = (xxhigh - xxlow) / 100;
xx = [xxlow : deltax : xxhigh].';
Exx = - 0.5 * log(sxi / (2 * pi)) + 0.5 * sxi * (xx - muxi) .^ 2;
Pxx = exp(- Exx);

nfig = nfig + 1;
figure(nfig);
plot(xx, Pxx);
xlabel('xi');
ylabel('P(xi)');
title('Conjugate prior to the skewStudent for xi');

% Then nu.
snu = prior.snu;
xxlow = - 5 / sqrt(snu);
xxhigh = 5 / sqrt(snu);
deltax = (xxhigh - xxlow) / 100;
xx = [xxlow : deltax : xxhigh].';
Exx = - 0.5 * log(snu / (2 * pi)) + 0.5 * snu * xx .^ 2;
Pxx = exp(- Exx);

nfig = nfig + 1;
figure(nfig);
plot(xx, Pxx);
xlabel('nu');
ylabel('P(nu)');
title('Conjugate prior to the skewStudent for nu');

% Next plot the conjugate priors to the logskewStudent.

prior = priors.logskewStudent;

% First m.

mlow = prior.mlow;
mhigh = prior.mhigh;
deltam = 0.001;
mm = [mlow : deltam : mhigh].';
am = prior.am;
bm = prior.bm;
Km = prior.Km;
Emm = - log(Km) + (am + bm) .* mm - bm .* mm .* log(mm) + bm .* gammaln(mm);
Pmm = exp(- Emm);

nfig = nfig + 1;
figure(nfig);
plot(mm, Pmm);
xlabel('m');
ylabel('P(m)');
title('Conjugate prior to the logskewStudent for m');

% Then s.

ms = prior.ms;
ss = prior.ss;
s = [0.01 : 0.01 : 8 ./ ss].';
Es = - ms .* log(ms .* ss) + gammaln(ms) - (ms - 1) * log(s) + s .* (ms * ss);
Ps = exp(-Es);

nfig = nfig + 1;
figure(nfig);
plot(s, Ps);
xlabel('s');
ylabel('P(s)');
title('Conjugate prior to the logskewStudent for s');

% Then xi.

muxi = prior.muxi;
sxi = prior.sxi;
xxlow = muxi - 5 / sqrt(sxi);
xxhigh = muxi + 5 / sqrt(sxi);
deltax = (xxhigh - xxlow) / 100;
xx = [xxlow : deltax : xxhigh].';
Exx = - 0.5 * log(sxi / (2 * pi)) + 0.5 * sxi * (xx - muxi) .^ 2;
Pxx = exp(- Exx);

nfig = nfig + 1;
figure(nfig);
plot(xx, Pxx);
xlabel('xi');
ylabel('P(xi)');
title('Conjugate prior to the logskewStudent for xi');

% Then nu.
snu = prior.snu;
xxlow = - 5 / sqrt(snu);
xxhigh = 5 / sqrt(snu);
deltax = (xxhigh - xxlow) / 100;
xx = [xxlow : deltax : xxhigh].';
Exx = - 0.5 * log(snu / (2 * pi)) + 0.5 * snu * xx .^ 2;
Pxx = exp(- Exx);

nfig = nfig + 1;
figure(nfig);
plot(xx, Pxx);
xlabel('nu');
ylabel('P(nu)');
title('Conjugate prior to the logskewStudent for nu');

% Next plot the conjugate priors to the Gamma powered.

prior = priors.Gammapow;

% First m.

mlow = prior.mlow;
mhigh = prior.mhigh;
deltam = 0.001;
mm = [mlow : deltam : mhigh].';
am = prior.am;
bm = prior.bm;
Km = prior.Km;
Emm = - log(Km) + (am + bm) .* mm - bm .* mm .* log(mm) + bm .* gammaln(mm);
Pmm = exp(- Emm);

nfig = nfig + 1;
figure(nfig);
plot(mm, Pmm);
xlabel('m');
ylabel('P(m)');
title('Conjugate prior to the Gammapow for m');

% Then s.

ms = prior.ms;
ss = prior.ss;
s = [0.001 : 0.001 : 8 ./ ss].';
Es = - ms .* log(ms .* ss) + gammaln(ms) - (ms - 1) * log(s) + s .* (ms * ss);
Ps = exp(-Es);

nfig = nfig + 1;
figure(nfig);
plot(s, Ps);
xlabel('s');
ylabel('P(s)');
title('Conjugate prior to the Gammapow for s');

% Then k.

Nk = prior.Nk;
ak = prior.ak;
bk = prior.bk;
ck = prior.ck;
Kk = prior.Kk;
klow = prior.klow;
khigh = prior.khigh;
k = [klow : 0.001 : khigh].';
Ek = - log(Kk) - ak .* log(k);
for n = 1 : Nk,
   Ek = Ek - k .* ck(n) .* log(bk(n)) + ck(n) .* bk(n) .^ k; 
end
Pk = exp(-Ek);

nfig = nfig + 1;
figure(nfig);
plot(k, Pk);
xlabel('k');
ylabel('P(k)');
title('Conjugate prior to the Gammapow for k');

return;


% Local Variables: 
% indent-line-function: indent-relative
% eval: (auto-fill-mode 0)
% End:
```
