## Supplementary material for "Elevated cerebrospinal fluid cytokine levels in tuberculous meningitis predict survival in response to dexamethasone": Software: plotpriorsamples.m

```
function plotpriorsamples(priors, Nsamples, ntype, lognotlin, lowx, highx, coloured);
% Usage: plotpriorsamples(priors, Nsamples, ntype, lognotlin, lowx, highx, coloured);
% This function plots Nsamples samples from the prior.
% If Nsamples is negative, it instead takes one sample from the prior,
%  and draws a random sample of size abs(Nsamples) from it, and makes 
%  a histogram of that. Note that such samples will automatically be 
%  edited to only include those in the prior's admissible range.
% If ntype is empty the type is chosen at random.
% x axis range is from lowx to highx.
% If coloured is 1 then a different colour or line style is used for each type,
%  else all are the same blue solid line.

% Change Log:
%
%     1.1          08:may:20    rfs      First version.
%     1.18         08:may:20    rfs      Now working.
%     1.26         09:may:20    rfs      Option added to make histograms instead.
%     1.76         26:may:20    rfs      Added legend.      
%     1.77         26:may:20    rfs      Fixed bug in figure title.

mytitl = ' /home/rfs/ramakrishnan/software/cytokines/rfs/SCCS/s.plotpriorsamples.m 1.102 20/09/03 08:54:05 ';

persistent mytitldone
if isempty(mytitldone),           
   mytitldone = titlfunction(mytitl);
end

global titls

if nargin < 3,
   ntype = [];
end
if nargin < 4,
   lognotlin = [];
end
if nargin < 5,
   lowx = [];
end
if nargin < 6,
   highx = [];
end
if nargin < 7,
   coloured = [];
end

if isempty(lognotlin),
   lognotlin = 1;
end
if isempty(lowx),
   lowx = priors.xrange(1);
end
if isempty(highx),
   highx = priors.xrange(2);
end
if isempty(coloured),
   coloured = ~isscalar(ntype);
end

Ntypes = length(priors.typelist);

if coloured,
   linestyles = {'g--', 'b--', 'r--', 'm--', 'c-', 'c--', 'r-', 'b-', 'm-'};
else
   linestyles = repmat({'b-'}, [1, Ntypes]);
end
      
if Nsamples < 0,
   Nhistsamples = abs(Nsamples);
   Nsamples = 1;
else
   Nhistsamples = 0;
end

Nx = 200;

if lognotlin,
   xx = exp(log(lowx) + [0 : Nx].' * (log(highx) - log(lowx)) / Nx);
else
   xx = lowx + [0 : Nx].' * (highx - lowx) / Nx;
end

Eyy = NaN(Nx + 1, Nsamples);
ntypes = NaN(Nsamples);

% Make the samples.
for nsample = 1 : Nsamples,

    [histsamples, parmstrue] = makesynthdata(priors, Nhistsamples, ntype);
    histsamples = histsamples{1};

    ntypes(nsample) = parmstrue{1}.ntype;
    theseparms = getfield(parmstrue{1}, priors.typelist{ntypes(nsample)});
    theseparms = struct2cell(theseparms);
    params = cell(length(theseparms) + 1, 1);
    [params{:}] = ndgrid(theseparms{:}, xx);
    Eyy(:, nsample) = squeeze(feval(priors.Efns{ntypes(nsample)}, params, [], 1));

end % for nsample = 1 : Nsamples

if Nhistsamples == 0,
   % Plot the samples.
   clf;
   hold on;
   for nsample = 1 : Nsamples,
      if lognotlin,
         plot(log(xx), exp(-Eyy(:, nsample) + log(xx)), linestyles{ntypes(nsample)});
      else
         plot(xx, exp(-Eyy(:, nsample)), linestyles{ntypes(nsample)});
      end
   end
   na = NaN;
   h = NaN(Ntypes, 1);
   for thisntype = 1 : Ntypes,
      h(thisntype) = plot(na, na, linestyles{thisntype});
   end
   hold off;
   legend(h, priors.typelist, 'Location', 'NorthWest');
   if isempty(ntype),
      title('Samples from prior');
   else
      title(sprintf('%s samples from prior', priors.typelist{ntypes(nsample)}));
   end
   if lognotlin,
      xlabel('log(x)');
      ylabel('P(log(x))');
   else
      xlabel('x');
      ylabel('P(x)');
   end
else % i.e. if we're plotting a histogram

   Nhistbins = ceil(Nhistsamples / 3);
   if lognotlin,
      xxhist = log(lowx) + [0 : Nhistbins] * (log(highx) - log(lowx)) / Nhistbins;
   else
      xxhist = lowx + [0 : Nhistbins] * (highx - lowx) / Nhistbins;
   end

   clf;
   hold on;
   if lognotlin,
      hist(log(histsamples), xxhist);
   else
      hist(histsamples, xxhist);
   end

   deltax = xxhist(2) - xxhist(1);

   if lognotlin,
      plot(log(xx), exp(-Eyy(:, nsample) + log(xx)) * Nhistsamples * deltax, linestyles{ntypes(nsample)});
   else
      plot(xx, exp(-Eyy(:, nsample)) * Nhistsamples * deltax, linestyles{ntypes(nsample)});
   end
   hold off;

end % if Nhistsamples == 0 or not

return;


% Local Variables: 
% indent-line-function: indent-relative
% eval: (auto-fill-mode 0)
% End:
```
