## Supplementary material for "Elevated cerebrospinal fluid cytokine levels in tuberculous meningitis predict survival in response to dexamethasone": Software: progammaprecalcs.m

```
function priorstruct = progammaprecalcs(priorstruct);
% Usage: priorstruct = progammaprecalcs(priorstruct);
% This function does some precalculations for proGamma distributions to save 
%  time later.

% Change Log:
%
%     1.1          07:may:20    rfs      First version.

mytitl = ' /home/rfs/ramakrishnan/software/cytokines/rfs/SCCS/s.progammaprecalcs.m 1.102 20/09/03 08:54:05 ';

persistent mytitldone
if isempty(mytitldone),           
   mytitldone = titlfunction(mytitl);
end

global titls

% First the normalisation constant.

deltam = 0.001;
mm = [priorstruct.mlow : deltam : priorstruct.mhigh].';
Emm = (priorstruct.am + priorstruct.bm) .* mm - priorstruct.bm .* mm .* log(mm) + priorstruct.bm .* gammaln(mm);
Pmm = exp(- Emm);
priorstruct.Km = 1 ./ (deltam * sum(Pmm));

return;


% Local Variables: 
% indent-line-function: indent-relative
% eval: (auto-fill-mode 0)
% End:
```
