## Supplementary material for "Elevated cerebrospinal fluid cytokine levels in tuberculous meningitis predict survival in response to dexamethasone": Software: propowprecalcs.m

```
function priorstruct = propowprecalcs(priorstruct);
% Usage: priorstruct = propowprecalcs(priorstruct);
% This function does some precalculations for the prior on Gammapow k to save 
%  time later.

% Change Log:
%
%     1.1          25:may:20    rfs      First version.

mytitl = ' /home/rfs/ramakrishnan/software/cytokines/rfs/SCCS/s.propowprecalcs.m 1.102 20/09/03 08:54:05 ';

persistent mytitldone
if isempty(mytitldone),           
   mytitldone = titlfunction(mytitl);
end

global titls

% First the normalisation constant.

deltak = 0.0001;
kk = [priorstruct.klow : deltak : priorstruct.khigh].';
Ekk = - priorstruct.ak .* log(kk);
for n = 1 : priorstruct.Nk,
   Ekk = Ekk - kk .* priorstruct.ck(n) .* log(priorstruct.bk(n)) + priorstruct.ck(n) .* priorstruct.bk(n) .^ kk;
end
Pkk = exp(- Ekk);
priorstruct.Kk = 1 ./ (deltak * sum(Pkk));

return;


% Local Variables: 
% indent-line-function: indent-relative
% eval: (auto-fill-mode 0)
% End:
```
