## Supplementary material for "Elevated cerebrospinal fluid cytokine levels in tuberculous meningitis predict survival in response to dexamethasone": Software: randdiscreteE.m

```
function [out] = randdiscreteE(sz, Es, nowarning);
% Usage: [out] = randdiscreteE(sz, Es, nowarning);
% Draws random samples from a finite discrete distribution.
% The output array out has size sz.
% The probability of each outcome is given by exp(-Es),
%  which need not sum to 1 over the appropriate vectors as it
%  will automatically be normalised before use. The possible 
%  outcomes are 1, 2, ..., M, and Es(m, K) gives minus the log of
%  the (unnormalised) probability of out(K) having value m.
% If all elements of such a vector are zero then the output distribution is uniform;
%  under such circumstances a warning is given unless nowarning
%  is passed and non-zero.
% probs should either be of size [M, 1] or of size [M, sz].

% Change Log:
%
%     1.1          02:may:20    rfs  First version.
%     1.4          02:may:20    rfs  Version number harmonisation.
%
%  /home/rfs/ramakrishnan/software/cytokines/rfs/SCCS/s.randdiscreteE.m 1.102 20/09/03 08:54:05

if nargin < 2,
   error('Not enough arguments');
end
if nargin < 3,
   nowarning = [];
end

if isempty(nowarning),
   nowarning = 0;
end

M = size(Es, 1);
N = prod(sz);

if M == 0,
   error('randdiscreteE cannot take samples from the empty set');
end

ind = find(sz ~= 1);
if isempty(ind),
   sz = [1];
else
   sz = sz(1 : ind(end));
end

if prod(size(Es)) == M,
   Es = repmat(Es(:), [1, sz]);
end

if length(size(Es)) ~= length([M, sz]),
   error('Es is neither a simple vector nor of the same dimensionality as sz');
end
if any(size(Es) ~= [M, sz]),
   error('Es is neither a simple vector nor of size [M, sz] for some M');
end

Es = reshape(Es, [M, N]);

% Normalise.
minEs = min(Es, [], 1);
Es = Es - reshape(minEs(ones(size(Es, 1), 1), :), size(Es));
probs = exp(- Es);
probsum = sum(probs, 1);
ind = find(probsum(:) == 0);
if ~isempty(ind),
   if ~nowarning,
      warning('randdiscreteE called with zero vector of probabilities');
   end
   probsum(ind) = 1;
end
probs = probs ./ repmat(probsum, [M, 1]);
probs(:, ind) = 1 / M;

probs = cumsum(probs, 1);

out = max(1, min(M, 1 + sum(repmat(rand(1, N), [M, 1]) > probs, 1)));

out = reshape(out, [sz, 1]);

return;


% Local Variables: 
% indent-line-function: indent-relative
% eval: (auto-fill-mode 0)
% End:
```
