## Supplementary material for "Elevated cerebrospinal fluid cytokine levels in tuberculous meningitis predict survival in response to dexamethasone": Software: runrealdata.m

```
function runrealdata(cyks, ncyks, tcount, maxtcount, selection, savesetid, savedir, ...
                     allcompprobslin, allcompprobslog, allcompprobsgeom, Etot);
% Usage: runrealdata(cyks, ncyks, tcount, maxtcount, selection, savesetid, savedir, ...
%                    allcompprobslin, allcompprobslog, allcompprobsgeom, Etot);
% This function applies comparegroups to the three genotypes from 
%  each cytokine number in the vector ncyks, selecting to
%  use in each group those with a 1 in the logical array selection,
%  excluding those with NaN values.
% cyks should be of size [Npatients, Ncyks] and contains the cytokine
%  measured values (in pg/ml).
% ncyks is a vector of arbitrary length asking for the question to be
%  done on each cytokine number in the vector.
% tcount is a vector of size [Npatients, 1] containing the number 
%  of T alleles for that patient; each entry is between 0 and 2;
%  alternatively it allocates patients to groups to be compared, numbered from 
%  0 to Ngroups - 1 .
% If the maximum value of tcount wanted is greater than the maximum value represented,
%  set maxtcount to be the maximum value wanted; it defaults to max(tcount).
% selection is a logical array of size [Npatients, 1] saying which are to
%  be included in the data sample.
% If allcompprobslin, allcompprobslog, allcompprobsgeom, and Etot are passed in then 
%  they should be from another previous run with fewer types, and only those 
%  elements which are NaN will be recalculated.

% Change Log:
%
%     1.1          15:may:20    rfs      First version, not yet debugged.
%     1.58         15:may:20    rfs      Version number harmonisation.
%     1.59         15:may:20    rfs      Trivial bugs fixed.
%     1.60         16:may:20    rfs      Typo leading to massive memory usage fixed; diary added.
%     1.61         16:may:20    rfs      Typo leading to crash after doing first ncyk fixed.
%     1.62         17:may:20    rfs      Fixed name of save file.
%     1.63         18:may:20    rfs      Now tells you which ncyk it's working on.
%     1.68         25:may:20    rfs      Now gets Ntypes from the prior.
%     1.72         26:may:20    rfs      Now takes in previously calculated values; not yet debugged.
%     1.84         31:may:20    rfs      Now sets Ngroups to be max(tcount) + 1.
%     1.85         03:jun:20    rfs      Added geometric means, not yet debugged.
%     1.86         04:jun:20    rfs      Bugs in above fixed, appears to be working.
%     1.95         04:jul:20    rfs      Added catcher for empty x.
%     1.96         04:jul:20    rfs      Fixed stupid typo.
%     1.99         18:aug:20    rfs      Made temporary savefilenames dependent on savesetid.
%     1.100        31:aug:20    rfs      Added maxtcount option.
%     1.101        02:sep:20    rfs      Fixed stupid typo in above.
%     1.102        03:sep:20    rfs      Reordered parameters to put maxtcount after tcount.

mytitl = ' /home/rfs/ramakrishnan/software/cytokines/rfs/SCCS/s.runrealdata.m 1.102 20/09/03 08:54:05 ';

persistent mytitldone
if isempty(mytitldone),           
   mytitldone = titlfunction(mytitl);
end

global titls

if nargin < 6,
   savesetid = '';
end
if nargin < 7,
   savedir = '';
end
if nargin < 8,
   allcompprobslin = [];
end
if nargin < 9,
   allcompprobslog = [];
end
if nargin < 10,
   allcompprobsgeom = [];
end
if nargin < 11,
   Etot = [];
end

if isempty(savedir),
   savedir = '.';
end
if isempty(maxtcount),
   maxtcount = max(tcount(:));
end

if ~isempty(savesetid),
   diary(sprintf('%s/matlabout.%s', savedir, savesetid));
end

cyks = cyks(selection, :);
tcount = tcount(selection);

Npatients = size(cyks, 1);
Ngroups = maxtcount + 1;
Nncyks = length(ncyks);
temppriors = setpriors(1);
Ntypes = length(temppriors.typelist);

Etypes = NaN(Ntypes, Nncyks);
compprobslin = NaN(Ngroups, Ngroups, Nncyks);
compprobslog = NaN(Ngroups, Ngroups, Nncyks);
compprobsgeom = NaN(Ngroups, Ngroups, Nncyks);

if isempty(Etot),
   Etot = NaN(Ntypes, Ngroups, Nncyks);
end
if isempty(allcompprobslin),
   allcompprobslin = NaN(Ngroups, Ngroups, Ntypes, Nncyks);
end
if isempty(allcompprobslog),
   allcompprobslog = NaN(Ngroups, Ngroups, Ntypes, Nncyks);
end
if isempty(allcompprobsgeom),
   allcompprobsgeom = NaN(Ngroups, Ngroups, Ntypes, Nncyks);
end

oldNtypes = size(Etot, 1);
if oldNtypes < Ntypes,
   extraNtypes = Ntypes - oldNtypes;
   Etot = cat(1, Etot, NaN(extraNtypes, Ngroups, Nncyks));
end

oldNtypes = size(allcompprobslin, 3);
if oldNtypes < Ntypes,
   extraNtypes = Ntypes - oldNtypes;
   allcompprobslin = cat(3, allcompprobslin, NaN(Ngroups, Ngroups, extraNtypes, Nncyks));
end

oldNtypes = size(allcompprobslog, 3);
if oldNtypes < Ntypes,
   extraNtypes = Ntypes - oldNtypes;
   allcompprobslog = cat(3, allcompprobslog, NaN(Ngroups, Ngroups, extraNtypes, Nncyks));
end

oldNtypes = size(allcompprobsgeom, 3);
if oldNtypes < Ntypes,
   extraNtypes = Ntypes - oldNtypes;
   allcompprobsgeom = cat(3, allcompprobsgeom, NaN(Ngroups, Ngroups, extraNtypes, Nncyks));
end

for nncyk = 1 : Nncyks,
   ncyk = ncyks(nncyk);

   fprintf('Starting on nncyk %d, ncyk %d.\n', nncyk, ncyk);

   thesecyks = cell(Ngroups, 1);

   for ngroup = 1 : Ngroups,
      rawcyks = cyks(tcount == ngroup - 1, ncyk);
      thesecyks{ngroup} = rawcyks(~isnan(rawcyks));
   end

   allthesecyks = cat(1, thesecyks{:});
   meancyk = exp(mean(log(allthesecyks)));

   if isnan(meancyk) || isinf(meancyk),
      meancyk = 1;
   end
   priors = setpriors(meancyk);

   Ngridlin = 150;
   xmeangridlin = (1e-2 + 1e2 * [0 : Ngridlin].' / Ngridlin) * meancyk;

   Ngridlog = 150;
   xmeangridlog = exp(log(meancyk) - 6 + 12 * [0 : Ngridlog].' / Ngridlog);

   nfig = 1;
   
   [thisEtypes, thiscompprobslin, thiscompprobslog, thiscompprobsgeom, ...
    thisallcompprobslin, thisallcompprobslog, thisallcompprobsgeom, thisEtot] ...
      = comparegroups(thesecyks, priors, xmeangridlin, xmeangridlog, [], nfig, ...
                      allcompprobslin(:, :, :, nncyk), allcompprobslog(:, :, :, nncyk), ...
                      allcompprobsgeom(:, :, :, nncyk), Etot(:, :, nncyk));

   Etypes(:, nncyk) = thisEtypes;
   compprobslin(:, :, nncyk) = thiscompprobslin;
   compprobslog(:, :, nncyk) = thiscompprobslog;
   compprobsgeom(:, :, nncyk) = thiscompprobsgeom;
   allcompprobslin(:, :, :, nncyk) = thisallcompprobslin;
   allcompprobslog(:, :, :, nncyk) = thisallcompprobslog;
   allcompprobsgeom(:, :, :, nncyk) = thisallcompprobsgeom;
   Etot(:, :, nncyk) = thisEtot;

   if ~isempty(savesetid),
      quicksave([savesetid, sprintf('.%d', ncyk)], savedir); % Note saved under cyk number, not number in ncyks.
   end

   close all

   save(sprintf('tempsavecyks.%s.mat', savesetid));

end % for nncyk = 1 : Nncyks

if ~isempty(savesetid),
   savefilename = sprintf('%s/save.%s.mat', savedir, savesetid);
   save(savefilename);
   if isunix,
      unix(sprintf('chmod a-w %s', savefilename));
   end
end

diary off;

return;


% Local Variables: 
% indent-line-function: indent-relative
% eval: (auto-fill-mode 0)
% End:
```
