## Supplementary material for "Elevated cerebrospinal fluid cytokine levels in tuberculous meningitis predict survival in response to dexamethasone": Software: setpriors.m

```
function priors = setpriors(xcentre, xwidth);
% Usage: priors = setpriors(xcentre, xwidth);
% This function sets the priors for all distribution types considered.
% All priors are "multiplied" by xcentre, typically around 1 ng/litre.
% xwidth is 1 at baseline, and increases if broader distributions are wanted.

% Change Log:
%
%     1.1          01:may:20    rfs      First version.
%     1.3          01:may:20    rfs      Now appears to be working with Gaussian only.
%     1.4          02:may:20    rfs      Now uses Etypes instead of ptypes.
%     1.5          05:may:20    rfs      Added log-Gaussian, seems to be working.
%     1.7          05:may:20    rfs      Added Student.
%     1.11         06:may:20    rfs      Added logStudent.
%     1.12         07:may:20    rfs      logStudent now working.
%     1.13         07:may:20    rfs      Added Gamma.
%     1.14         07:may:20    rfs      Added invGamma.
%     1.15         07:may:20    rfs      Added skewStudent.
%     1.16         07:may:20    rfs      Reparameterised the skewStudent.
%     1.17         07:may:20    rfs      Added logskewStudent.
%     1.18         08:may:20    rfs      Adjusted various parameters.
%     1.19         08:may:20    rfs      Changed from nck to xcentre parameter for scaling.
%     1.20         08:may:20    rfs      xwidth introduced.
%     1.21         08:may:20    rfs      Default width doubled.
%     1.30         11:may:20    rfs      xrange restored to [-6,6] independent of d,
%                                        and many other changes arising from initial 
%                                        formal test run.
%     1.31         11:may:20    rfs      Ngridmax and shrinklimit set here now; a few more changes to 
%                                        priors to avoid needing editing.
%     1.32         11:may:20    rfs      Now sets Ngridmax to an array so as to determine
%                                        which parameters get finer grids.
%     1.36         11:may:20    rfs      Initial bounds for m set wider.
%     1.46         13:may:20    rfs      Reparameterised nu with its cube root nucr.
%     1.56         15:may:20    rfs      Reparameterised nu, fixed bug not using nutrans,
%                                        and adjusted priors as a result.
%     1.57         15:may:20    rfs      Reviewed priors and adjusted again.
%     1.64         20:may:20    rfs      Adjusted priors to be more similar from distribution to distribution,
%                                        in the light of review when doing methods section.
%     1.68         25:may:20    rfs      Added Gammapow.
%     1.69         25:may:20    rfs      Adjust prior for Gammapow.
%     1.73         26:may:20    rfs      Removed block on types 7 and 8.
%     1.88         12:jun:20    rfs      Added figure offsets for the different families.
%     1.91         19:jun:20    rfs      Now handles info from Thuong 2017 supplementary.
%     1.93         27:jun:20    rfs      Permanently set prior prob of non-log skew Student to 0
%                                        and fixed figure counting resulting.

mytitl = ' /home/rfs/ramakrishnan/software/cytokines/rfs/SCCS/s.setpriors.m 1.102 20/09/03 08:54:05 ';

persistent mytitldone
if isempty(mytitldone),           
   mytitldone = titlfunction(mytitl);
end

global titls

if nargin < 1,
   xcentre = [];
end
if nargin < 2,
   xwidth = [];
end

if isempty(xcentre),
   xcentre = 1;
end
if isempty(xwidth),
   xwidth = 1;
end

% The following must all be in matching order.
typelist = {'Gaussian', 'logGaussian', 'Student', 'logStudent', ...
            'Gamma', 'invGamma', 'skewStudent', 'logskewStudent', 'Gammapow'};
Ntypes = length(typelist);
posonly = [0, 1, 0, 1, 1, 1, 0, 1, 1].';
Efns = {@Egauss, @Eloggauss, @Estudent, @Elogstudent, ...
        @Egamma, @Einvgamma, @Eskewstudent, @Elogskewstudent, @Egammapow};
ditherfns = {@dithergau, @ditherlg, @ditherstud, @ditherls, @dithergam, ...
             @ditherig, @ditherss, @ditherlss, @dithergp};
shrinklimit = repmat(0.01, [1, Ntypes]);
Nparams = [2, 2, 3, 3, 2, 2, 4, 4, 3].';
Ngridmax = cell(Ntypes, 1);
for ntype = 1 : Ntypes,
   Ngridmax{ntype} = repmat(40, [Nparams(ntype), 1]);
end
for ntype = 7 : 8,
   Ngridmax{ntype}(2) = 20; % Exception for (log)skewStudent as we don't have quite enough memory.
end
Ngridmax{9} = [160; 40; 160]; % Exception for Gammapow as not fine enough as it stood.

Etypes = repmat(log(Ntypes), [Ntypes, 1]);

% Add these lines if you want to exclude (log-)skew-Students.
% Etypes = repmat(log(Ntypes - 2), [Ntypes, 1]);
% Etypes(7:8) = Inf;

% Add these lines if you want to exclude just the non-log skew Student.
Etypes = repmat(log(Ntypes - 1), [Ntypes, 1]);
Etypes(7) = Inf;

% Calculate the subfigure numbers in which the various parameters are plotted during runrealdata.m .
Nsubfigures = NaN(Ntypes, 1);
for ntype = 1 : Ntypes,
   if ~isinf(Etypes(ntype)),
      Nsubfigures(ntype) = 2 + nchoosek(Nparams(ntype), 2);
   else
      Nsubfigures(ntype) = 0;
   end
end
groupstep = sum(Nsubfigures);
familyoffsets = cumsum(Nsubfigures);
familyoffsets = [0; familyoffsets(1 : Ntypes - 1)] + 1;

% The conjugate prior for mu and s (=1/sigma^2) jointly on a Gaussian is given by:
%  P(s) = ((ms * ss) ^ ms / Gamma(ms)) * s ^ (ms - 1) * exp(-ms * ss * s)
%  P(mu|s) = sqrt(Kmu * s / (2 * pi)) * exp(-0.5 * Kmu * s * (mu - mumu) ^ 2),
% and similarly for the log-Gaussian.

% The Student will be parameterised by mu and m and s, where r = m * s.
% The conjugate prior for mu on a Student is Gaussian with parameters mumu and smu.
% The conjugate prior for m on a Student is proGamma given by:
%  P(m | a, b) = K * exp(-(a + b) .* m) .* m .^ (b .* m) ./ gamma(m) .^ b, 
%  with a > 0 and b > -1, and K is the integration constant;
%  a and b will in the code below be denoted am and bm;
%  to ensure numerical integrability, the range of m will be limited to mlow to mhigh.
% The conjugate prior for s on a Student is Gamma with parameters ms and ss, with rs = ms * ss.
% Similarly for the log-Student.

% For the skewStudent (and similarly the log-skewStudent) see Eskewstudent.m for
%  the adjusted parameterisation; m and s have the same prior families as for the Student,
%  while xi is Gaussian with mean muxi and scale sxi,
%  and nu is Gaussian with zero mean and scale snu, but parameterised as nu = nutrans(nucr).

% The Gamma will be parameterised by m and s, where r = m * s.
% The conjugate prior for m on a Gamma is proGamma given by:
%  P(m | a, b) = K * exp(-(a + b) .* m) .* m .^ (b .* m) ./ gamma(m) .^ b, 
%  with a > 0 and b > -1, and K is the integration constant;
%  a and b will in the code below be denoted am and bm;
%  to ensure numerical integrability, the range of m will be limited to mlow to mhigh.
% The conjugate prior for s on a Gamma is Gamma with parameters ms and ss, with rs = ms * ss.
% Similarly for the inverse Gamma.

% Gammapow denotes the distribution such that for some positive power k (a parameter),
%  x .^ k is Gamma distributed, i.e. with r = m * s ^ k,
%  P(x | m, s, k) = (r ^ m / Gamma(m)) * x ^ (m * k - 1) * exp(-r * x ^ k).
% Parameters are m, s, and k, which must all be positive (the version with k negative
%  has been excluded as inverse-Gamma fit is so bad).
% The priors on m and r are as for the Gamma distribution.
% The prior on k is 
%  P(k | N, a, b, c) = K * k ^ a * prod(b .^ (k .* c) * exp(-c .* b .^ k))
%  where N is a non-negative integer, a is a positive real, and b and c are vectors
%  of positive reals of size [N, 1], and K is a normalisation constant.

c = xcentre;
d = 2 * xwidth;

% xrange gives an absolute limit on permissible data values.
xrange = [exp(-6), exp(+6)] * c;

gaussian = struct('ss', 100 * d .^ 2 * c .^ 2, 'ms', 4, 'Kmu', 0.2, 'mumu', 150 * c, ...
                  'bounds', [[0.1; 5] * c, [0.1; 10]] / (c * d), ... % mu then s
                  'absbounds', [[-Inf; Inf], [1e-6 / (c * d); Inf]]);

loggaussian = struct('ss', 0.1 * d .^ 2, 'ms', 1.5, 'Kmu', 0.2, 'mumu', log(c), ...
                     'bounds', [[-4; 4] + log(c), [0.1; 10] / d .^ 2], ... % mu then s
                     'absbounds', [[-Inf; Inf], [1e-6; Inf]]);

student = struct('mumu', 150 * c, 'smu', (1 / 20) .^ 2 / (c * d) ^ 2, ...
                 'am', 0.1, 'bm', 1, ...
                 'mlow', 1e-3, 'mhigh', 100, ...
                 'ms', 1.1, 'ss', (1 / 10) .^ 2 / (c * d) ^ 2, ...
                 'bounds', [[10; 300] * c, [0.1; 100], [0.001; 10] * (c * d) ^ 2], ... % mu then m then s
                 'absbounds', [[-Inf; Inf], [1e-6; Inf], [1e-6; Inf] * (c * d) ^ 2]);

logstudent = struct('mumu', log(c), 'smu', 1, ...
                    'am', 0.2, 'bm', 4, ...
                    'mlow', 1e-3, 'mhigh', 100, ...
                    'ms', 1.1, 'ss', 4 / d .^ 2, ...
                    'bounds', [[-2; 2] + log(c), [0.1; 100], [0.1; 10] * d .^ 2], ... % mu then m then s
                    'absbounds', [[-Inf; Inf], [1e-6; Inf], [1e-6; Inf] * d .^ 2]);

% Note that "gamma" is being redefined here,
%  so we cannot use the function gamma in this function (not that we'd want to).
gamma = struct('am', 0.08 * d ^ 2, 'bm', 1.5, ...
               'mlow', 1e-3, 'mhigh', 300, ...
               'ms', 1, 'ss', 2 * c, ...
               'bounds', [[0.1; 100], [10; 60] / c], ... % m then s
               'absbounds', [[1e-6; Inf], [1e-6; Inf] / c]);

invgamma = struct('am', 0.08 * d ^ 2, 'bm', 1.5, ...
                  'mlow', 1e-3, 'mhigh', 300, ...
                  'ms', 1, 'ss', 0.5 / c, ...
                  'bounds', [[0.1; 100], [0.003; 0.5] * c], ... % m then s
                  'absbounds', [[1e-6; Inf], [1e-6; Inf] * c]);

skewstudent = struct('muxi', 150 * c, 'sxi', (1 / 20) .^ 2 / (c * d) ^ 2, ...
                     'am', 0.1, 'bm', 1, ...
                     'mlow', 0.5 + 1e-6, 'mhigh', 150, ...
                     'ms', 2, 'ss', (1 / 10) .^ 2 / (c * d) ^ 2, ...
                     'snu', 1 / d ^ 2, ...
                     'bounds', [[10 ; 300] * c, [0.6; 100], [25; 250] * (c * d) ^ 2, [-15; 15] * d], ... % xi then m then s then nucr
                     'absbounds', [[-Inf; Inf], [0.5 + 1e-6; Inf], [1e-6; Inf] * (c * d) ^ 2, [-Inf; Inf]]);

logskewstudent = struct('muxi', log(c), 'sxi', 1, ...
                        'am', 0.2, 'bm', 4, ...
                        'mlow', 0.5 + 1e-6, 'mhigh', 150, ...
                        'ms', 1.1, 'ss', 4 / d .^ 2, ...
                        'snu', 0.1 / d ^ 2, ...
                        'bounds', [[-2; 2] + log(c), [0.6; 100], [0.1; 10] * d ^ 2, [-30; 30] * d], ... % xi then m then s then nucr
                        'absbounds', [[-Inf; Inf], [0.5 + 1e-6; Inf], [1e-6; Inf] * d .^ 2, [-Inf; Inf]]);

gammapow = struct('am', 0.08 * d ^ 2, 'bm', 1.5, ...
                  'mlow', 1e-3, 'mhigh', 300, ...
                  'ms', 1, 'ss', 1 * c, ...
                  'Nk', 2, 'ak', 1, 'bk', [0.2; 0.2], 'ck', [0.5; 0.5], ...
                  'klow', 0.1, 'khigh', 10, ...
                  'bounds', [[0.1; 100], [10; 60] / c, [0.4; 2.5]], ... % m then s then k
                  'absbounds', [[1e-6; Inf], [1e-6; Inf] / c, [0.1; 10]]);

% Do the precalcs for the proGamma distributions.
student = progammaprecalcs(student);
logstudent = progammaprecalcs(logstudent);
gamma = progammaprecalcs(gamma);
invgamma = progammaprecalcs(invgamma);
skewstudent = progammaprecalcs(skewstudent);
logskewstudent = progammaprecalcs(logskewstudent);
gammapow = progammaprecalcs(gammapow);
gammapow = propowprecalcs(gammapow);

priors = struct('typelist', {typelist}, ...
                'posonly', posonly, ...
                'Etypes', Etypes, ...
                'Efns', {Efns}, ...
                'Gaussian', gaussian, ...
                'logGaussian', loggaussian, ...
                'Student', student, ...
                'logStudent', logstudent, ...
                'Gamma', gamma, ...
                'invGamma', invgamma, ...
                'skewStudent', skewstudent, ...
                'logskewStudent', logskewstudent, ...
                'Gammapow', gammapow, ...
                'xrange', xrange, ...
                'Ngridmax', {Ngridmax}, ...
                'shrinklimit', shrinklimit, ...
                'groupstep', groupstep, ...
                'familyoffsets', familyoffsets, ...
                'Nparams', Nparams, ...
                'ditherfns', {ditherfns});

return;


% Local Variables: 
% indent-line-function: indent-relative
% eval: (auto-fill-mode 0)
% End:
```
