## Supplementary material for "Elevated cerebrospinal fluid cytokine levels in tuberculous meningitis predict survival in response to dexamethasone": Software: TAGS

ars.m,947
function y 1,0
% bounds contains lbound,34,1467
%  tangents fitted to the log density,35,1528
%  tangents fitted to the log density, and ubound,35,1528
vals 37,1601
m 38,1622
vals 39,1637
grads 40,1654
% RFS: There is a serious risk that this calculation will end up giving NaNs,42,1681
bounds 45,1910
% RFS: The following is to make sure that bounds 47,2069
boundedhigh 56,2617
   vals 64,2861
   maxvals 65,2889
   m 66,2913
   vals 67,2933
   grads 68,2959
   bounds 70,2989
   boundedlow 72,3151
   boundedhigh 73,3205
    Nrejects 88,3635
    done 89,3653
       k 92,3747
    u 106,4218
    z1 108,4233
    maxvals 109,4255
    m 110,4286
    vals 111,4307
    z1 112,4334
    l 114,4358
        loc 124,4583
    points 126,4613
    vals 132,4723
    grads 133,4770
    bounds 136,4867
    %     nlogf 142,5213
    %     logf 143,5249
    maxvals 145,5274
    m 146,5299
    vals 147,5320
y 149,5351

calccompprobs.m,454
function compprob 1,0
% Usage: compprob 2,44
% For each cell,4,144
% HOWEVER,9,448
% HOWEVER, where there are equal values,9,448
%  less than the second. The easiest way of fixing this,10,518
%  less than the second. The easiest way of fixing this, though it is rather inefficient,10,518
%  is to call it twice giving the cells both ways round,11,608
mytitl 18,829
x 32,1116
Px 33,1135
end37,1328
Px 40,1366
cumPx 41,1383
compprob 43,1407

checkdither.m,268
mytitl 13,458
Ncyks 22,677
Ntcounts 23,703
Nhivs 24,738
         cold 31,918
         cnew 32,959
         cmin 34,1001
         cmax 35,1036
         Nbins 37,1072
         cc 38,1093
         hold 40,1152
         hnew 41,1184
         keyboard;47,1402

comparegroups.m,1281
%  distribution types given in priors,10,706
% xs should be a cell array of size 12,886
%  each cell of which contains a vector of values of x;13,937
%  distributions will have their mean numerically measured with linear weighting;17,1154
%  distributions will have their arithmetic mean numerically measured with a logarithmic grid;20,1340
%  log(21,1435
Ngroups 86,4193
Ntypes 96,4365
parmmaxes 98,4400
xmeanvalslin 99,4435
xmeanvalslog 100,4473
xmeanvalsgeom 101,4511
Pxmeanvals 102,4550
compprobslin 130,5429
compprobslog 131,5461
compprobsgeom 132,5493
      fprintf('Starting ngroup %d, test distribution type %d\n', ngroup, ntype)149,6077
         fprintf('Running gridassess on ngroup %d, ntype %d\n', ngroup, ntype)153,6251
                            nfig, xmeangridlin, xmeangridlog)157,6529
         parmmaxes{parmmaxes160,6637
         Pxmeanvals{Pxmeanvals161,6688
         xmeanvalslin{xmeanvalslin162,6741
         xmeanvalslog{xmeanvalslog163,6798
         xmeanvalsgeom{xmeanvalsgeom164,6855
Etypes 176,7098
Etypes 177,7129
Ptypes 178,7164
Ptypes 179,7187
allcompprobslin 225,9204
allcompprobslog 226,9236
allcompprobsgeom 227,9268
Ptypesreshaped 229,9303
compprobslin 230,9353
compprobslog 233,9555
compprobsgeom 236,9757

ditherdo.m,1336
function 1,0
% Usage:Usage2,83
% Inputs: cyks is a [Npatients, Ncyks] array of cytokine values,3,166
%  which have not been previously dithered,4,231
% This function dithers the cytokine values that are so jammed,6,349
%  doing so over a wide range,7,413
%  savesetids{1 + hiv} and ncyk,10,608
%     1.89         12:jun:20    rfs      Fixed new NaNs appearing;30,1833
mytitl 34,2089
Nncyks 43,2305
Nhivs 44,2329
Ntcounts 45,2358
absmin 47,2394
Ntrieslong 48,2410
Ntriesshort 49,2428
Ntypes 51,2448
groupstep 52,2482
familyoffsets 53,2512
Nparams 54,2550
      thismax 62,2736
   thismin 64,2764
            nfig 80,3327
            firstfig 82,3394
            ch 90,3670
               ch 96,3907
         Etypes 109,4342
         Etypes 110,4377
         Etypes 111,4417
         lowtop 113,4462
         highbot 114,4496
         ind 116,4536
         cold 120,4665
         cnew 121,4698
         fprintf('ncyk = %d, thistcount = %d, thishiv = %d:\n', ncyk, thistcount, thishiv)123,4747
            oldc 126,4903
            elseif thisc 130,5065
               thisc 133,5211
               keyboard;144,5734
function out 158,5920
% Usage: out 159,5994
ntype 166,6343
ditherfns 168,6383
parms 170,6414
out 172,6450
out = feval(ditherfns{ntype}, low, high, Ntries, parms{parms172,6450

dithergam.m,425
function out 1,0
% Usage: out 2,51
% Draws a sample from the Gamma distribution with the given parameters,3,102
% Does this by the dumbest least efficient way available to save programming time,5,223
mytitl 12,451
ntried 21,668
r 22,680
fprintf('low = %g, high = %g, m = %g, s = %g:\n', low, high, m, s)23,691
found 24,759
   ntried 33,899
   fprintf('\rTried %d times ...         ', ntried)34,923
   out 40,1110

dithergau.m,420
function out 1,0
% Usage: out 2,52
% Draws a sample from the Gaussian distribution with the given parameters,3,104
% Does this by the dumbest least efficient way available to save programming time,5,228
mytitl 12,456
ntried 21,673
fprintf('low = %g, high = %g, mu = %g, s = %g:\n', low, high, mu, s)22,685
found 23,755
   ntried 32,897
   fprintf('\rTried %d times ...         ', ntried)33,921
   out 39,1108

dithergp.m,358
function out 1,0
% Usage: out 2,53
% Does this by the dumbest least efficient way available to save programming time,5,230
mytitl 12,458
ntried 21,674
r 22,686
fprintf('low = %g, high = %g, m = %g, s = %g, k = %g:\n', low, high, m, s, k)23,697
found 24,776
   ntried 33,928
   fprintf('\rTried %d times ...         ', ntried)34,952
   out 40,1139

ditherig.m,432
function out 1,0
% Usage: out 2,50
% Draws a sample from the inverseGamma distribution with the given parameters,3,100
% Does this by the dumbest least efficient way available to save programming time,5,228
mytitl 12,456
ntried 21,672
r 22,684
fprintf('low = %g, high = %g, m = %g, s = %g:\n', low, high, m, s)23,695
found 24,763
   ntried 33,908
   fprintf('\rTried %d times ...         ', ntried)34,932
   out 40,1119

ditherlg.m,423
function out 1,0
% Usage: out 2,51
% Draws a sample from the logGaussian distribution with the given parameters,3,102
% Does this by the dumbest least efficient way available to save programming time,5,229
mytitl 12,457
ntried 21,673
fprintf('low = %g, high = %g, mu = %g, s = %g:\n', low, high, mu, s)22,685
found 23,755
   ntried 32,902
   fprintf('\rTried %d times ...         ', ntried)33,926
   out 39,1113

ditherls.m,443
function out 1,0
% Usage: out 2,54
% Draws a sample from the logStudent distribution with the given parameters,3,108
% Does this by the dumbest least efficient way available to save programming time,5,234
mytitl 12,462
ntried 21,678
r 22,690
fprintf('low = %g, high = %g, mu = %g, s = %g, m = %g:\n', low, high, mu, s, m)23,701
found 24,782
   ntried 33,953
   fprintf('\rTried %d times ...         ', ntried)34,977
   out 40,1164

ditherlss.m,401
function out 1,0
% Usage: out 2,61
% Does this by the dumbest least efficient way available to save programming time,5,252
mytitl 12,480
ntried 21,697
nu 22,709
r 23,729
mu 24,740
fprintf('low = %g, high = %g, xi = %g, m = %g, s = %g, nucr = %g:\n', low, high, xi, m, s, nucr)25,801
found 26,899
   ntried 35,1077
   fprintf('\rTried %d times ...         ', ntried)36,1101
   out 42,1288

ditherss.m,485
function out 1,0
% Usage: out 2,60
% Draws a sample from the skewStudent distribution with the given parameters,3,120
% Does this by the dumbest least efficient way available to save programming time,5,247
mytitl 12,475
ntried 21,691
nu 22,703
r 23,723
mu 24,734
fprintf('low = %g, high = %g, xi = %g, m = %g, s = %g, nucr = %g:\n', low, high, xi, m, s, nucr)25,795
found 26,893
   ntried 35,1066
   fprintf('\rTried %d times ...         ', ntried)36,1090
   out 42,1277

ditherstud.m,440
function out 1,0
% Usage: out 2,56
% Draws a sample from the Student distribution with the given parameters,3,112
% Does this by the dumbest least efficient way available to save programming time,5,235
mytitl 12,463
ntried 21,681
r 22,693
fprintf('low = %g, high = %g, mu = %g, s = %g, m = %g:\n', low, high, mu, s, m)23,704
found 24,785
   ntried 33,951
   fprintf('\rTried %d times ...         ', ntried)34,975
   out 40,1162

Eadd.m,118
function 1,0
% Usage:Usage2,29
%  y 4,85
sz1 21,524
sz2 22,540
y 32,727
ind1 34,744
ind2 36,822
ind3 38,901

Egamma.m,620
function E 1,0
% Usage: E 2,47
%  in the array x 4,168
%  in the array x = parmgrid{3} at the parameters m 4,168
%  in the array x = parmgrid{3} at the parameters m = parmgrid{1}, s 4,168
%  in the array x = parmgrid{3} at the parameters m = parmgrid{1}, s = parmgrid{2}, r 4,168
   warned 37,875
Nparams 40,895
m 46,1007
s 47,1024
x 48,1041
r 49,1058
Elike 51,1071
   ms 55,1183
   ss 56,1201
   rs 57,1219
   am 58,1237
   bm 59,1255
   Km 60,1273
   sz 62,1292
   szwanted 63,1309
   m 64,1334
   s 65,1383
   Epriors 67,1433
   Epriorm 68,1505
   Eprior 70,1586
   E 72,1618
   E 74,1663

Egammapow.m,690
function E 1,0
% Usage: E 2,50
%  in the array x 4,174
%  in the array x = parmgrid{3} at the parameters m 4,174
%  s 5,242
%  s = parmgrid{2}, k 5,242
%  s = parmgrid{2}, k = parmgrid{3}, r 5,242
mytitl 15,626
   warned 40,1077
Nparams 43,1097
m 49,1209
s 50,1226
k 51,1243
x 52,1260
r 53,1277
Elike 55,1295
   ms 59,1428
   ss 60,1446
   rs 61,1464
   am 62,1482
   bm 63,1500
   Km 64,1518
   Nk 65,1536
   ak 66,1554
   bk 67,1572
   ck 68,1590
   Kk 69,1608
   sz 71,1627
   szwanted 72,1664
   sz 73,1695
   m 74,1724
   s 75,1773
   k 76,1822
   Epriors 78,1872
   Epriorm 79,1944
   Epriork 80,2024
   Eprior 85,2163
   E 87,2205
   E 89,2250

Egauss.m,609
function E 1,0
% Usage: E 2,47
%  in the array x 4,168
%  in the array x = parmgrid{3} at the parameters mu 4,168
%  in the array x = parmgrid{3} at the parameters mu = parmgrid{1,} s parmgrid{parmgrid4,168
%  each of which is an array of the same dimension,5,251
%     1.1          01:may:20    rfs      First version,10,361
mytitl 14,588
   warned 39,1033
Nparams 42,1053
mu 48,1165
s 49,1183
x 50,1200
Elike 52,1218
   ms 56,1330
   ss 57,1348
   mumu 58,1366
   Kmu 59,1388
   sz 61,1409
   szwanted 62,1427
   mu 63,1452
   s 64,1503
   Eprior 66,1553
   E 69,1719
   E 71,1764

Einvgamma.m,620
function E 1,0
% Usage: E 2,50
%  in the array x 4,174
%  in the array x = parmgrid{3} at the parameters m 4,174
%  in the array x = parmgrid{3} at the parameters m = parmgrid{1}, s 4,174
%  in the array x = parmgrid{3} at the parameters m = parmgrid{1}, s = parmgrid{2}, r 4,174
   warned 37,887
Nparams 40,907
m 46,1022
s 47,1039
x 48,1056
r 49,1073
Elike 51,1086
   ms 55,1201
   ss 56,1219
   rs 57,1237
   am 58,1255
   bm 59,1273
   Km 60,1291
   sz 62,1310
   szwanted 63,1327
   m 64,1352
   s 65,1401
   Epriors 67,1451
   Epriorm 68,1523
   Eprior 70,1604
   E 72,1636
   E 74,1681

Eloggauss.m,620
function E 1,0
% Usage: E 2,50
%  in the array x 4,174
%  in the array x = parmgrid{3} at the parameters mu 4,174
%  in the array x = parmgrid{3} at the parameters mu = parmgrid{1,} s parmgrid{parmgrid4,174
%  each of which is an array of the same dimension,5,257
%  in priors,6,336
%  in priors, under a logGaussian distribution,6,336
mytitl 14,611
   warned 39,1062
Nparams 42,1082
mu 48,1197
s 49,1215
x 50,1232
Elike 52,1250
   ms 56,1379
   ss 57,1397
   mumu 58,1415
   Kmu 59,1437
   sz 61,1458
   szwanted 62,1476
   mu 63,1501
   s 64,1552
   Eprior 66,1602
   E 69,1768
   E 71,1813

Elogskewstudent.m,1077
function E 1,0
% Usage: E 2,56
%  in the array x 4,186
%  in the array x = parmgrid{5} at the parameters m 4,186
%  in the array x = parmgrid{5} at the parameters m = parmgrid{3}, s 4,186
%  xi 5,271
%  xi = parmgrid{1}, nu 5,271
%  each of which is an array of the same dimension,6,320
%  where r 7,373
   warned 56,2358
Nparams 59,2378
xi 65,2499
m 66,2517
s 67,2534
nucr 68,2551
x 69,2571
nu 71,2589
   Nx 81,2934
   mu 94,3483
   y 96,3553
   z 97,3573
   w 98,3598
   Eterm1 100,3618
   Eterm2 102,3707
   ind 104,3821
   Eterms 105,3837
   Eterm1neg 108,3921
   Eterm2neg 109,3950
   Eterm2neg 110,3979
   Elike 120,4319
   ms 128,4583
   ss 129,4601
   rs 130,4619
   muxi 131,4636
   sxi 132,4658
   am 133,4678
   bm 134,4696
   Km 135,4714
   snu 136,4732
   sz 138,4753
   szwanted 139,4790
   sz 140,4821
   xi 141,4850
   s 142,4901
   m 143,4950
   nu 144,4999
   nucr 145,5050
   Epriorxi 147,5106
   Epriors 148,5179
   Epriorm 149,5251
   Epriornucr 150,5331
   Eprior 152,5425
   E 154,5481
   E 156,5526

Elogstudent.m,795
function E 1,0
% Usage: E 2,52
%  in the array x 4,178
%  in the array x = parmgrid{4} at the parameters m 4,178
%  in the array x = parmgrid{4} at the parameters m = parmgrid{3}, s 4,178
%  in the array x = parmgrid{4} at the parameters m = parmgrid{3}, s = parmgrid{2}, mu 4,178
%  each of which is an array of the same dimension,5,280
%  where r 6,333
   warned 40,1112
Nparams 43,1132
mu 49,1249
m 50,1267
s 51,1284
x 52,1301
r 54,1319
Elike 56,1332
   ms 61,1527
   ss 62,1545
   rs 63,1563
   mumu 64,1580
   smu 65,1602
   am 66,1622
   bm 67,1640
   Km 68,1658
   sz 70,1677
   szwanted 71,1715
   sz 72,1746
   mu 73,1775
   s 74,1826
   m 75,1875
   Epriormu 77,1925
   Epriors 78,1998
   Epriorm 79,2070
   Eprior 81,2151
   E 83,2194
   E 85,2239

Eskewstudent.m,1160
function E 1,0
% Usage: E 2,53
%  in the array x 4,180
%  in the array x = parmgrid{5} at the parameters m 4,180
%  in the array x = parmgrid{5} at the parameters m = parmgrid{3}, s 4,180
%  xi 5,265
%  xi = parmgrid{1}, nucr 5,265
%  each of which is an array of the same dimension,6,307
%  where r 7,360
%                                        and nu are too tightly coupled;20,1121
   warned 57,2406
Nparams 60,2426
xi 66,2544
m 67,2562
s 68,2579
nucr 69,2596
x 70,2616
nu 72,2634
   Nx 82,2980
   mu 95,3511
   y 97,3581
   z 98,3596
   w 99,3621
   Eterm1 101,3641
   Eterm2 103,3730
   ind 105,3844
   Eterms 106,3860
   Eterm1neg 109,3944
   Eterm2neg 110,3973
   Eterm2neg 111,4002
   Elike 121,4342
   ms 129,4595
   ss 130,4613
   rs 131,4631
   muxi 132,4648
   sxi 133,4670
   am 134,4690
   bm 135,4708
   Km 136,4726
   snu 137,4744
   sz 139,4765
   szwanted 140,4802
   sz 141,4833
   xi 142,4862
   s 143,4913
   m 144,4962
   nu 145,5011
   nucr 146,5062
   Epriorxi 148,5118
   Epriors 149,5191
   Epriorm 150,5263
   Epriornucr 151,5343
   Eprior 153,5437
   E 155,5493
   E 157,5538

Estudent.m,795
function E 1,0
% Usage: E 2,49
%  in the array x 4,172
%  in the array x = parmgrid{4} at the parameters m 4,172
%  in the array x = parmgrid{4} at the parameters m = parmgrid{3}, s 4,172
%  in the array x = parmgrid{4} at the parameters m = parmgrid{3}, s = parmgrid{2}, mu 4,172
%  each of which is an array of the same dimension,5,274
%  where r 6,327
   warned 40,1114
Nparams 43,1134
mu 49,1248
m 50,1266
s 51,1283
x 52,1300
r 54,1318
Elike 56,1331
   ms 61,1509
   ss 62,1527
   rs 63,1545
   mumu 64,1562
   smu 65,1584
   am 66,1604
   bm 67,1622
   Km 68,1640
   sz 70,1659
   szwanted 71,1697
   sz 72,1728
   mu 73,1757
   s 74,1808
   m 75,1857
   Epriormu 77,1907
   Epriors 78,1980
   Epriorm 79,2052
   Eprior 81,2133
   E 83,2176
   E 85,2221

Esub.m,106
function 1,0
% Usage:Usage2,29
%  y 4,85
sz1 23,621
sz2 24,637
y 38,960
ind1 40,982
ind3 42,1060

Esum.m,138
function 1,0
% Usage:Usage2,27
%  y 4,81
shift 21,503
ind 22,526
sz 25,584
sz 26,598
repper 27,631
x 30,676
y 32,708
y 34,738

gridassess.m,1104
% This function takes some data x,5,362
%  maximum number of grid points to be used priors.Ngridmax,Ngridmax6,398
%  the index number of the distribution type to be assessed as defined by setpriors.m,m7,459
%   corresponding to parameter nparam,14,928
%   output corresponding to x,15,999
%   of same size as parmgrid{1}, except that the last dimension is one longer,18,1177
%   of same size as parmgrid{1}, except that the last dimension is one longer, with Eprior being last,18,1177
%  note that xmeanvalslog does not contain logarithms,38,2680
% If xmeangridlog is passed as above,40,2819
%     1.5          05:may:20    rfs      Also plots fit versus true versus histogram,48,3167
%     1.8          06:may:20    rfs      Added computations of means,53,3606
%     1.31         11:may:20    rfs      Now only allows one Ngrid value to advance per round,58,4069
Ngridmax 100,5594
shrinklimit 101,5629
typename 102,5670
prior 103,5705
bounds 104,5741
absbounds 105,5764
Nparams 111,5845
      parmstruevals 125,6404
         parmstruevals 128,6531
Ngrid 138,6910
niter 144,7051

ksample.m,83
function y 1,0
logf 18,481
gradlogf 19,531
p_points 24,662
k 33,900
y 35,917

loghypergeom1F1v2.m,429
function out 1,0
% Usage: out 2,51
debugging 25,992
w 37,1446
discrim 38,1471
sdiscrim 40,1511
high 41,1542
low 42,1569
ind 44,1596
% The potential biggest terms are now roughly at positions high and zero in the series,51,1765
n1 54,1903
n3 55,1928
Eterm1 56,1953
Eterm3 57,1980
indforterm1 80,2860
% Having identified the positions and values of the biggest terms in the series to be summed,93,3322
out 95,3453

log_k_density.m,137
function y 1,0
b 27,930
c 28,940
sz 30,951
k 31,965
krep 34,993
b 35,1019
c 36,1042
d 38,1066
dd 40,1088
y 43,1121
y 45,1190

log_k_derivative.m,130
function y 1,0
b 22,686
c 23,696
sz 25,707
k 26,721
kN 30,750
b 31,774
c 32,797
dd 34,821
ee 36,861
y 39,894
y 41,940

log_m_density.m,26
function y 1,0
y 21,611

log_m_derivative.m,26
function y 1,0
y 19,534

makesynthdata.m,4214
function 1,0
% Usage:Usage2,60
% parmstrue is a cell array of size 4,144
%  element of Nx;11,628
%     1.18         08:may:20    rfs      ntype made a parameter to enable forcing it;27,1505
mytitl 36,2118
Ntypes 49,2374
sz 51,2409
Nx 52,2424
Ngroups 53,2436
type 59,2535
parmstrue 61,2567
         ss 69,2705
         ms 70,2729
         rs 71,2753
         Kmu 72,2776
         mumu 73,2802
         strue 75,2834
         mutrue 76,2879
         x{x78,2938
         parmstrueGaussian 80,3006
         parmstrue{parmstrue82,3117
         prior 86,3277
         ss 87,3314
         ms 88,3338
         rs 89,3362
         Kmu 90,3385
         mumu 91,3411
         strue 93,3443
         mutrue 94,3488
         x{x96,3550
         parmstruelogGaussian 98,3626
         parmstrue{parmstrue100,3743
         prior 104,3905
         mumu 105,3938
         smu 106,3966
         am 107,3992
         bm 108,4016
         mlow 109,4040
         mhigh 110,4068
         ms 111,4098
         ss 112,4122
         mutrue 114,4150
         mtrue 115,4201
         mtrue 116,4236
         strue 117,4283
         rtrue 118,4336
         x{x120,4372
         parmstrueStudent 122,4480
         parmstrue{parmstrue126,4680
         prior 130,4823
         mumu 131,4859
         smu 132,4887
         am 133,4913
         bm 134,4937
         mlow 135,4961
         mhigh 136,4989
         ms 137,5019
         ss 138,5043
         mutrue 140,5071
         mtrue 141,5122
         mtrue 142,5157
         strue 143,5204
         rtrue 144,5257
         x{x146,5293
         parmstruelogStudent 148,5406
         parmstrue{parmstrue152,5609
         prior 156,5753
         am 157,5784
         bm 158,5808
         mlow 159,5832
         mhigh 160,5860
         Km 161,5890
         ms 162,5914
         ss 163,5938
         rs 164,5962
         mtrue 166,5990
         mtrue 167,6025
         strue 168,6072
         rtrue 169,6118
         x{x171,6154
         parmstrueGamma 173,6223
         parmstrue{parmstrue176,6362
         prior 180,6507
         am 181,6541
         bm 182,6565
         mlow 183,6589
         mhigh 184,6617
         Km 185,6647
         ms 186,6671
         ss 187,6695
         rs 188,6719
         mtrue 190,6747
         mtrue 191,6782
         strue 192,6829
         rtrue 193,6875
         x{x195,6911
         parmstrueinvGamma 197,6985
         parmstrue{parmstrue200,7130
         prior 204,7284
         muxi 205,7321
         sxi 206,7349
         am 207,7375
         bm 208,7399
         mlow 209,7423
         mhigh 210,7451
         ms 211,7481
         ss 212,7505
         snu 213,7529
         xitrue 215,7559
         mtrue 216,7610
         mtrue 217,7645
         strue 218,7692
         rtrue 219,7745
         nutrue 220,7777
         nucrtrue 221,7821
         mutrue 222,7861
         x{x224,7961
         parmstrueskewStudent 226,8080
         parmstrue{parmstrue231,8357
         prior 235,8520
         muxi 236,8560
         sxi 237,8588
         am 238,8614
         bm 239,8638
         mlow 240,8662
         mhigh 241,8690
         ms 242,8720
         ss 243,8744
         snu 244,8768
         xitrue 246,8798
         mtrue 247,8849
         mtrue 248,8884
         strue 249,8931
         rtrue 250,8984
         nutrue 251,9016
         nucrtrue 252,9060
         mutrue 253,9100
         x{x255,9200
         parmstruelogskewStudent 257,9324
         parmstrue{parmstrue262,9613
         prior 266,9776
         am 267,9810
         bm 268,9834
         mlow 269,9858
         mhigh 270,9886
         Km 271,9916
         ms 272,9940
         ss 273,9964
         rs 274,9988
         Kk 275,10012
         Nk 276,10036
         ak 277,10060
         bk 278,10084
         ck 279,10108
         mtrue 281,10133
         mtrue 282,10168
         strue 283,10215
         ktrue 284,10261
         rtrue 285,10303
         x{x287,10344
         parmstrueGammapow 289,10429
         parmstrue{parmstrue293,10630
      otherwise,296,10758
   ind 300,10834
parmstrue 315,11284

m_sample.m,314
function y 1,0
% Usage: y 2,33
% Original Usage: y 3,66
r 27,867
k 28,874
x 29,881
    y1 34,991
    N 36,1023
    z1 38,1054
    X1 40,1094
    z2 42,1120
    X2 44,1175
    z3 46,1201
    X3 48,1257
    b 51,1288
    a 52,1305
    y 53,1342
           logf 61,1610
           gradlogf 62,1663

nutransderiv.m,118
function nuderiv 1,0
% Usage: nuderiv 2,39
% If you change these,20,599
alpha 22,705
beta 23,718
nuderiv 25,732

nutransinv.m,492
function nucr 1,0
% Usage: nucr 2,32
% This function applies the transformation to nu inverse to that in nutrans.m;m3,64
% If you change these,22,668
alpha 24,774
beta 25,787
% Solve the equation x ^ 3 - y * x ^ 2 + alpha * x - beta * y 27,801
q 31,916
r 32,927
% The equation is now x ^ 3 + p * x ^ 2 + q * x + r 34,945
b 38,1069
A 40,1109
B 41,1161
nucr 43,1214
function y 48,1247
% Usage: y 49,1269
% Calculates the real cube root of the real number x,50,1291
y 53,1405

nutrans.m,103
function nu 1,0
% Usage: nu 2,29
% If you change these,20,510
alpha 22,621
beta 23,634
nu 25,648

piecewise_exponential_density.m,297
function y 1,0
% Usage: y 2,77
%  at the points xx,4,231
%   bounds gives the changeover points from each piece to the next;5,259
%   points is a list of x values one in each piece;6,327
%   vals are the unnormalised log-densities at points;7,379
mytitl 15,688
Npieces 24,925
vals 26,956

piecewise_exponential.m,239
function y 1,0
maxvals 23,854
vals 31,1066
shifted_bounds 33,1090
bounds 34,1122
index 35,1148
weights 36,1169
zindex 38,1377
z 44,1583
u 45,1628
bz 47,1639
sbz 48,1655
   y 57,1971
   y 60,2128
    a 64,2226
    y 65,2283

plotpriorsamples.m,593
% If Nsamples is negative,4,227
% If Nsamples is negative, it instead takes one sample from the prior,4,227
%  and draws a random sample of size abs(5,298
Ntypes 59,1660
   linestyles 64,1790
   Nsamples 69,1896
   Nhistsamples 71,1918
Nx 74,1944
ntypes 83,2126
    histsamples 89,2275
    theseparms 92,2352
    theseparms 93,2427
    params 94,2469
    [params{:}] = ndgrid(theseparms{:}, xx)95,2515
   hold on;103,2731
   na 111,2979
   h 112,2992
   hold off;116,3111
      xxhist 136,3647
   clf;139,3723
   hold on;140,3731
   deltax 147,3847
   hold off;154,4114

plotpriors.m,2711
prior 26,862
ss 27,887
ms 28,902
Kmu 29,917
mumu 30,934
Nplot 32,954
muax 34,968
sax 35,1034
Egaussconj 38,1137
Pgaussconj 41,1320
nfig 43,1353
nfig 45,1364
clf;47,1395
colorbar;51,1463
prior 59,1632
ss 60,1660
ms 61,1675
Kmu 62,1690
mumu 63,1707
Nplot 65,1727
muax 67,1741
sax 68,1807
Eloggaussconj 71,1910
Ploggaussconj 74,2096
nfig 76,2135
clf;78,2166
colorbar;82,2237
prior 90,2406
mlow 94,2443
mhigh 95,2462
deltam 96,2483
mm 97,2499
bm 99,2546
Km 100,2561
Emm 101,2576
Pmm 102,2653
nfig 104,2672
ms 113,2806
ss 114,2821
s 115,2836
Ps 117,2946
nfig 119,2962
mumu 128,3095
smu 129,3114
xxlow 130,3131
xxhigh 131,3161
deltax 132,3192
xx 133,3225
Pxx 135,3325
nfig 137,3344
prior 146,3523
mlow 150,3563
mhigh 151,3582
deltam 152,3603
mm 153,3619
bm 155,3666
Km 156,3681
Emm 157,3696
Pmm 158,3773
nfig 160,3792
ms 169,3929
ss 170,3944
s 171,3959
Ps 173,4071
nfig 175,4087
mumu 184,4223
smu 185,4242
xxlow 186,4259
xxhigh 187,4289
deltax 188,4320
xx 189,4353
Pxx 191,4453
nfig 193,4472
prior 202,4649
mlow 206,4684
mhigh 207,4703
deltam 208,4724
mm 209,4740
bm 211,4787
Km 212,4802
Emm 213,4817
Pmm 214,4894
nfig 216,4913
ms 225,5045
ss 226,5060
s 227,5075
Ps 229,5187
nfig 231,5203
prior 240,5373
mlow 244,5411
mhigh 245,5430
deltam 246,5451
mm 247,5467
bm 249,5514
Km 250,5529
Emm 251,5544
Pmm 252,5621
nfig 254,5640
ms 263,5775
ss 264,5790
s 265,5805
Ps 267,5915
nfig 269,5931
prior 278,6107
mlow 282,6148
mhigh 283,6167
deltam 284,6188
mm 285,6204
bm 287,6251
Km 288,6266
Emm 289,6281
Pmm 290,6358
nfig 292,6377
ms 301,6515
ss 302,6530
s 303,6545
Ps 305,6655
nfig 307,6671
muxi 316,6808
sxi 317,6827
xxlow 318,6844
xxhigh 319,6874
deltax 320,6905
xx 321,6938
Pxx 323,7038
nfig 325,7057
snu 333,7198
xxlow 334,7215
xxhigh 335,7240
deltax 336,7264
xx 337,7297
Pxx 339,7388
nfig 341,7407
prior 350,7594
mlow 354,7638
mhigh 355,7657
deltam 356,7678
mm 357,7694
bm 359,7741
Km 360,7756
Emm 361,7771
Pmm 362,7848
nfig 364,7867
ms 373,8008
ss 374,8023
s 375,8038
Ps 377,8148
nfig 379,8164
muxi 388,8304
sxi 389,8323
xxlow 390,8340
xxhigh 391,8370
deltax 392,8401
xx 393,8434
Pxx 395,8534
nfig 397,8553
snu 405,8697
xxlow 406,8714
xxhigh 407,8739
deltax 408,8763
xx 409,8796
Pxx 411,8887
nfig 413,8906
prior 422,9095
mlow 426,9133
mhigh 427,9152
deltam 428,9173
mm 429,9189
bm 431,9236
Km 432,9251
Emm 433,9266
Pmm 434,9343
nfig 436,9362
ms 445,9497
ss 446,9512
s 447,9527
Ps 449,9639
nfig 451,9655
Nk 460,9788
ak 461,9803
bk 462,9818
ck 463,9833
Kk 464,9848
klow 465,9863
khigh 466,9882
k 467,9903
Pk 472,10046
nfig 474,10062

progammaprecalcs.m,117
function priorstruct 1,0
% Usage: priorstruct 2,54
deltam 21,536
mm 22,552
Pmm 24,721
priorstruct.Km Km25,739

propowprecalcs.m,117
function priorstruct 1,0
% Usage: priorstruct 2,52
deltak 21,531
kk 22,548
Pkk 27,784
priorstruct.Kk Kk28,802

randdiscreteE.m,391
function 1,0
% Usage:Usage2,51
% The probability of each outcome is given by exp(-Es)Es5,198
%  outcomes are 8,378
%  outcomes are 1, 2, ..., M,8,378
%  outcomes are 1, 2, ..., M, and Es(8,378
M 33,1112
N 34,1129
ind 40,1226
   sz 44,1282
Es 58,1606
minEs 61,1646
Es 62,1670
probs 63,1731
probsum 64,1750
ind 65,1775
probs 72,1945
probs 75,2011
out 77,2038
out 79,2109

runrealdata.m,1562
%  each cytokine number in the vector ncyks,6,381
%  use in each group those with a 1 in the logical array selection,7,439
%  of T alleles for that patient; each entry is between 0 and 14,828
%  alternatively it allocates patients to groups to be compared,15,893
% If allcompprobslin,19,1104
% If allcompprobslin, allcompprobslog,19,1104
% If allcompprobslin, allcompprobslog, allcompprobsgeom,19,1104
%  they should be from another previous run with fewer types,20,1190
%     1.1          15:may:20    rfs      First version,25,1333
%     1.60         16:may:20    rfs      Typo leading to massive memory usage fixed;28,1539
%     1.72         26:may:20    rfs      Now takes in previously calculated values;33,1952
%     1.85         03:jun:20    rfs      Added geometric means,35,2135
%     1.86         04:jun:20    rfs      Bugs in above fixed,36,2217
mytitl 38,2303
cyks 74,2895
tcount 75,2922
Npatients 77,2951
Ngroups 78,2978
Nncyks 79,3005
temppriors 80,3029
Ntypes 81,3056
Etypes 83,3095
compprobslin 84,3125
compprobslog 85,3171
compprobsgeom 86,3217
oldNtypes 101,3609
   Etot 104,3696
oldNtypes 107,3760
   allcompprobslin 110,3858
oldNtypes 113,3953
   allcompprobslog 116,4051
oldNtypes 119,4146
   allcompprobsgeom 122,4245
   fprintf('Starting on nncyk %d, ncyk %d.\n', nncyk, ncyk)128,4391
   thesecyks 130,4453
      thesecyks{thesecyks134,4566
   allthesecyks 137,4626
   meancyk 138,4666
   priors 140,4710
   Ngridlin 142,4743
   xmeangridlin 143,4762
   xmeangridlog 146,4854
diary off;183,6157

setpriors.m,877
function priors 1,0
% Usage: priors 2,46
typelist 68,3283
Ntypes 70,3430
posonly 71,3457
ditherfns 74,3631
shrinklimit 76,3766
Nparams 77,3807
Nsubfigures 94,4446
groupstep 98,4561
familyoffsets 99,4591
familyoffsets 100,4628
%  a and b will in the code below be denoted am and bm;112,5316
%  to ensure numerical integrability,113,5372
%  a and b will in the code below be denoted am and bm;126,6170
%  to ensure numerical integrability,127,6226
d 143,7128
xrange 146,7206
gaussian 148,7240
loggaussian 152,7481
student 156,7716
logstudent 163,8140
% Note that "gamma" is being redefined here,170,8529
invgamma 178,8923
skewstudent 184,9212
logskewstudent 192,9750
gammapow 200,10260
logstudent 210,10784
gamma 211,10827
invgamma 212,10860
skewstudent 213,10899
logskewstudent 214,10944
gammapow 215,10995
gammapow 216,11034
priors 218,11072

testids.m,1788
% Constructs Ngroups datasets of each distribution type,7,344
% If allcompprobslin,10,564
% If allcompprobslin, allcompprobslog,10,564
% If allcompprobslin, allcompprobslog, allcompprobsgeom,10,564
% If allcompprobslin, allcompprobslog, allcompprobsgeom, and Etot are passed in,10,564
% If allxs and parmstrues are passed in,12,743
%     1.28         09:may:20    rfs      Rejected any synthetic data with more than 3% edits;21,1188
%                                        fixed bug in setting xmeangridlog;22,1282
priors 90,2980
Ntypes 92,3001
Nmeangrid 94,3036
% Allocate storage,96,3053
% Allocate storage, or expand it,96,3053
Etypes 98,3104
Etypes = NaN(Ntypes, Ntypes); % Etype(98,3104
compprobslog 100,3254
compprobsgeom 101,3300
truelin 102,3347
truelog 103,3388
truegeom 104,3429
   parmstrues 123,4032
oldNtypes 126,4076
   Etot 129,4163
oldNtypes 134,4316
   allcompprobslin 137,4414
oldNtypes 142,4629
   allcompprobslog 145,4727
oldNtypes 150,4942
   allcompprobsgeom 153,5041
oldNtypes 158,5260
   allxs 161,5348
oldNtypes 164,5407
   parmstrues 167,5500
Nxs 170,5569
   fprintf('Starting ntypetrue = %d\n', ntypetrue)178,5704
   catxs 193,6178
   deltax 194,6204
   xmeangridlin 195,6255
      xmeangridlog 202,6517
      nfig 213,6721
   meanlin 236,7731
   meanlog 237,7761
   meangeom 238,7791
      thisparmstrue 244,7951
      thisparmstrue 245,8026
      parmsnd 246,8076
      [parmsnd{:}] = ndgrid(thisparmstrue{:}, xmeangridlin)249,8153
      E 250,8214
      E = squeeze(feval(priors.Efns{ntypetrue}, parmsnd,250,8214
      E 251,8280
      P 252,8302
      P 253,8321
      [parmsnd{:}] = ndgrid(thisparmstrue{:}, xmeangridlog)259,8484
      E 260,8545
      E 261,8631
      P 262,8653
      P 263,8672

startscript.m,0

restartscript.m,0
