## Supplementary material for "Elevated cerebrospinal fluid cytokine levels in tuberculous meningitis predict survival in response to dexamethasone": Software: testids.m

```
function testids(seed, savesetid, savedir, Ngroups, ...
                 allcompprobslin, allcompprobslog, allcompprobsgeom, ...
                 Etot, allxs, parmstrues);
% Usage: testids(seed, savesetid, savedir, Ngroups, ...
%                allcompprobslin, allcompprobslog, allcompprobsgeom, ...
%                Etot, allxs, parmstrues);
% Constructs Ngroups datasets of each distribution type, then checks
%  (a) whether each group's distribution type is correctly identified; and
%  (b) whether the ordering of means in each group is correctly identified.
% If allcompprobslin, allcompprobslog, allcompprobsgeom, and Etot are passed in, then they are 
%  assumed to contain previously calculated values that do not need recalculating.
% If allxs and parmstrues are passed in, then they are used instead of drawing
%  the test sets from random generators.

% Change Log:
%
%     1.1          08:may:20    rfs      First version.
%     1.22         08:may:20    rfs      Version number harmonisation.
%     1.25         09:may:20    rfs      Added temporary saving after each true type.
%     1.27         09:may:20    rfs      Rejected any synthetic data with more than 10% edits.
%     1.28         09:may:20    rfs      Rejected any synthetic data with more than 3% edits;
%                                        fixed bug in setting xmeangridlog; restricted xmeangridlin to positive.
%     1.29         09:may:20    rfs      Fixed bug calculating truelin and truelog.
%     1.31         11:may:20    rfs      Moved setting Ngridmax and shrinklimit to setpriors.m .
%     1.39         12:may:20    rfs      Removed suppression of truelog for non-posonly true distributions.
%     1.53         14:may:20    rfs      Reduced size of group 2 slightly to try to not exceed memory limits.
%     1.66         21:may:20    rfs      Made Ngroups arbitrary.
%     1.67         21:may:20    rfs      Saves Etot and allcompprobslin/log also.
%     1.71         26:may:20    rfs      Allows picking up values already calculated.
%     1.85         03:jun:20    rfs      Added geometric means, not yet debugged.

mytitl = ' /home/rfs/ramakrishnan/software/cytokines/rfs/SCCS/s.testids.m 1.102 20/09/03 08:54:05 ';

persistent mytitldone
if isempty(mytitldone),           
   mytitldone = titlfunction(mytitl);
end

global titls

if nargin < 1,
   seed = [];
end
if nargin < 2,
   savesetid = '';
end
if nargin < 3,
   savedir = '';
end
if nargin < 4,
   Ngroups = [];
end
if nargin < 5,
   allcompprobslin = [];
end
if nargin < 6,
   allcompprobslog = [];
end
if nargin < 7,
   allcompprobsgeom = [];
end
if nargin < 8,
   Etot = [];
end
if nargin < 9,
   allxs = [];
end
if nargin < 10,
   parmstrues = [];
end

if isempty(savesetid),
   warning('savesetid is empty');
end
if isempty(savedir),
   savedir = '.';
end
if isempty(Ngroups),
   Ngroups = 2;
end

if ~isempty(seed),
   rng(seed);
end

if ~isempty(savesetid),
   diary(sprintf('%s/matlabout.%s', savedir, savesetid));
end

priors = setpriors;

Ntypes = length(priors.typelist);

Nmeangrid = 50;

% Allocate storage, or expand it, as appropriate.

Etypes = NaN(Ntypes, Ntypes); % Etype(ntype1, ntype2) is E for true ntype2 being thought to be ntype1.
compprobslin = NaN(Ngroups, Ngroups, Ntypes); 
compprobslog = NaN(Ngroups, Ngroups, Ntypes);
compprobsgeom = NaN(Ngroups, Ngroups, Ntypes);
truelin = NaN(Ngroups, Ngroups, Ntypes);
truelog = NaN(Ngroups, Ngroups, Ntypes);
truegeom = NaN(Ngroups, Ngroups, Ntypes);

if isempty(Etot),
   Etot = NaN(Ntypes, Ngroups, Ntypes); % similarly for Etot(ntype1, ngroup, ntype2).
end
if isempty(allcompprobslin),
   allcompprobslin = NaN(Ngroups, Ngroups, Ntypes, Ntypes);
end
if isempty(allcompprobslog),
   allcompprobslog = NaN(Ngroups, Ngroups, Ntypes, Ntypes);
end
if isempty(allcompprobsgeom),
   allcompprobsgeom = NaN(Ngroups, Ngroups, Ntypes, Ntypes);
end
if isempty(allxs) ~= isempty(parmstrues),
   error('If you pass in allxs then you must also pass in parmstrues');
end
if isempty(allxs),
   allxs = cell(Ngroups, Ntypes);
   parmstrues = cell(Ngroups, Ntypes);
end

oldNtypes = size(Etot, 1);
if oldNtypes < Ntypes,
   extraNtypes = Ntypes - oldNtypes;
   Etot = cat(3, cat(1, Etot, ...
                     NaN(extraNtypes, Ngroups, oldNtypes)), ...
              NaN(Ntypes, Ngroups, extraNtypes));
end

oldNtypes = size(allcompprobslin, 3);
if oldNtypes < Ntypes,
   extraNtypes = Ntypes - oldNtypes;
   allcompprobslin = cat(4, cat(3, allcompprobslin, ...
                                NaN(Ngroups, Ngroups, extraNtypes, oldNtypes)), ...
                         NaN(Ngroups, Ngroups, Ntypes, extraNtypes));
end

oldNtypes = size(allcompprobslog, 3);
if oldNtypes < Ntypes,
   extraNtypes = Ntypes - oldNtypes;
   allcompprobslog = cat(4, cat(3, allcompprobslog, ...
                                NaN(Ngroups, Ngroups, extraNtypes, oldNtypes)), ...
                         NaN(Ngroups, Ngroups, Ntypes, extraNtypes));
end

oldNtypes = size(allcompprobsgeom, 3);
if oldNtypes < Ntypes,
   extraNtypes = Ntypes - oldNtypes;
   allcompprobsgeom = cat(4, cat(3, allcompprobsgeom, ...
                                 NaN(Ngroups, Ngroups, extraNtypes, oldNtypes)), ...
                          NaN(Ngroups, Ngroups, Ntypes, extraNtypes));
end

oldNtypes = size(allxs, 2);
if oldNtypes < Ntypes,
   extraNtypes = Ntypes - oldNtypes;
   allxs = cat(2, allxs, cell(Ngroups, extraNtypes));
end

oldNtypes = size(parmstrues, 2);
if oldNtypes < Ntypes,
   extraNtypes = Ntypes - oldNtypes;
   parmstrues = cat(2, parmstrues, cell(Ngroups, extraNtypes));
end

Nxs = 140 + round(rand(Ngroups, 1) * 20);

for ntypetrue = 1 : Ntypes,

   if priors.Etypes(ntypetrue) == Inf,
      continue;
   end

   fprintf('Starting ntypetrue = %d\n', ntypetrue);

   if any(cellfun(@isempty, allxs(:, ntypetrue))),
      % Make the test data, rejecting anything which has more than 3% edited out.
      made = 0;
      while ~made,
         [xs, parmstrue] = makesynthdata(priors, Nxs, ntypetrue);
         if all(cellfun(@length, xs) > 0.97 * Nxs),
            made = 1;
         end
      end % while ~made
      parmstrues(:, ntypetrue) = parmstrue;
   end

   % Make xmeangridlin.
   catxs = cat(1, xs{:});
   deltax = (max(catxs) - min(catxs)) / Nmeangrid;
   xmeangridlin = min(catxs) + [-4 : Nmeangrid + 4].' * deltax;
   xmeangridlin = xmeangridlin(xmeangridlin > 0);

   if priors.posonly(ntypetrue) || min(catxs) > 0,
   
      % Make xmeangridlog.
      deltalx = (log(max(catxs)) - log(min(catxs))) / Nmeangrid;
      xmeangridlog = exp(log(min(catxs)) + [-4 : Nmeangrid + 4].' * deltalx);

   else
      xmeangridlog = [];
   end

   allxs(:, ntypetrue) = xs(:);

   if isempty(savesetid),
      nfig = [];
   else
      nfig = 1;
   end

   [thisEtypes, thiscompprobslin, thiscompprobslog, thiscompprobsgeom, ...
    thisallcompprobslin, thisallcompprobslog, thisallcompprobsgeom, thisEtot] ...
      = comparegroups(xs, priors, xmeangridlin, xmeangridlog, parmstrue, nfig, ...
                      allcompprobslin(:, :, :, ntypetrue), allcompprobslog(:, :, :, ntypetrue), ...
                      allcompprobsgeom(:, :, :, ntypetrue), Etot(:, :, ntypetrue));

   Etypes(:, ntypetrue) = thisEtypes;
   Etot(:, :, ntypetrue) = thisEtot;
   compprobslin(:, :, ntypetrue) = thiscompprobslin;
   allcompprobslin(:, :, :, ntypetrue) = thisallcompprobslin;
   if ~isempty(thiscompprobslog),
      compprobslog(:, :, ntypetrue) = thiscompprobslog;
      allcompprobslog(:, :, :, ntypetrue) = thisallcompprobslog;
   end
   if ~isempty(thiscompprobsgeom),
      compprobsgeom(:, :, ntypetrue) = thiscompprobsgeom;
      allcompprobsgeom(:, :, :, ntypetrue) = thisallcompprobsgeom;
   end

   % Work out the true comparison state.
   meanlin = NaN(Ngroups, 1);
   meanlog = NaN(Ngroups, 1);
   meangeom = NaN(Ngroups, 1);
   for ngroup = 1 : Ngroups,

      % Calculate the mean at the true parameter values.

      thisparmstrue = parmstrue{ngroup};
      thisparmstrue = getfield(thisparmstrue, priors.typelist{ntypetrue});
      thisparmstrue = struct2cell(thisparmstrue);
      parmsnd = cell(length(thisparmstrue) + 1, 1);

      % Linear version.
      [parmsnd{:}] = ndgrid(thisparmstrue{:}, xmeangridlin);
      E = squeeze(feval(priors.Efns{ntypetrue}, parmsnd, [], 1));
      E = E - min(E);
      P = exp(-E);
      P = P / sum(P);
      meanlin(ngroup) = sum(P .* xmeangridlin);

      %  Previously had next paragraph dependent on
      %  if priors.posonly(ntypetrue),

      [parmsnd{:}] = ndgrid(thisparmstrue{:}, xmeangridlog);
      E = squeeze(feval(priors.Efns{ntypetrue}, parmsnd, [], 1)) - log(xmeangridlog);
      E = E - min(E);
      P = exp(-E);
      P = P / sum(P);
      meanlog(ngroup) = sum(P .* xmeangridlog);
      meangeom(ngroup) = exp(sum(P .* log(xmeangridlog)));

   end % for ngroup = 1 : Ngroups

   for ngroup1 = 1 : Ngroups,
      for ngroup2 = 1 : Ngroups,
         if meanlin(ngroup1) > meanlin(ngroup2),
            truelin(ngroup1, ngroup2, ntypetrue) = 1;
         elseif meanlin(ngroup2) > meanlin(ngroup1),
            truelin(ngroup1, ngroup2, ntypetrue) = 0;
         else
            truelin(ngroup1, ngroup2, ntypetrue) = 0.5;
         end
      end
   end

   for ngroup1 = 1 : Ngroups,
      for ngroup2 = 1 : Ngroups,
         if meanlog(ngroup1) > meanlog(ngroup2),
            truelog(ngroup1, ngroup2, ntypetrue) = 1;
         elseif meanlog(ngroup2) > meanlog(ngroup1),
            truelog(ngroup1, ngroup2, ntypetrue) = 0;
         else
            truelog(ngroup1, ngroup2, ntypetrue) = 0.5;
         end
      end
   end

   for ngroup1 = 1 : Ngroups,
      for ngroup2 = 1 : Ngroups,
         if meangeom(ngroup1) > meangeom(ngroup2),
            truegeom(ngroup1, ngroup2, ntypetrue) = 1;
         elseif meangeom(ngroup2) > meangeom(ngroup1),
            truegeom(ngroup1, ngroup2, ntypetrue) = 0;
         else
            truegeom(ngroup1, ngroup2, ntypetrue) = 0.5;
         end
      end
   end

   if ~isempty(savesetid),
      quicksave(sprintf('%s.%d', savesetid, ntypetrue), savedir);
      close all
   end

   % Put in a temporary save so that we can examine what was going on before all runs finished.
   save tempsave.mat

end % for ntypetrue = 1 : Ntypes

if ~isempty(savesetid),
   save(sprintf('%s/save.%s.mat', savedir, savesetid));
   if isunix,
      unix(sprintf('chmod a-w %s/save.%s.mat', savedir, savesetid));
   end
end

diary off

return;


% Local Variables: 
% indent-line-function: indent-relative
% eval: (auto-fill-mode 0)
% End:
```
